# The RNA phosphatase PIR-1 regulates endogenous small RNA pathways in *C. elegans*

**DOI:** 10.1101/2020.08.03.235143

**Authors:** Daniel A Chaves, Hui Dai, Lichao Li, James J Moresco, Myung Eun Oh, Darryl Conte, John R Yates, Craig C Mello, Weifeng Gu

**Affiliations:** Dept. of Molecular Medicine, University of Massachusetts Medical School; Dept. Of Molecular, Cell and Systems Biology, University of California, Riverside; Center for Genetics of Host Defense, UT Southwestern Medical Center; RNA Therapeutics Institute, University of Massachusetts Medical School; Department of Molecular Medicine, Scripps Research Institute; Faculdade de Medicina, Universidade de Lisboa, Portugal

## Abstract

Eukaryotic cells regulate 5’ triphosphorylated (ppp-) RNAs to promote cellular functions and prevent recognition by antiviral RNA sensors. For example, RNA capping enzymes possess triphosphatase domains that remove the γ phosphates of ppp-RNAs during RNA capping. Members of the closely related PIR1 family of RNA polyphosphatases remove both the β and γ phosphates from ppp-RNAs. Here we show that *C. elegans* PIR-1 dephosphorylates ppp-RNAs made by cellular RdRPs and is required for the maturation of 26G-RNAs, Dicer-dependent small RNAs that regulate thousands of genes during spermatogenesis and embryogenesis. PIR-1 also regulates the CSR-1 22G-RNA pathway and has critical functions in both somatic and germline development. Our findings suggest that PIR-1 modulates both Dicer-dependent and - independent Argonaute pathways, and provide insight into how cells and viruses use a conserved RNA phosphatase to regulate and respond to ppp-RNA species.

## INTRODUCTION

Cells can modify and sense the phosphorylation status of RNA 5’ ends to regulate gene expression, control RNA stability, and mediate antiviral defense (Hornung et al., 2006; Kato et al., 2006; Shatkin, 1976). For example, the eukaryotic RNA polymerase, Pol II, recruits a capping enzyme that co-transcriptionally modifies the 5’ end of its RNA products. A key enzymatic modality in this capping enzyme is an RNA triphosphatase domain related to the cysteine phosphatase superfamily of protein and RNA phosphatases (Deshpande et al., 1999; Shatkin, 1976; Takagi et al., 1998; Yuan et al., 1998). After removing the γ phosphate from a nascent transcript, capping enzyme installs a guanine-nucleotide cap that masks the 5’ end from cellular nucleases and sensors that recognize RNAs made by viral polymerases (Shatkin, 1976). Cellular and viral homologs of the triphosphatase domain of capping enzymes include the PIR-1 (<u>P</u>hosphatase that Interacts with <u>R</u>NA and Ribonucleoprotein Particle <u>1</u>) family of RNA polyphosphatases, which catalyze the removal of γ and β phosphates from <u>trip</u>hosphorylated <u>RNA</u>s (ppp-RNA) *in vitro* (Deshpande et al., 1999; Takagi et al., 1998; Yuan et al., 1998). However, the cellular functions and targets of PIR-1 are largely unknown.

The *C. elegans* PIR-1 homolog was identified as a binding partner of the <u>RNA</u> interference (RNAi) factor, Dicer (Duchaine et al., 2006). RNAi plays important roles in regulating gene expression and viral immunity in diverse organisms (Baulcombe, 2004; Hannon, 2002; McCaffrey et al., 2002; Pal-Bhadra et al., 2002). Dicer encodes a multifunctional protein with <u>d</u>ouble-<u>s</u>tranded (ds)RNA-binding motifs, a DExH/D helicase motif, and a bidentate RNase III domain (Macrae et al., 2006); Dicer is known to bind and then process dsRNA substrates into <u>s</u>hort-interfering (si)RNAs or <u>mi</u>cro(mi)RNAs that guide Argonaute co-factors to mediate genetic silencing (Bernstein et al., 2001; Grishok et al., 2001; Hannon, 2002). *In vitro* studies suggest that Dicer is not sensitive to the 5’ phosphorylation status of its substrates (Welker et al., 2011), raising the question of how and why Dicer associates with PIR-1.

In *C. elegans*, Dicer (DCR-1) functions in several small RNA pathways, including the miRNA pathway and the endogenous (exo-) and exogenous (endo-) RNAi pathways triggered by dsRNAs (Duchaine et al., 2006; Grishok et al., 2001; Welker et al., 2011). Upon exposure to exogenous or viral dsRNAs, DCR-1 processes dsRNAs into short 23-<u>n</u>ucleotide (nt) duplex siRNAs with monophosphorylated 5’ ends and 3’ 2-nt overhangs (Ashe et al., 2013; Coffman et al., 2017; Guo et al., 2013). These diced siRNAs are loaded onto the Argonaute RDE-1 (Tabara et al., 1999), which cannot silence targets alone (Gu et al., 2009; Steiner et al., 2009; Yigit et al., 2006). Instead, RDE-1 recruits cellular <u>R</u>NA-<u>d</u>ependent <u>R</u>NA <u>P</u>olymerase (RdRP) to generate/amplify the silencing signal (Pak and Fire, 2007; Yigit et al., 2006). These RdRPs prefer to initiate transcription at C residues located 5’ of a purine on template RNAs, and thus produce 22-nt products that contain a 5’-triphosphorylated (ppp-)G residue, so called 22G-RNAs (Claycomb et al., 2009; Gu et al., 2012; Gu et al., 2009; Pak and Fire, 2007; Yigit et al., 2006). 22G-RNAs are then loaded onto <u>w</u>orm-specific <u>A</u>r<u>go</u>nautes (WAGOs) (Claycomb et al., 2009; Gu et al., 2009). Unlike other Argonautes, which usually bind monophosphorylated (p-)RNA guides, WAGOs directly accommodate the ppp-RNA guides synthesized by RdRP (Claycomb et al., 2009; Gu et al., 2009).

In addition to its key role in the exo-RNAi pathway, DCR-1 also functions in endogenous small RNA pathways (Duchaine et al., 2006; Fire et al., 1998; Grishok et al., 2001; Ruby et al., 2006; Welker et al., 2010). Several genes that function in non-essential endo-RNAi pathways were identified as mutants with <u>e</u>nhanced exo-<u>R</u>NAi (ERI mutants), perhaps because these endo-RNAi pathways compete for downstream components that are limiting for robust exo-RNAi (Duchaine et al., 2006; Fischer et al., 2011; Kennedy et al., 2004; Ruby et al., 2006; Simmer et al., 2003; Simmer et al., 2002; Yigit et al., 2006). The ERI genes and their associated factors define two major endo-RNAi pathways both employing 26-nt antisense RNAs usually starting with G, 26G-RNAs. The ERGO-1 Argonaute and two redundant Argonautes, ALG-3 and ALG-4 (ALG-3/4), engage 26G-RNAs during embryogenesis and spermatogenesis respectively, and like RDE-1 can trigger the biogenesis of the RdRP-mediated WAGO-dependent 22G-RNAs (Conine et al., 2010; Gent et al., 2010; Han et al., 2009; Pavelec et al., 2009; Vasale et al., 2010; Welker et al., 2010; Zhang et al., 2011). Interestingly, 26G-RNAs are RdRP products themselves, however unlike 22G-RNAs, 26G-RNAs are monophosphorylated and are processed by Dicer in the context of the Dicer/ERI protein complex (Duchaine et al., 2006; Thivierge et al., 2011). Precisely how 26G-RNAs are processed by the Dicer/ERI complex and how 26G-RNAs acquire their 5’ mono-phosphorylated state are largely unknown.

The CSR-1 Argonaute, which has been proposed to promote or modulate rather than silence germline gene expression (Claycomb et al., 2009; Gerson-Gurwitz et al., 2016; Seth et al., 2013), engages 22G-RNAs targeting the majority of germline-expressed mRNAs, including many spermatogenesis mRNAs that depend on the ALG-3/4 26G-RNA pathway (Conine et al., 2010; Conine et al., 2013; Han et al., 2009). However, the vast majority of mRNAs targeted by CSR-1 22G-RNAs are expressed outside of spermatogenesis (Claycomb et al., 2009), and 26G-RNAs targeting these non-spermatogenesis CSR-1 targets have not been identified. Thus, it is not known whether 26G-RNAs and Dicer regulate non-spermatogenesis CSR-1 22G-RNAs.

Here we show that *C. elegans* PIR-1 is an RNA polyphosphatase required for germline development and for endogenous small RNA pathways. *pir-1* mutants exhibit a strong depletion of 22G-RNA species that depend on ALG-3/4 for their amplification, but also exhibit a striking more than 2-fold reduction in nearly all CSR-1 pathway 22G-RNAs. Recombinant PIR-1, like its vertebrate and viral homologs, removes γ and β phosphates from ppp-RNAs. Catalytically dead PIR-1 binds ppp-RNAs but not p-RNAs *in vitro*. Null and catalytically dead *pir-1* mutants exhibit dramatically delayed larval development and male and hermaphrodite infertility. PIR-1 copurifies with the DCR-1/ERI complex and PIR-1 activity is essential to make ERI-dependent 26G-RNAs that engage ALG-3/4. Our analyses suggest a model whereby 26G-RNAs are made in a unique phased manner by successive rounds mRNA processing by the Dicer/ERI complex, and that PIR-1 promotes this mechanism by removing a diphosphate group from the 5’-end of 26G-RNA precursors, likely to facilitate loading into Argonautes. Our findings implicate PIR-1 as a regulator of endogenous Argonaute pathways that process their small-RNA cofactors from RdRP products.

## RESULTS

### *C. elegans* PIR-1 is an RNA polyphosphatase

Previous studies have shown that vertebrate and viral homologs of PIR-1 have polyphosphatase activity that depends on a conserved cysteine in the catalytic motif H<u>C</u>X_5_RXG (**Figure 1A**; Deshpande et al., 1999; Takagi et al., 1998; Yuan et al., 1998). To characterize the enzymatic activity of *C. elegans* PIR-1, we purified recombinant wild-type (WT) PIR-1 protein, as well as recombinant mutant PIR-1(C150S), in which the catalytic cysteine is replaced with serine (**Figure S1A**). We then incubated ppp-RNAs with these recombinant proteins, and assessed the 5’ phosphorylation status of reaction products using <u>Terminator</u>® exonuclease, which degrades 5’ p-RNA but not diphosphorylated (pp-) or ppp-RNAs. WT PIR-1, but not PIR-1(C150S), efficiently converted ppp-RNAs into substrates that were degraded by Terminator exonuclease (**Figure 1B**). WT PIR-1 also dephosphorylated ppp-RNAs duplexed with RNA or DNA (**Figure 1C, Figure S1B** and **S1C**). Notably, PIR-1(C150S), but not WT PIR-1 protein, remained bound to ppp-RNA but not p-RNA substrates in electrophoretic mobility shift assays (**Figure 1D** and **S1C**); this shift was indeed caused by PIR-1(C150S) instead of any contaminations, as confirmed by the Western blot analysis (**Figure 1E**). This mobility shift was detected using 50 mM Tris-Cl buffer (pH 8.0) but not when native protein gel buffer containing 25 mM Tris and 192 mM Glycine (pH 8.3) was used instead (**Figure S1D**), indicating that PIR-1(C150S) binds ppp-RNA non-covalently. Thus, like its vertebrate and viral homologs PIR-1 is an RNA polyphosphatase that converts ppp-RNA to p-RNA. Whereas WT PIR-1 rapidly releases p-RNA products, the catalytically dead PIR-1(C150S) selectively binds and remains bound to ppp-RNA substrates.

**Figure 1.**
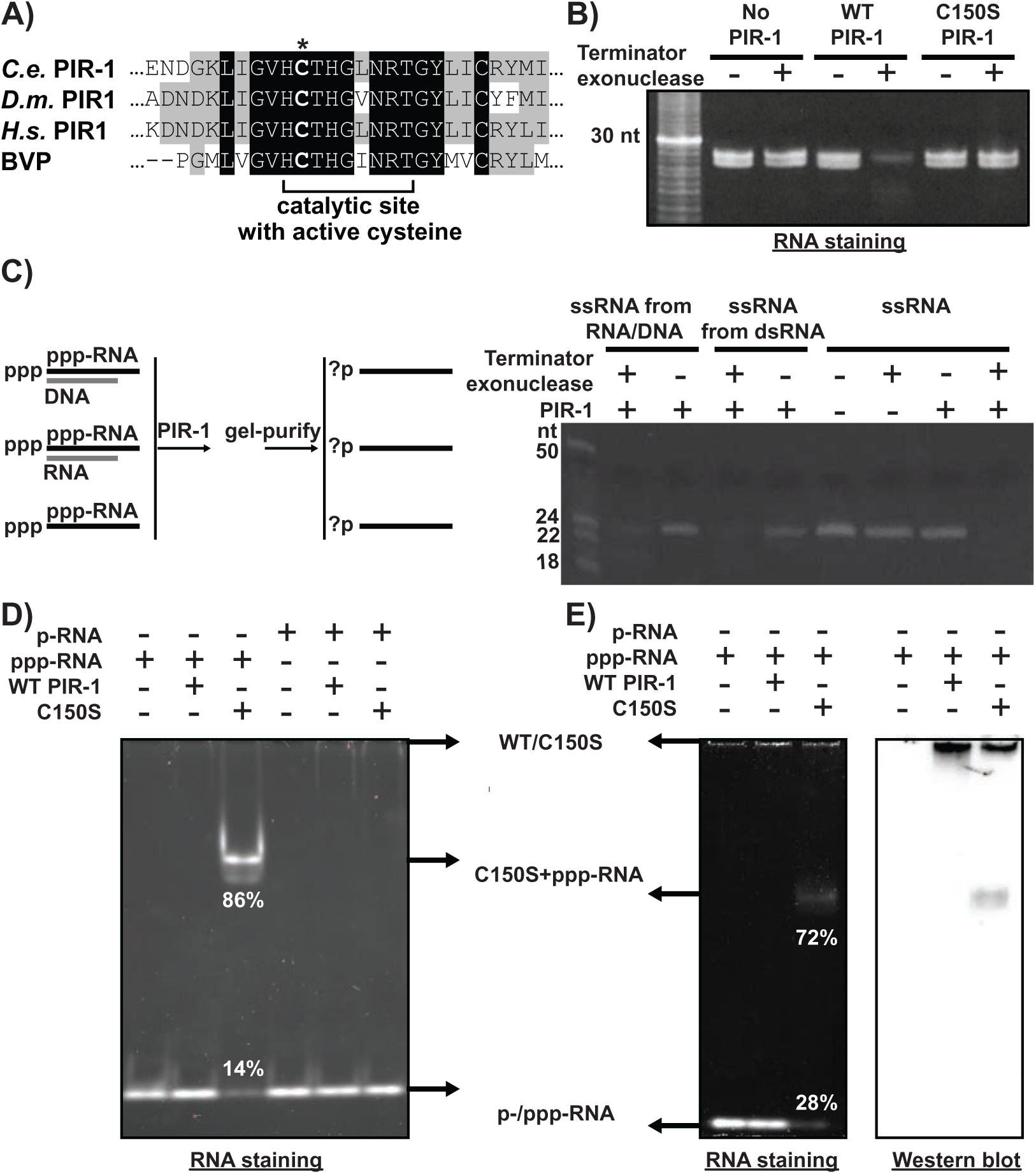
PIR-1 is an RNA polyphosphatase. **A)** The active site alignment of Baculovirus phosphatase (BVP) and PIR-1 orthologs from *C. elegans* (*C.e.*), *Drosophila* (*D.m.*), and human (*H.s.*), with an asterisk indicating the catalytic cysteine, and gray and black colors representing partly and completely identical residues, respectively. **B)** Recombinant WT PIR-1 but not C150S mutant protein converts *in vitro* transcribed 26-27 nt single-stranded ppp-RNAs to p-RNAs, which are then destroyed by Terminator exonuclease. **C)** Single-stranded ppp-RNAs were annealed with a complementary RNA or DNA oligo, digested with WT PIR-1, gel-purified to remove the oligos, and digested with Terminator. **D)** C150S PIR-1 binds single-stranded ppp-RNAs but not p-RNAs while WT PIR-1 binds neither of them stably. The result was resolved on a 15% native PAGE gel and visualized with SYBR Gold RNA staining. **E)** SYBR Gold RNA staining and Western blot analysis verify that C150S but not WT PIR-1 binds and co-migrates with single-stranded ppp-RNAs, as resolved using a 15% native PAGE gel with SYBR Gold RNA staining or using Western blot to detect His-tagged PIR-1, respectively. Arrows indicate the positions of substrate/product RNAs, WT/C150S PIR-1, and PIR-1/RNA complexes. **See also Figure S1.**

### PIR-1 associates with the ERI complex

*C. elegans* PIR-1 was previously identified as a DCR-1-interacting protein (Duchaine et al., 2006). To characterize PIR-1 complexes, we performed PIR-1 immuno<u>p</u>recipitation (**IP**) and analyzed the immunoprecipitates using <u>Mu</u>lti<u>d</u>imensional <u>P</u>rotein Identification <u>T</u>echnology (MudPIT; Schirmer et al., 2003). To facilitate the identification of proteins that specifically interact with PIR-1, we rescued a *pir-1*(*tm3198*) null mutant with a *pir-1::gfp* transgene and labeled the *pir-1::gfp* worms with light nitrogen (^14^N). In parallel, we labeled control WT worms with heavy nitrogen (^15^N). We then mixed ^14^N-labeled *pir-1::gfp* worms with an equal number of ^15^N-labeled control worms, prepared worm lysates, and immunoprecipitated PIR-1::GFP using anti-GFP antibodies. We analyzed the GFP immunoprecipitates by MudPIT and identified candidate PIR-1 interactors as proteins with a minimum of 10 spectral counts for ^14^N-labeled peptides and no spectral counts for ^15^N-labeled peptides. These studies revealed that PIR-1 interacts with the core proteins of the ERI complex (**Table 1** and **S1;** Fischer et al., 2011; Gabel and Ruvkun, 2008; Kennedy et al., 2004; Pavelec et al., 2009; Simmer et al., 2002; Thivierge et al., 2011; Timmons, 2004; Zhang et al., 2011). Similar results were obtained using a *pir-1::3×flag-*rescued strain and FLAG IP (**Table S1**).

**Table 1.** PIR-1 Interactors Identified in PIR-1::GFP IP Using Gravid Adult Worms.

| **Protein** | **Amino Acid Number** | **Spectral Counts<sup>a</sup>** | **Protein Coverage** |
| --- | --- | --- | --- |
| PIR-1 | 233 | 202 | 42.9% |
| DCR-1 | 1845 | 137 | 31.2% |
| RRF-3 | 1780 | 68 | 18.0% |
| DRH-3 | 1119 | 59 | 23.1% |
| ERI-5 | 531 | 24 | 22.0 |
| RDE-4 | 385 | 13 | 28.6 |
| ERI-3 | 578 | 16 | 10.7 |
| ERI-1<sup>b</sup> | 582 | 7 | 12.7% |

<sup>a</sup>The number of tandem mass spectra matching peptides derived from each protein
<sup>b</sup>The spectral count is less than 10.

Using western blot analyses, we confirmed that DCR-1, DRH-3, RRF-3, ERI-1b, and RDE-8 interact with PIR-1 at all developmental stages (**Figure 2A, 2B, S2A** and **S2B**). PIR-1 did not co-IP with ERI-1a (**Figure 2A**), an isoform of ERI-1 known to processes the 3′ end of 5.8S rRNA (Gabel and Ruvkun, 2008). Several Argonaute-dependent small RNA pathway factors that are not part of the ERI complex, including the 3′-to-5′ exonuclease MUT-7, the RdRPs RRF-1 and EGO-1, and the Argonautes CSR-1 and WAGOs, were not detected in PIR-1 immunoprecipitates (**Figure 2A, 2B, S2A, S2B, Table 1 and Table S1**). These Western blot studies identified two PIR-1 isoforms, PIR-1a and PIR-1b, which differ in size by approximately 2 to 4 kDa on denaturing polyacrylamide gels (**Figure 2 and S2**). The molecular basis for this mobility difference remains to be identified. Both isoforms were detected at all larval and adult stages. However, only PIR-1b was detected in embryos, where it associated with several components of the ERI complex (**Figure 2B**). The association of PIR-1b with the ERI complex was confirmed by gel filtration chromatography in which PIR-1b associated with a >440-kDa complex that included DCR-1, DRH-3, ERI-1b, and RDE-8 (**Figure 2C**). Genetic analyses revealed that the interaction between PIR-1 and the ERI complex depends on DCR-1 and DRH-3 but not on ERI-1 or RDE-4 (**Figure S2B-D**). As expected, reciprocal immunoprecipitation of DRH-3 or DCR-1 pulled down PIR-1 (**Figure S2E;** Duchaine et al., 2006; Gu et al., 2009). Interestingly, PIR-1a::GFP expression as detected by GFP IP appeared to depend on *drh-3(+)* activity while PIR-1b::GFP expression required *dcr-1(+)* activity (**Figure S2B** and **S2C**).

**Figure 2.**
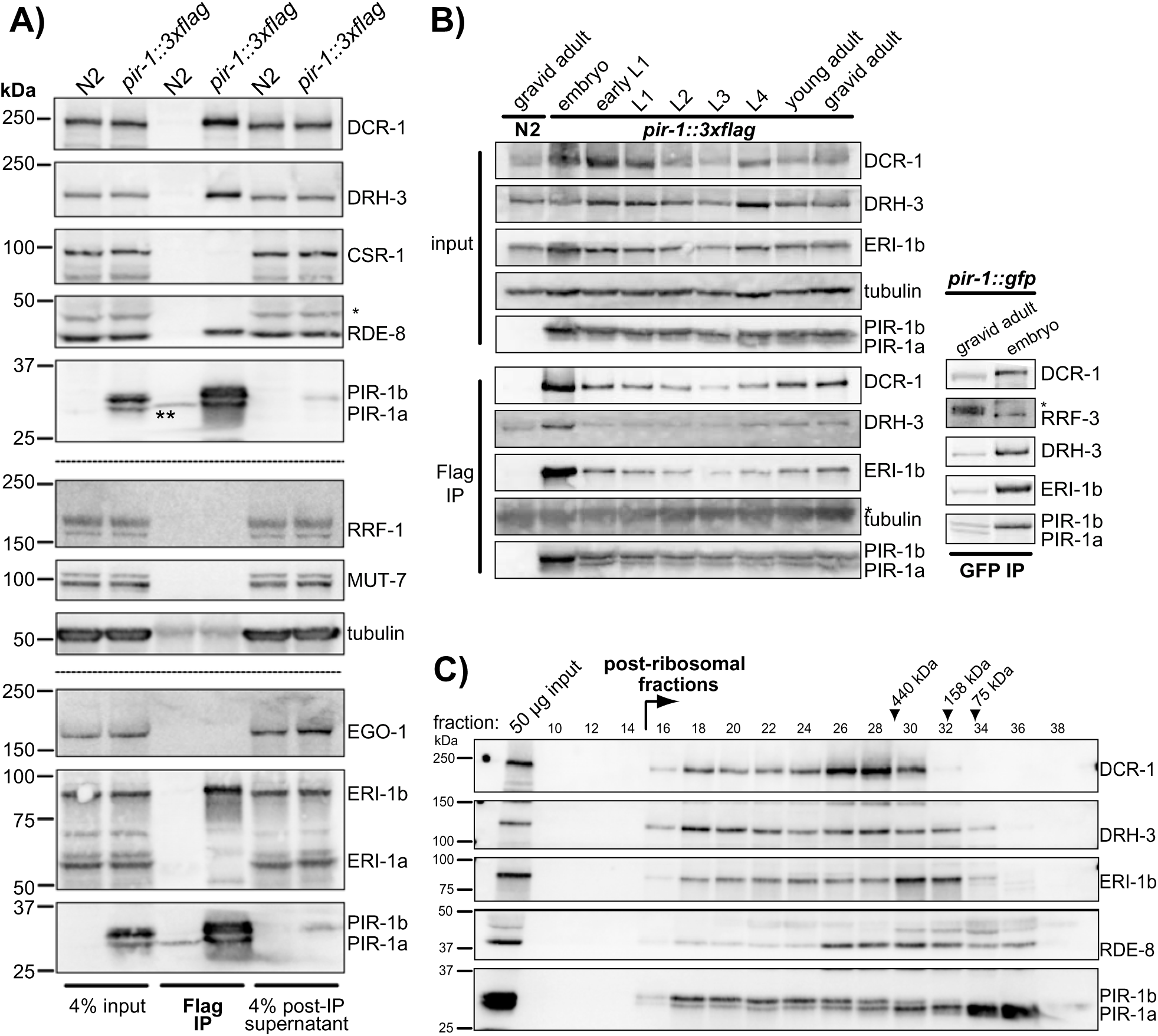
PIR-1 Interacts with the ERI Complex. **A)** Western blot analyses of proteins, as indicated, in PIR-1 IP from *pir-1::3xflag-*rescued young adults and control N2 worms show that PIR-1 exhibits two isoforms, PIR-1a and PIR-1b, and that PIR-1 interacts with DCR-1, DRH-3, RDE-8, and ER1-1b. The * and ** indicate unspecified bands. **B)** Western blot analyses (left) of proteins, as indicated, in PIR-1 IP from *pir-1::3xflag-*rescued worms across developmental stages and from gravid N2 worms. The * in the tubulin panel indicates a signal caused by non-specific binding of the secondary antibody to the heavy chain of anti-Flag antibody used in the IP. Western blot analysis (right) of proteins in PIR-1 IP from single-copy *pir-1::gfp-*rescued worms at gravid adult and embryo stages. Only PIR-1b is expressed in embryos. The * indicates an unspecified band. **C)** Gel filtration of total protein extracts from *pir-1::3xflag*-resucued adult worms followed by Western blot analyses. Arrows indicate the size of fractions based on the molecular weight standards. **See also Figure S2, Table 1 and Table S1.**

### PIR-1 is an essential protein broadly expressed in nucleus and cytoplasm

A previous study identified PIR-1 as a Dicer interactor and described a mutation, *pir-1(tm1496),* which causes a fully penetrant larval lethal phenotype at the early L4 stage (Duchaine et al., 2006). The *tm1496* deletion also removes the promoter and part of the neighboring essential gene *sec-5* (**Figure 3A**), perhaps contributing to the early L4 arrest phenotype. To further explore the function of PIR-1 and its role in Dicer/ERI complexes, we therefore generated a second deletion allele (*tm3198*) and a catalytic C150S mutant allele (*wg1000*; **Figure 3A**). Both of these new alleles caused identical fully penetrant phenotypes. Homozygotes matured to late larval and adult stages and were invariably sterile (**Figure 3** and see below). The *tm3198* allele deletes the first intron and most of the second exon of *pir-1* (**Figure 3A**), which is expected to shift the *pir-1* open reading frame and cause premature translation termination. As expected, reverse transcription and <u>q</u>uantitative <u>PCR</u> (RT-qPCR) analyses revealed that the *tm3198* allele abolishes *pir-1* mRNA expression but does not alter the expression of the neighboring *sec-5* mRNA (**Figure S3A**). Finally, the lethal phenotypes associated with *tm3198* were fully rescued by a single-copy *pir-1::gfp* fusion gene driven by the *pir-1 promoter* and 3’ UTR, indicating that the *tm3198* phenotypes result from the loss of *pir-1(+)* activity.

**Figure 3.**
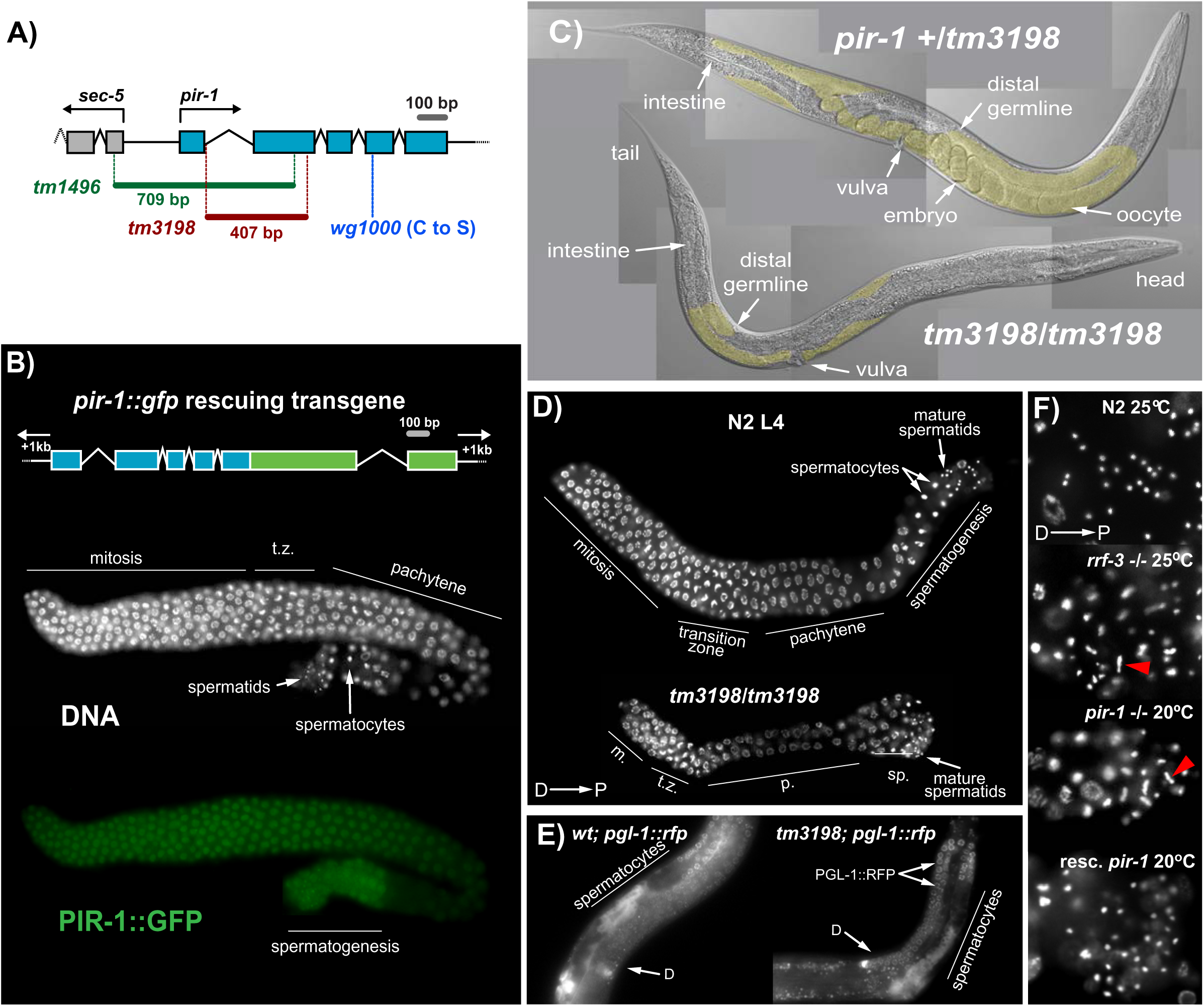
PIR-1 is essential for somatic and germline development. For all figures, ‘D’, distal; ‘P’, proximal; ‘m.’, mitosis; ‘t.z.’, transition zone; and ‘p.’, patchytene. All images use hermaphrodites. **A)** Diagram of WT, *tm1496*, *tm3198* and *wg1000 pir-1* alleles. **B)** Diagram of the transgene used for the rescue experiment and images of a fixed L4 germline from the rescued strain stained with DAPI and anti-GFP. **C)** Images of a live *pir-1* +/*tm3198* worm and a terminally arrested live *tm3198/tm3198* worm grown at 20°C for 96 hours. Germlines highlighted in yellow are partly concealed by intestine tissues. **D)** Images of the fixed DAPI-stained germlines of a late L4 N2 worm and an arrested *pir-1* (*tm3198/tm3198*) worm grown at 20°C. **E)** Images of a live N2 L4 worm and a live arrested *pir-1* mutant worm both expressing fluorescent PGL-1::RFP, which marks P granules and is absent from germline cells differentiating into sperms. **F)** Images of the proximal portions of fixed DAPI-stained L4 germlines from N2, arrested *pir-1* mutant, *rrf-3* mutant, and *pir-1* mutant rescued (resc.) with a single-copy *pir-1::gfp* transgene. Compact round dots represent mature spermatid DNA. Red arrowheads point to aberrant DNA bridges between dividing spermatocytes, which are absent from WT and rescued *pir-1* mutant worms. **See also Figure S3.**

RT-qPCR analyses revealed that in adults with fully developed gonads and embryos, *pir-1* mRNA levels are much higher than in larval stages (**Figure S3B**). Analysis of PIR-1::GFP revealed nuclear and cytoplasmic staining in most germline and somatic cells (**Figure 3B/S3C&D**). In the germlines of L4-stage hermaphrodite worms (i.e., during spermatogenesis), PIR-1::GFP was uniformly expressed at high levels in germ cells from the proliferative mitotic zone to the meiotic mid-pachytene region (**Figure 3B**). PIR-1::GFP levels were reduced in germ cells transitioning through diplotene and meiosis I and II (i.e., through the bend in the ovotestis), and then increased again just before cells begin spermatogenesis. In adult hermaphrodites (i.e., during oogenesis), PIR-1::GFP was also highly expressed in the distal germline and reduced through the bend in the ovotestis, but we did not detect PIR-1::GFP signal in maturing oocytes nor in the embryonic germline. PIR-1::GFP was detected in most somatic nuclei throughout development, exhibiting the highest level in the large polyploid nuclei of intestinal cells (**Figure S3C)**.

The majority of *tm3198* homozygotes (62%) arrested as sterile adults (**Figure S3E**), frequently with a protruding vulva and occasionally ruptured at the vulva (**Figure 3C**/**S3F**). Approximately 21% of worms made deformed oocytes, but none made progeny (**Figure S3F**). Approximately one quarter of *tm3198* animals arrested as viable L4-like larvae that survived for nearly a normal life span with apparently normal motility. Close examination of the germlines of these L4-like arrested larvae revealed features typical of normal L4 germline including a mitotic zone, a transition zone, an extended zone of meiotic nuclei undergoing pachytene, and a spermatogenic zone including spermatocytes and spermatids (**Figure 3D**). A PGL-1::RFP reporter was expressed in a WT pattern throughout the distal germline but not in the proximal spermatogenic region of these arrested L4-like worms, suggesting that they transitioned properly to spermatogenic gene expression (**Figure 3E**). We noticed that many dividing spermatocytes in *pir-1* germlines exhibited abnormal meiotic figures, indicative of DNA-bridging (**Figure 3F**). Similar defects were previously described for mutants in ERI components. For example, loss-of-function mutations in *rrf-3*, *eri-1*, and *eri-3*, and the helicase-domain mutant *dcr-1(mg375)* have all been reported to cause similar DNA-bridging phenotypes when grown at 25°C (**Figure 3F**). These ERI pathway mutants all make defective spermatids (Conine et al., 2010; Han et al., 2009; Simmer et al., 2002; Timmons, 2004). To summarize, *pir-1* mutants exhibit a spectrum of defects at larval and adult stages similar to, and in some respects, such as the larval arrest and oogenesis defects, more severe than other Dicer-ERI complex co-factors.

### PIR-1 is not required for miRNA or piRNA biogenesis

We next explored how *pir-1* mutations affect endogenous small RNA levels. To obtain large numbers of *pir-1* homozygotes, we used a strategy to select against heterozygotes in which *pir-1* is covered by the inversion balancer *mnC1.* Three redundant glutamate-gated chloride channels (AVR-14, AVR-15, and GLC-1) render *C. elegans* sensitive to the nematicidal drug ivermectin (Dent et al., 2000). We crossed *pir-1* into an *avr-14(ad1302)*; *avr-15(ad1051)*; *glc-1(pk54)* triple mutant (*avr3x*) background, and balanced *pir-1* with an *mnC1* balancer that also carries a rescuing *avr-15(+)* transgene (Supplementary Strain List). In the presence of ivermectin, the *pir-1/mnC1* heterozygotes (which express AVR-15) arrest as L1 larvae, but *pir-1* homozygotes (which do not express AVR-15) grow to late larval stages and adulthood (see Experimental Procedures; Dent et al., 2000; Duchaine et al., 2006). We grew synchronized populations of *pir-1* homozygous or control (*avr3x* or N2) worms to extract RNA and generate small RNA libraries for high throughput sequencing. We noted that *pir-1* mutants grew more slowly, both in size and developmental landmarks (e.g., adult cuticle and vulval differentiation), so we prepared samples from *pir-1* mutants grown for 3 days or for 7 days to attain parity in developmental stage with WT populations. To obtain a snapshot of all the different classes of Argonaute-associated small RNAs, we pretreated the small RNA samples with <u>T</u>obacco <u>A</u>cid <u>P</u>yrophosphatase (**TAP**) or with purified recombinant PIR-1 protein, both of which convert ppp-RNAs to p-RNAs. This approach allowed us to simultaneously recover p-RNAs including 26G-RNAs, miRNAs, and piRNA/21U-RNAs, and ppp-RNAs including 22G-RNAs.

Analysis of the small RNA sequencing data revealed that miRNA and piRNA species were largely unaffected in *pir-1* mutants (**Figure 4A**). Comparing small RNAs from temporally matched *pir-1* and control populations (i.e., *pir-1* and *avr3x* on ivermectin for 3 days), we found that *pir-1(tm3198)* expressed more miRNAs but fewer piRNAs and 22G-RNAs (when normalized to total genome mapping reads, including all authentic small RNA species). These findings are likely explained by the developmental delay of *pir-1(tm3198)* animals which causes a relatively smaller germline-to-soma ratio in the 3-day old *pir-1* worms: miRNAs are abundant in the soma, whereas 21Us and most 22G-RNA species are abundant in the germline (**Figure 4A**). Consistent with this idea, in 7-day *pir-1(tm3198)* animals, which appear developmentally similar to 3 day WT or *avr3x* worms, piRNAs and miRNAs were increased to a similar level, and in proportion to the corresponding decrease in 22G-RNA levels (**Figure 4A**). Moreover, developmentally matched (7-day) *pir-1* and control (3-day) *avr3x* worms expressed similar levels of DCR-1 and PRG-1 proteins, factors required for generating miRNAs and binding piRNAs respectively. These observations suggest that the biogenesis of miRNAs and 21Us is temporally delayed in *pir-1* mutants but is not likely to be directly regulated by PIR-1(+) activity (**Figure 4B**). Indeed, when we normalized our small RNA data to piRNA levels, we observed similar levels of miRNAs in control worms and developmentally matched *pir-1* mutants, but 22G-RNA levels were significantly lower in *pir-1* worms (**Figure 4C**). Thus *pir-1* mutants do not exhibit defects in miRNA levels, consistent with our finding that the seam cell numbers (16 on each side of the worm) and adult alae differentiation (hallmarks of miRNA function; are not perturbed in *pir-1* mutants.

**Figure 4.**
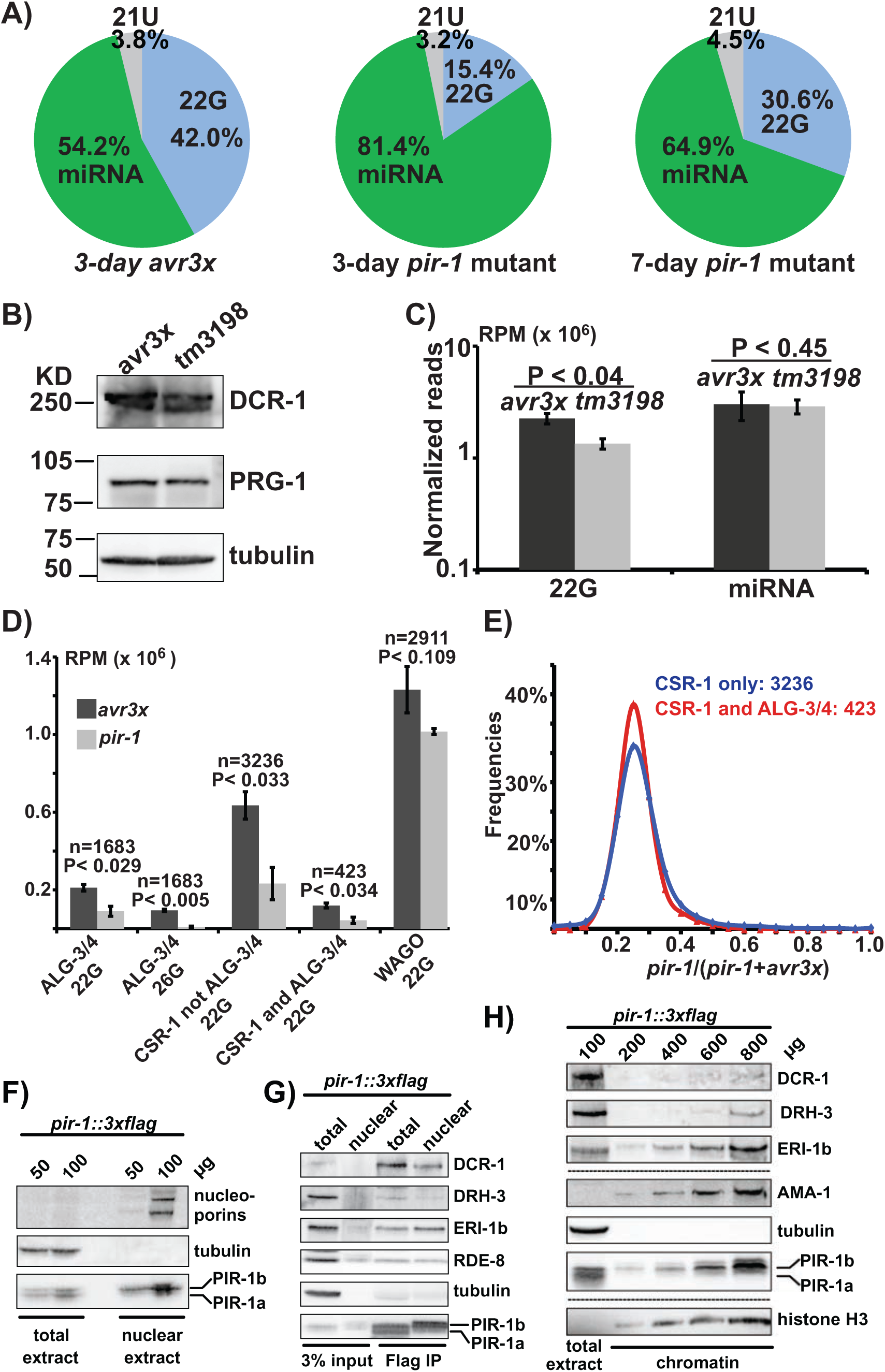
PIR-1 is required for the biogenesis of 26G-RNAs and non-WAGO-bound 22G-RNAs. **A)** Small RNAs from the *pir-1*(*tm3198/tm3198*) mutant and control *avr3x* were cloned with TAP treatment, and normalized to the total reads of 22G-RNAs, miRNAs, 26G-RNAs, and 21U-RNAs. For each strain, the small RNA composition was calculated as the average of two replicas. **B)** Western blot analyses indicate that the *avr3x* control and *pir-1* mutant express similar amount of DCR-1 and PRG-1, as normalized to tubulin. **C)** Comparison of 22G-RNAs and miRNAs between the *pir-1* mutant grown for 7 days and the *avr3x* at a similar developmental stage. ‘RPM’: reads per million. The read number was normalized to the total 21U-RNA reads. The P values were calculated based on two replicas each of WT and *pir-1* mutant samples using unpaired student’s t-test (one-tailed for 22G-RNA and two-tailed for miRNA)**. D)** Total 26G-RNAs and/or 22G-RNAs derived from each small RNA pathway with the ‘n’ number of target genes were obtained as the averages of two replicas of *avr3x* or *pir-1* mutant samples, normalized to total 21U-RNA reads, and used to calculate the one-tailed P values of Student’s t-test. Each positive or negative error bar represents one standard error. **E)** For each gene, the ratio of 22G-RNAs in *pir-1* (*tm3198/tm3198*) to total 22Gs in *pir-1* and *avr3x* was obtained, and then a histogram of these ratios was generated using 20 ratio bins. **F)** Western blot analyses of proteins, as indicated, in total and nuclear extracts from *pir-1::3xflag-*rescued young adults, with nucleoporin as a positive control for nuclear enrichment and tubulin as a negative control. **G)** Western blot analyses of proteins, as indicated, in the PIR-1 IP from total and nuclear extracts of *pir-1::3xflag-*rescued young adults with tubulin as a negative control. **H)** Western blot analyses of potential chromatin-interacting proteins, as indicated, using increasing amounts of purified chromatin from *pir-1::3xflag-*rescued young adults. AMA-1 (large subunit of RNA polymerase II) and histone H3 were used as positive markers for chromatin association with tubulin as a negative control. **See also Figure S4 and Table S2.**

### PIR-1 is required for ERI pathway 26G- and 22G-RNAs

Among the most dramatically affected small RNA species in *pir-1* mutants were 26G-RNAs that depend on the ALG-3/4 Argonautes. These 26G-RNAs are templated from the mRNAs of 1683 target genes, including many genes that play critical roles during spermatogenesis (**Table S2;** Conine et al., 2010). We found that 26G-RNAs were approximately 10-fold less abundant in *pir-1* mutants than in WT populations (normalized to 21U-RNA levels; one-tailed t-test, P < 0.005; **Figure 4D** and **Figure S4A**). Moreover, 22G-RNAs that are amplified downstream of ALG-3/4 targeting (Conine et al., 2010) were also significantly lower in *pir-1* mutants (∼2.4-fold; one-tailed t-test, P < 0.0029; **Figure 4D** and **Figure S4B**). ALG-3/4-independent WAGO 22G-RNAs were not significantly downregulated in *pir-1* mutants (one-tailed t-test, P < 0.109, or two-tailed P <0.218, **Figure 4D** and **Figure S4C;** Conine et al., 2010; Han et al., 2009; Pavelec et al., 2009).

### All CSR-1-bound 22G-RNAs are reduced in *pir-1* mutants

The CSR-1 Argonaute engages 22G-RNAs targeting thousands of germline mRNAs. Roughly 11% (∼423) of CSR-1 target genes are also targeted by the ALG-3/4-dependent ERI pathway (**Table S2;** Conine et al., 2010). However, most CSR-1 target genes have no known upstream Argonautes. We found that compared to WT worms, *pir-1* mutants make significantly (∼3-fold) fewer 22G-RNAs for both categories of CSR-1 target genes (one-tailed t-test, P <0.034 and 0.033, respectively **Figure 4D** and **Figure S4D**). Moreover, we found that both classes of CSR-1 22G-RNAs exhibit similar ratios of reads in the mutant to total reads in the mutant and WT (mutant/[mutant+WT]), with the same medians and similar variances (**Figure 4E**), suggesting that PIR-1 may regulate all CSR-1 targets.

Consistent with the idea that the catalytic activity of PIR-1 plays a role in the biogenesis of germline small RNAs, we found that the *pir-1(C150S)* catalytic mutant exhibits the same spectrum of small RNA changes observed in the null mutants (**Figure S4E**).

### The ERI complex and 26G-RNAs copurify with nuclei

To examine whether nuclear localization of PIR-1 relates to its role in 22G-RNA biogenesis, we examined whether the ERI complex and 22G-RNAs copurify with nuclei. To monitor the purity of nuclear extracts we measured the enrichment of nucleoporin and depletion of tubulin (**Figure 4F;** see Extended Experimental Procedures). Western blot analyses revealed that PIR-1b but not PIR-1a preferentially accumulated in nuclei (**Figure 4F**). PIR-1b, ERI-1b, and RDE-8 were detected in input and PIR-1 IP samples from nuclear extracts; DCR-1 and DRH-3, however, were only detected in the nuclear extracts after PIR-1 IP (**Figure 4G**). Moreover, PIR-1b, DCR-1, DRH-3, ERI-1b, and RDE-8 were detected in the chromatin fractions, which enrich histone H3 while completely depleting tubulin (**Figure 4H**). Taken together these findings suggest that PIR-1b and other ERI components associate with nuclei and/or chromatin.

We also monitored small RNA levels in nuclear fractions. We found that 21Us were only slightly enriched in the nuclear samples, while 26G-RNAs were significantly enriched (Wilcoxon Signed Rank Test, one-tailed, P < 0.0001 for both species). In contrast, WAGO and CSR-1 22G-RNA species were significantly depleted from the nuclear fraction (Wilcoxon Signed Rank Test, one-tailed, P < 0.0001 for each group of 22Gs; **Figure S4F**). We found that miRNAs were neither enriched nor depleted. These findings suggest that 26G-RNAs are produced and/or function in nuclei.

### 26G-RNAs are generated in a phased manner

26G-RNAs are unique among *C. elegans* small RNA species in that their biogenesis depends on both RdRP and Dicer. However, why they are longer than typical Dicer products and how their 5’ ends become mono-instead of triphosphorylated as is typical of other *C. elegans* RdRP products remains mysterious. To investigate the role of PIR-1 in 26G-RNA biogenesis we used bioinformatics to analyze the distribution patterns of small RNAs associated with 26G-RNA target sites. To do this we compiled a metagene analysis of all available ALG-3/4 and ERGO-1 26G-RNA target sequences centered on the 26G-RNA and including about 40-nt upstream and downstream sequences. We then analyzed small RNAs mapping to this interval including both antisense small RNAs (RdRP-derived) and sense small RNAs (from mRNA cleavage). The frequency of each small RNA species was plotted according to its 5’ nt position, and color-coded according to its length (**Figure 5** and **Figure S5A**). The position of the C-residue of the mRNA corresponding to the 5’ G of the antisense 26G-RNA was defined as –1. As expected, for both ALG-3/4 and ERGO-1 26G-RNA pathways, the most abundant antisense species were 26G-RNAs located at the –1 position (**Figure 5 and S5A**). Interestingly, this analysis revealed additional phased 26G-RNA peaks located at approximately 23-nt intervals upstream and downstream of –1 (**Figure 5A** and **Figure S5A**). Mirroring the central and phased 26G-RNAs, we observed an identical distribution pattern of mRNA (i.e., sense-stranded) fragments that likely correspond to Dicer products (**Figure 5** and **S5A**). For the ALG-3/4 pathway, most of these mRNA fragments were 22 nucleotides long (sense 22mer-RNA) with their 5’ ends at –23 and their 3’ ends at –2, just upstream of the –1 C residue (**Figure 5A** and **S5A**). For the ERGO-1 pathway the most abundant sense-stranded small RNAs were 19 nucleotides long, with 5’ ends at –23 and 3’ ends at –5 (**Figure 5B**). These findings suggest that an associated nucleolytic activity removes the –1 C residue (and a few additional nucleotides for ERGO-1 templates) after it templates 26G-RNA initiation (see Discussion). For both the ALG-3/4 and ERGO-1 pathways, the sense RNA 5’ ends align 3 nucleotides downstream of the 26G-RNA 3’ ends. Taken together, these findings suggest that the sense RNAs positioned at –23 in the metagene analysis represent a signature of Dicer processing on duplex 26G-RNA precursors (see Discussion).

**Figure 5.**
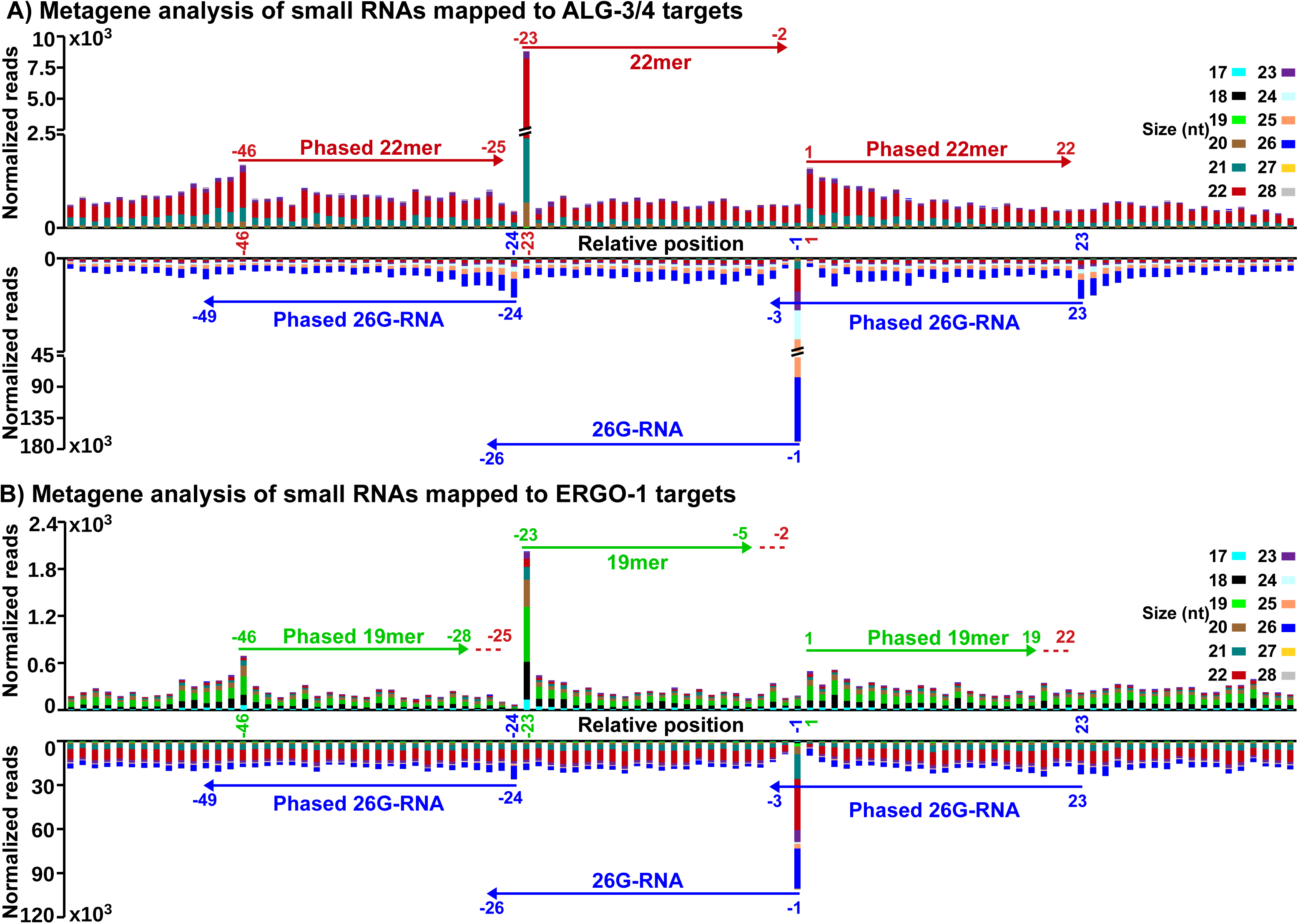
Metagene analysis of small RNAs around 26G-RNA loci. In the ALG-3/4 (**A**, *fog-2* male) and ERGO-1 (**B**, WT embryo) pathways, the position of each 26G-RNA is represented by its very 5’ G, defined as -1 using template mRNA nt C; small RNA reads (’Y’ axis) of various sizes mapped to each position of the template -60 (59 nts upstream of -1) to 40 region (’X’ axis) are obtained for each 26G-RNA and accumulated for all 26G-RNAs. In both (**A**) and (**B**), top panel represents sense RNAs derived from mRNAs with peak sizes at 22 and 19 nts respectively; bottom represents antisense RNAs with peaks at 26 nts; dotted lines represent potential degradation events for generating sense 19mer-RNAs from potential 22mer-RNAs in the ERGO-1 pathway. All coordinates are relative to mRNA -1 C; read number is normalized to total 21U-RNAs for germline expression and total miRNAs for somatic expression in the ALG3/4 and ERGO-1 pathways respectively. **See also Figure S5.**

The above analysis suggests that template mRNAs are processed stepwise by RdRP and Dicer, with RdRP initiating at a C residue and then re-initiating recursively at the first available C residue after each Dicer cleavage event. We further tested this idea by simulating 26G-RNA biogenesis on a computer-generated transcriptome containing random RNA sequences and 26G-RNA densities similar to those in our data sets. Simulated 26G-RNAs were generated on targets by initiating at a randomly selected C residue and then recursively at the first C residue at least 23-nt upstream of the initial template C, propagating the 26G-RNA synthesis toward the 5’ end of target mRNAs in a unidirectional manner. Strikingly, this simulation produced exactly the same metagene pattern observed in our experimental data including the overall shape, the symmetric phased distribution, the loss of phasing at distances greater than 40 nts, and other minor details (**Figure S5B-D**). More details and findings of this simulation analysis are provided in the Supplementary Discussion with Figure S6.

### *pir-1* mutants exhibit defects in 26G-RNA maturation

We next utilized the 26G-RNA metagene sequence space described above to analyze *pir-1* mutants. We found that both 26G-RNAs and the presumptive dicer-mediated sense 22mer-RNA species (positioned at -23 in the metagene) were dramatically reduced in the *pir-1* mutants (**Figure 6A**). Interestingly, 26G-RNAs were disproportionately reduced (**Figure 6B**). The ratio of 26G-RNAs to -23 22mer-RNAs was 31:1 for WT animals, 3:1 for the *pir-1* null mutant and 11:1 for the *pir-1(C150S).* Phased 26G-RNAs were not detected. These finding suggest that PIR-1 activity is required both for the processivity of the Dicer ERI complex and for the maturation of antisense 26G-RNAs.

**Figure 6.**
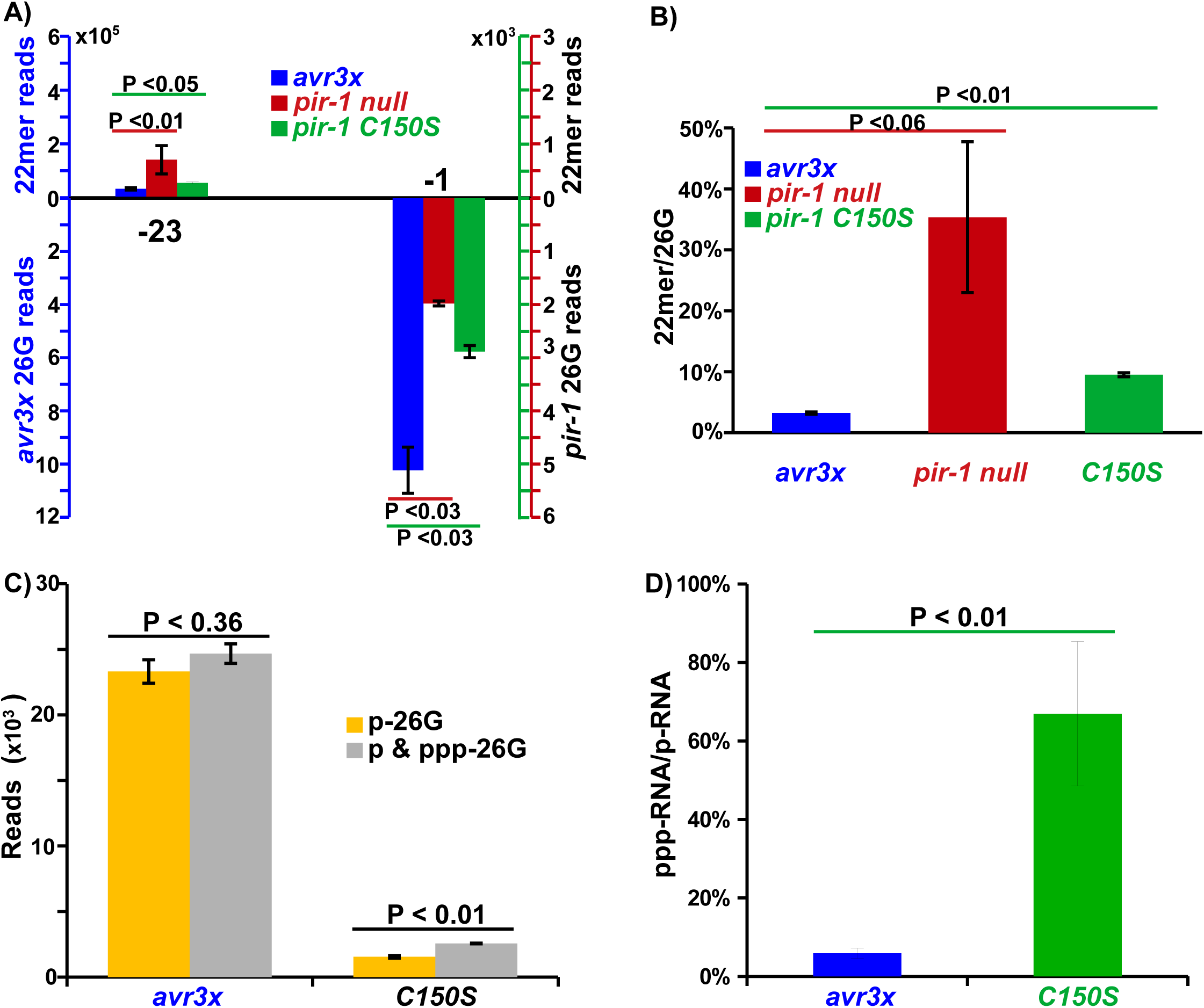
Comparison of small RNAs in the ALG-3/4 pathway in WT and *pir-1* mutants. **A)** Only sense 22mer-RNAs at -23 and 26G-RNAs at -1 were displayed in the metagene analyses (See Figure 5) of small RNAs around ALG-3/4 26G-RNA loci. The small RNAs were cloned using TAP or recombinant PIR-1 treatment. To visually compare the 22mer-RNA/26G-RNA ratio, the reads in the WT (blue) refers to the left ‘Y’ axis (blue) and those in the *pir-1* null (red) and catalytic (green) mutants refer to the right ‘Y’ axis (red and green), which is 1/200^th^ of the left ‘Y’ axis. **B**) The ratios of the above 22mer-RNA to 26G-RNA were obtained for WT and *pir-1* mutants using two replicas of each sample. **C)** Comparisons of p- and ppp-26G-RNAs in WT and *pir-1* catalytic mutants: p- and ppp-26G-RNAs were cloned together with recombinant PIR-1 treatment, which converts ppp-RNAs to p-RNAs for cloning; p-26G-RNAs alone were cloned without recombinant PIR-1 treatment. **D)** Ratios of ppp-26G-RNA to p-26G-RNA in the WT and *pir-1* catalytic mutant. All P values (one-tailed for A, B and D; two-tailed for C) were obtained using unpaired Student’s t-test based on two replicas of each sample; each positive or negative error bar represent one standard error; small RNA reads were normalized to total 21U-RNAs; reads or ratios were averaged using two replicas. **See also Figure S6.**

Since PIR-1 is an RNA phosphatase, one possible explanation for the above finding is that dephosphorylation of the 26G-RNA precursors promote maturation. If so, we reasoned that the 26G-RNAs in animals expressing the PIR-1(C150S) catalytic mutant should exhibit an increase in the relative amount of ppp-26G-RNA. To explore this possibility, we generated small RNA sequencing libraries using a ligation-dependent method that requires a 5’ monophosphate for efficient cloning (Gu et al., 2011; Li et al., 2019). For each mutant and WT sample, we prepared libraries with or without pretreating the RNA with recombinant PIR-1. As expected, most (94%) 26G-RNAs in WT worms were recovered without PIR-1 digestion when normalized to those with recombinant PIR-1 treatment, suggesting that these 26G-RNAs bear 5’ monophosphate. In contrast, we found that ∼40% of 26G-RNAs present in the *pir-1* mutants were resistant to ligation-dependent cloning unless treated with recombinant PIR-1, suggesting that they contain a 5’ triphosphate group (Figure 6C and 6D). Taken together these findings suggest that PIR-1 dephosphorylates 26G-RNA precursors and is required for efficient 26G-RNA maturation by the Dicer ERI complex.

## DISCUSSION

Eukaryotic cells can sense and modify structural features of RNAs to regulate their stability and functions, and to distinguish self-from viral-RNAs. For example, the Dicer protein binds dsRNAs and processes them into duplexed siRNAs and miRNAs that engage Argonaute proteins to mediate sequence-specific viral immunity and mRNA regulation. Conversely, the human RIG-I protein, which contains a Dicer-related helicase domain, detects duplex ppp-RNAs produced by viral RdRPs and then initiates a non-sequence specific cascade of secondary signals that promote viral immunity (Hornung et al., 2006; Kato et al., 2006). Here we have shown that the Dicer-interacting protein PIR-1, like its human and insect virus homologs, removes the β and γ phosphates from ppp-RNAs *in vitro*, generating 5’ p-RNAs. *In vivo,* PIR-1 is required for fertility and for the accumulation of 26G-RNAs antisense to hundreds of spermatogenesis mRNAs.

26G-RNAs are an enigmatic species of Dicer product best understood for their role in spermatogenesis where along with their AGO-related Argonaute co-factors, ALG-3/4, they promote spermatogenesis-specific gene regulation and epigenetic inheritance (Conine et al., 2010; Conine et al., 2013; Han et al., 2009). During embryonic development 26G-RNAs engage the Argonaute ERGO-1 to regulate a group of repetitive RNAs of unknown functions. Mutations that inactivate the ERGO-1 pathway cause enhanced RNAi (ERI phenotypes; Fischer et al., 2011; Gent et al., 2010; Han et al., 2009; Kennedy et al., 2004; Simmer et al., 2002; Thivierge et al., 2011; Vasale et al., 2010; Zhang et al., 2011). While our genetic studies only revealed a role for PIR-1 in the larval-stage ALG-3/4 26G-RNA pathway, it is likely, as previously shown for other RNAi components including Dicer and RDE-1 (Parrish and Fire, 2001; Tabara et al., 1999; Tabara et al., 2002), that the embryonic functions of PIR-1, including its possible function in the ERI pathway are rescued in embryos of heterozygous mothers by maternally provided PIR-1(+) activity.

### A model for 26G-RNA biogenesis

26G-RNAs are processed by Dicer from triphosphorylated duplex RNAs templated from mRNAs by the cellular RdRP, RRF-3. Our findings are consistent with a model in which 26G-RNAs are generated in a phased manner along target mRNAs through the recursive re-initiation by RRF-3 at the first available C nt 5’ of successive Dicer-mediated cleavage events (**Figure 7**). After each round of RdRP transcription, a 3’-5’ exonuclease associated with the ERI complex, possibly ERI-1b digests the mRNA, removing the transcription start site C residue (located at –1 in the Model **Figure 7**), to generate a dsRNA with a 1-nt recessed 3’ end. Binding of the mRNA 3’ OH within the PAZ domain of Dicer and engagement of the Dicer helicase domain could then position the enzyme to generate the 5’ end of an mRNA-derived 22mer-RNA (at –23 relative to the initiator C residue) and a ppp-26G-RNA with a 3-nt 3’ overhang (**Figure 7**; see also Welker et al., 2011).

**Figure 7.**
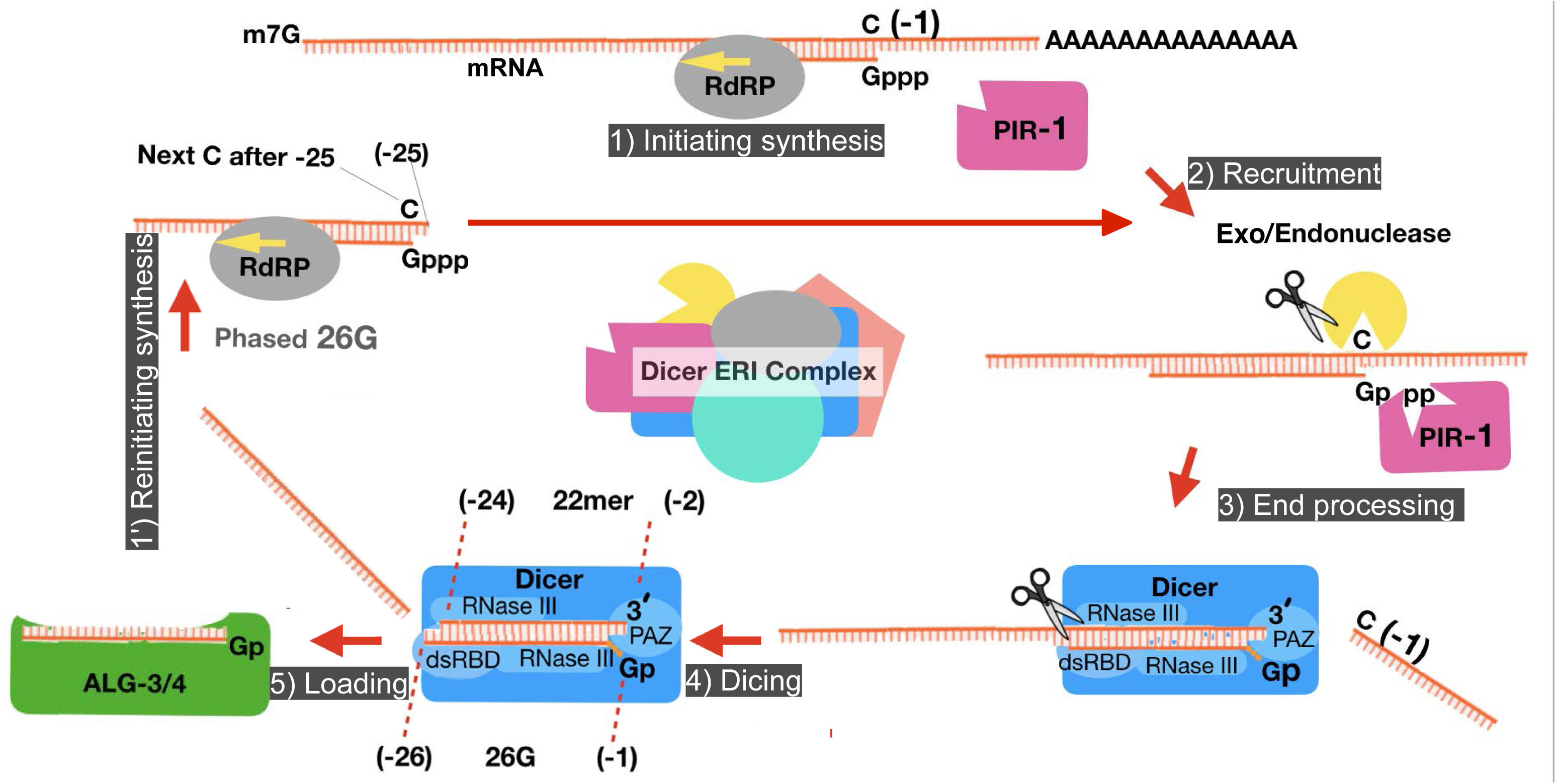
Model of 26G-RNA biogenesis. **Step 1**-iniating synthesis: RRF-3 (RdRP) utilizes mRNAs as templates to synthesize precursor ppp-26G-RNAs, which could be a little bit longer than 26 nts; **Step 2**-PIR-1 recruitment: PIR-1 recognizes/binds the triphosphate group and recruits other factors including Dicer; **Step 3**-end processing: mRNA is processed to generate a one-nt recessive 3’ end lacking -1 C and PIR-1 removes a diphosphate group from precursor 26G-RNAs; **Step 4**-cleavage: Dicer cleaves the dsRNAs to generate duplex siRNAs composed of 26G-RNAs with a 1-nt 5’ overhang and 3-nt 3’ overhang and sense 22mer-RNAs; **Step 5**-26G-RNA loading: 26G-RNAs are loaded to ALG-3/4; **Step 1’**-reinitiating synthesis: RRF-3 reinitiates using the first available template C nt in the 3’ region of each 23nt-shortened mRNA to synthesize a phased precursor 26G-RNA which is then processed via step 2-5 to generate another mature 26G-RNA; the cycle continues until RRF-3 reaches the 5’ end of template mRNAs. Removal of the -1 C could occur after Dicing. The ERGO-1 pathway likely utilizes a similar mechanism in which ERGO-1 binds 26G-RNAs and sense 22mer-RNAs are subjected to degradation, generating a 19mer size peak.

*In vitro* studies suggest that Dicer is not sensitive to the phosphorylation status of the substrate 5’ end (Welker et al., 2011; Zhang et al., 2002). Thus removal of the diphosphate by PIR-1 could occur before or after Dicing (**Figure 7**). Indeed, we found that the levels of sense-stranded 22-mer RNA fragments (presumptive Dicer products) and ppp-26G-RNAs were increased in *pir-1* mutants compared to WT worms. Thus, dicing still occurs in *pir-1* mutants but maturation into p-26G-RNAs appears to be reduced. Perhaps the transfer of the diced product to the downstream ALG-3/4 Argonautes, whose AGO-clade homologs are known to prefer mono-phosphorylated guide RNAs, occurs inefficiently when the diphosphate is not removed.

### PIR-1 exhibits ppp-RNA-specific binding activity

*In vitro* studies on PIR-1 revealed a surprising activity associated with the presumptive catalytically dead C150S lesion. This mutation behaved like a strong loss of function allele, causing small-RNA and developmental defects identical to those caused by a *pir-1* null mutation. However, we found that PIR-1(C150S) nevertheless bound specifically to ppp-RNAs in our gel-shift assays. Structural studies on members of the cysteine phosphatase superfamily to which PIR-1 belongs have shown that during catalysis the cysteine motif generates a covalent cysteinyl-S-phosphate intermediate that is later hydrolyzed in a two-step reaction (Sankhala et al., 2014; Takagi et al., 1998). The substitution of serine for cysteine in PIR-1 C150S replaces the reactive sulfhydrl group of cysteine with a hydroxyl group, preventing formation of the covalent linkage. The finding that this catalytically dead protein retains its ppp-RNA-specific binding activity suggests that substrate recognition is separable from catalysis in PIR-1. Thus it is possible that PIR-1 utilizes its affinity for ppp-RNAs to recognize RRF-3 products and to help recruit Dicer and other ERI complex co-factors to the nascent duplex.

It is interesting to note that a baculovirus-encoded PIR-1 homolog, PTP, functions as a virulence factor that promotes a fascinating behavioral change in infected host caterpillars (Katsuma et al., 2012). Ingested virus spreads to the brain, and the infection eventually causes the caterpillar to migrate to upper foliage, where the dying animal ‘liquifies’—a process thought to maximize dispersal of the virus (Katsuma et al., 2012). Interestingly, *ptp* null mutants were partially defective in brain infectivity and behavioral modification, but PTP C119S mutants supported both activities (Katsuma et al., 2012), suggesting that PTP provides a purely structural capacity to promote virulence, e.g., through its interaction with viral capsid protein (Katsuma et al., 2012). However, if PTP C119S selectively binds ppp-RNA—similar to PIR-1 C150S—then it remains possible PTP C119S interacts with and promotes viral packaging of cellular or viral ppp-RNAs that function as small-RNA cues that alter host behavior. This possibility is particularly intriguing as a growing number of reports have described the modulation of neural and behavioral activity by small RNAs originating in other tissues (Bharadwaj and Hall, 2017; Cai et al., 2018; Hou et al., 2019; Posner et al., 2019).

### PIR-1 is required for robust levels of CSR-1 22G-RNAs

We were surprised to find that *pir-1* mutants exhibit significantly reduced levels of all CSR-1 22G-RNAs. The biogenesis of 22G-RNAs does not require Dicer. Instead 22G-RNAs appear to be produced directly by the RdRP EGO-1 and are then loaded, without further processing, as ppp-RNAs onto their downstream Argonaute co-factors. It is therefore intriguing that CSR-1 22G-RNAs but not WAGO 22G-RNA levels were depleted in *pir-1* mutants. The upstream events in the WAGO 22G-RNA pathway differ from events involved in the CSR-1 pathway. For example, WAGO 22G-RNA biogenesis is initiated by RDE-1 guided by an siRNA processed by Dicer or by Piwi Argonaute (PRG-1) guided by a piRNA. When RDE-1 and PRG-1 bind target mRNAs, they recruit cellular RdRPs that synthesizing WAGO 22G-RNAs (Ashe et al., 2012; Bagijn et al., 2012; Grentzinger et al., 2012; Lee et al., 2012; Pak and Fire, 2007; Shen et al., 2018; Shirayama et al., 2012; Yigit et al., 2006; Zhang et al., 2018). Whether an upstream Argonaute functions in the CSR-1 pathway is unknown. Although 26G-RNAs have not been detected for most CSR-1 targets, perhaps they are short lived, developmentally restricted, for example to larvae, or are simply very low abundance, and have been missed. Further investigation will be required to understand this connection between PIR-1 and CSR-1.

### PIR-1 plays critical roles in larval development

A striking feature of the PIR-1 mutant phenotype is the dramatically slowed development of homozygous larvae. *pir-1* mutants take nearly twice as long as WT animals to reach adulthood, and yet appear to behave like otherwise healthy WT larvae for the course of a nearly normal lifespan of 16 to 18 days. Other mutants that perturb 26G-RNA pathways do not exhibit delayed development phenotypes (Conine et al., 2010; Fischer et al., 2011; Gent et al., 2010; Han et al., 2009; Kennedy et al., 2004; Pavelec et al., 2009; Simmer et al., 2002; Thivierge et al., 2011; Timmons, 2004; Vasale et al., 2010; Zhang et al., 2011). Perhaps the presence of cellular RdRPs in *C. elegans* makes RNA phosphatase activity essential in order to ensure that accumulating ppp-RNA products do not compromise RNA homeostasis or activate heretofore unknown innate immunity mechanisms. Conceivably, the absence of PIR-1 activity could trigger a diapause that is normally triggered only when an excessive cytoplasmic accumulation of viral ppp-RNAs overwhelms the capacity of PIR-1 and Dicer mediated immunity. If reversible, a diapause in response to ppp-RNA might allow animals to postpone reproduction until after the viral infection is cleared. The role of PIR-1 in development and its possible function in anti-viral immunity will require further investigation.

## EXPERIMENTAL PROCEDURES

### Worm Strains

The *C. elegans* Bristol N2 strain and its derivatives used in this study were cultured essentially as described (Brenner, 1974). 10-25 µg/L Ivermectin was added to NGM plates for selecting *pir-1* homozygous worms. A list of strains was detailed in the Supplementary Materials.

### RNA Extraction, Cloning and Sequencing

RNA was extracted with TRI Reagent (MRC, Inc.), according to the manufacturer’s protocol. Nuclei were isolated from predominately L4 *avr3x* animals grown on ivermectin, as described in Extended Experimental Procedures. After the second purification using a sucrose solution, nuclear pellets were lysed with TRI Reagent (MRC) for RNA extraction. Cloning of small RNAs and read analysis were performed as described in the Extended Experimental Procedures. Small RNA libraries were prepared essentially as described (Gu et al., 2011; Li et al., 2019). Briefly, ∼1 µg of total RNA was used for cloning small RNAs either via the conventional ligation-based method or the one-pot cloning method; Tobacco Acid Pyrophosphatase (Epicentre, discontinued) or recombinant PIR-1 was used to dephosphorylate ppp-RNAs for cloning ppp-RNAs when needed while no such treatment was required for cloning p-RNAs. Libraries were sequenced using Illumina NextSeq, HiSeq 4000, and Genome Analyzer II.

### Bioinformatics

High-throughput sequencing reads were processed and mapped to *C. elegans* genome and annotations (WormBase release WS215) using Bowtie 0.12.7 (Langmead et al., 2009) and further analyzed using custom PERL scripts as deposited to GitHub at https://github.com/guweifengucr/WGlab_small_RNA_analysis. The Generic Genome Browser was used to visualize the alignments (Stein et al., 2002).

### Immunoprecipitation, Western blot, Proteomics Analysis, and PIR-1 Activity Assays

The IP and Western blot were performed as described previously (Gu et al., 2009). Total RNAs were extracted from IP and input samples and cloned for small RNA analyses using the above protocol. The MudPIT analysis for PIR-1 interactors was detailed in Extended Experimental Procedures.

The PIR-1-mediated dephosphorylation and ppp-RNA binding assays were detailed in Extended Experimental Procedures.

### Microscopy

Imaging of live animals and tissues was detailed in Extended Experimental Procedures.

## ACKNOWLEDGMENTS

W.G. is supported by NIH grant GM124349 and C.C.M. is a Howard Hughes Medical Institute Investigator and is supported by NIH grant GM058800.

## ACCESSION NUMBER

High-throughput sequencing data is available from the GEO DataSets under the series number GSE150690 with a private access token ‘cbwlkmqudjinryh’ for the reviewers.

## SUPPLEMENTARY STRAIN LIST

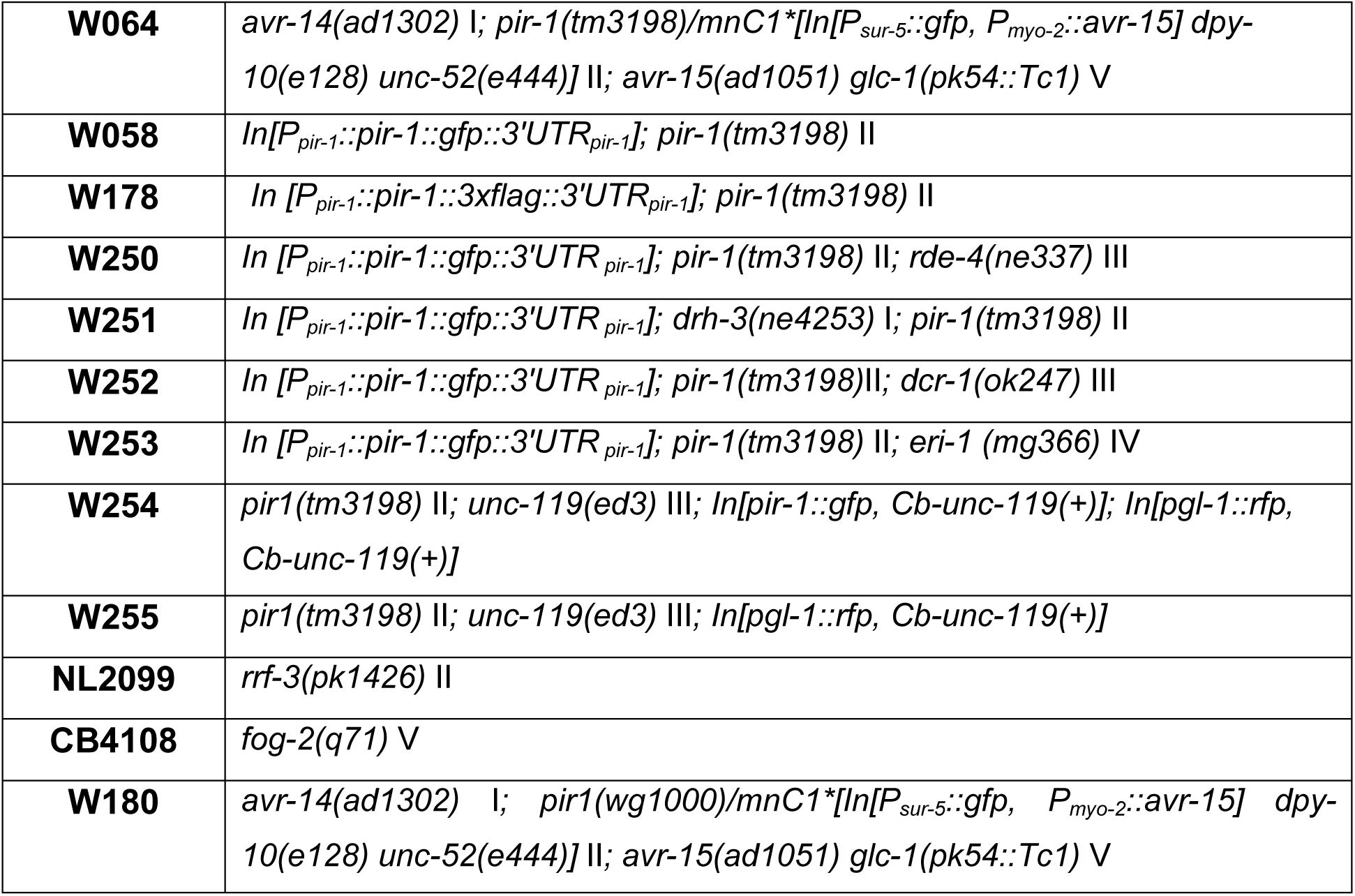

|  |  |
| --- | --- |
| **W064** | *avr-14(ad1302) I; pir-1(tm3198)/mnC1\*[In[P<sub>sur-5</sub>::gfp, P<sub>myo-2</sub>::avr-15] dpy-10(e128) unc-52(e444)] II; avr-15(ad1051) glc-1(pk54::Tc1) V* |
| **W058** | *In[P<sub>pir-1</sub>::pir-1::gfp::3'UTR<sub>pir-1</sub>]; pir-1(tm3198) II* |
| **W178** | *In [P<sub>pir-1</sub>::pir-1::3xflag::3'UTR<sub>pir-1</sub>]; pir-1(tm3198) II* |
| **W250** | *In [P<sub>pir-1</sub>::pir-1::gfp::3'UTR<sub>pir-1</sub>]; pir-1(tm3198) II; rde-4(ne337) III* |
| **W251** | *In [P<sub>pir-1</sub>::pir-1::gfp::3'UTR<sub>pir-1</sub>]; drh-3(ne4253) I; pir-1(tm3198) II* |
| **W252** | *In [P<sub>pir-1</sub>::pir-1::gfp::3'UTR<sub>pir-1</sub>]; pir-1(tm3198)II; dcr-1(ok247) III* |
| **W253** | *In [P<sub>pir-1</sub>::pir-1::gfp::3'UTR<sub>pir-1</sub>]; pir-1(tm3198) II; eri-1 (mg366) IV* |
| **W254** | *pir1(tm3198) II; unc-119(ed3) III; In[pir-1::gfp, Cb-unc-119(+)]; In[pgl-1::rfp, Cb-unc-119(+)]* |
| **W255** | *pir1(tm3198) II; unc-119(ed3) III; In[pgl-1::rfp, Cb-unc-119(+)]* |
| **NL2099** | *rrf-3(pk1426) II* |
| **CB4108** | *fog-2(q71) V* |
| **W180** | *avr-14(ad1302) I; pir1(wg1000)/mnC1\*[In[P<sub>sur-5</sub>::gfp, P<sub>myo-2</sub>::avr-15] dpy-10(e128) unc-52(e444)] II; avr-15(ad1051) glc-1(pk54::Tc1) V* |

## SUPPLEMENTARY DISCUSSION

### Using a simulation model to recapitulate the biogenesis of 26G-RNAs

The symmetric distribution patterns of 26G-RNAs and sense 22mer-RNAs in our analysis (**Figure 5A**) suggest that they are synthesized in a ‘recursive’ mode, in which any 26G-RNA can serve as an initiating 26G-RNA to generate another phased 26G-RNA using template sequence at least 23 nts upstream. Each RNA, therefore, is surrounded by another RNA either 23 nts upstream or downstream, unless this RNA is mapped to the very 5’ and 3’ ends of templates.

However, this model seems incompatible with our observation that only two (at -24 and 23 positions in **Figure 5**) rather than multiple clusters of phased 26G-RNAs were generated. Moreover, it is difficult to understand why the small RNA profile in the ALG-3/4 pathway is much more obvious than that in the ERGO-1 pathway (**Figure 5A** vs. **5B**). We propose that these phenomena are likely caused by different small RNA densities and distribution patterns. To test this hypothesis, we designed a simulation algorithm using parameters obtained from the experimental data. In each round of simulation, one thousand of RNA molecules, each with a 1000-nt random sequence containing 21% C (frequency in template mRNAs), are generated; 20 C’s are randomly selected as -1 C’s and used to generate initial 26G-RNAs; if the -24 (23 nts upstream of -1) is C, a phased 26G-RNA is generated; otherwise, the next available upstream template C is selected; the next round of phased 26G-RNAs starting at -47 regions are generated using the same rule and so on (**Figure S5B**). The selected C’s for both the initial 26G-RNAs and phased 26G-RNAs in the ALG-3/4 pathway are limited to the 5’ and 3’ 10% of mRNAs since most ALG-3/4-bound 26G-RNAs are located there (**Figure S4A**). To achieve the best simulation result, each C could fail to generate a 26G-RNA at 30-40% rate, and if a failure occurs, next upstream C can serve as a template nt also with a failure rate of 30-40%. This failure rate, the only parameter not obtained from the experimental data, simply mimics RNA degradation or other competing processes, and was determined based on the best fitting results. To minimize variations due to randomization, we obtained the average results from 100 rounds of simulations.

Using these parameters, the simulation generates similar density of 26G-RNAs as the experimental data in both ALG-3/4 and ERGO-1 pathways (**Figure S5C&D**). The higher density of 26G-RNAs in the ERGO-1 pathway may simply reflect that each initial 26G-RNA can generate more phased 26G-RNAs since the template region is not limited to mRNA ends. The 26G-RNA distribution patterns in the simulation data share the same details as those obtained from the experimental data, especially in the ALG-3/4 pathway; phased 26G-RNAs in the -24 regions form an obvious cluster while phased 26G-RNAs at -47 regions are barely observable, being buried in the background signals (**Figure S5C&D**). The more obvious distribution pattern of 26G-RNAs in the ALG-3/4 pathway may reflect that 26G-RNA biogenesis process propagates less along template mRNAs since they are restricted to the 5’ and 3’ ends, generating more phased 26G-RNAs than other ‘out-of-phase’’ 26G-RNAs proportionally. The simulation control utilizes the same parameters and algorithm but only allows for generating initial 26G-RNAs. As expected, there are no phased 26G-RNA peaks. Although our simulation algorithm perfectly recapitulates the overall distribution of 26G-RNAs, it fails to generate a depletion of 26G-RNAs at the -2 position, as exhibited in the experimental data. We speculate that such depletion could represent a transcription initiation motif for the initial 26G-RNAs.

### Phased 26G-RNAs are not caused by a higher frequency of nt C in the -24 template region

Phased 26G-RNAs may be artificially caused by a specific sequence motif. For example, for any given C on template mRNAs, the 23-nt upstream and downstream regions enrich C nts. This motif, if existing, generates more initial 26G-RNAs, which are randomly selected and mistakenly assigned as ‘phased’ in our analysis. To test this, we examined the distribution of nt C’s flanking any specified C’s (designated as -1) on template mRNAs. This is basically a motif analysis, since each genomic C locus has a weight 1 instead of the number of reads in the high-throughput data analysis which can be considered as an expression analysis (**Figure 5** and **S5**). Interestingly, nt C’s especially in the ALG-3/4 pathway exhibit a regular wave-like pattern with a 1-nt peak followed by a 2-nt valley flanking the -1 C’s, the major peak (**Figure S6)**. However, these flanking peaks and valleys are so tiny that the flanking C contents appear constant at ∼21%, suggesting that the template C distribution cannot generate a semi-phased distribution pattern of 26G-RNAs.

We also performed a similar motif analysis only using template C’s which generate 26G-RNAs in the experimental data. If a 26G-RNA locus is selected, it could represent a phased 26G-RNA, meaning there is an initial 26G-RNA locus at the 23 position and therefore the 23 position enriches C nts. By contrast, the -24 position won’t enrich C nts since for any give -1 C the biogenesis of phased 26G-RNAs at -24 just follows the genomic C frequency, i.e., 21%. As expected, this motif analysis exhibits a sub-peak at the 23 position and no obvious sub-peak at -24 (**Figure S6**). Also consistent with our hypothesis, motif analyses using the simulation data generated similar asymmetric patterns and such patterns were dependent on allowing initial 26G-RNAs to generate phased 26G-RNAs along the template RNAs from 3’ to 5’ (**Figure S6**). In conclusion, this asymmetric motif pattern strengthens our model that RRF-3 synthesizes 26G-RNAs using a ‘recursive’ mode and moves along template RNAs from 3’ to 5’.

Alternatively the above motif analysis may suggest that RRF-3 prefers template C’s with an additional C at the 23 position, generating a sub-peak. However, this model cannot explain the symmetric distribution pattern of phased 26G-RNA reads flanking the initial 26G-RNAs (**Figure 5 and S5**).

### Extended Experimental Procedures

#### MudPIT Analysis

For the ^15^N/^14^N experiment, air-dried pellets were dissolved in 60 µl of 0.1% RapiGest SF (Waters Corporations) in 50 mM ammonium bicarbonate. Proteins were reduced with 5 mM Tris (2-carboxyethyl) phosphine hydrochloride (Sigma-Aldrich) and alkylated with 10 mM iodoacetamide (Sigma-Aldrich). Proteins were digested for 18 hr at 37°C with 0.5 µg trypsin (Promega). The digestion was stopped with 5% formic acid. After 1 hr incubation at 37°C, debris was removed by centrifugation for 30 min at 18,000x *g*.

Protein and peptide identification and protein quantification were performed with Integrated Proteomics Pipeline - IP2 (Integrated Proteomics Applications http://www.integratedproteomics.com/). Tandem mass spectra were extracted from raw files using RawExtract 1.9.9.2 and were searched (both light and heavy) against the WormBase database (WP236) with reversed sequences using ProLuCID. The search space included all fully-tryptic peptide candidates. Carbamidomethylation (+57.02146) of cysteine was considered as a static modification. Peptide candidates were filtered using DTASelect, with the parameters -p 2 -y 1 --trypstat --pfp .01 -DM 10 --DB --dm -in. Quantification was performed using Census.

#### Isolation of Nuclei and Chromatin

All steps were carried out on ice or at 4°C. The frozen worm pellets were resuspended with 2 volumes of Buffer A (10 mM Tris-HCl pH 8.0, 250 mM sucrose, 10 mM MgCl_2_, 1 mM EGTA) without detergents and with protease and phosphatase inhibitors (Roche protease and phosphatase inhibitor cocktail tablets and 1 mM PMSF). For inputs, a fraction of this worm suspension was briefly spun down and Buffer A was replaced with 3 volumes of lysis buffer (25 mM HEPES-KOH pH 7.5, 150 mM KCl, 0.5% NP-40, 0.1% Triton X-100, protease and phosphatase inhibitors) and crushed with a metal douncer on ice with 100 strokes. Lysates were cleared by centrifugation at 10,000x *g* for 15 min. For nuclear isolation, worms were crushed with 30 strokes on ice and debris was removed by centrifugation at 500x *g* for 1 min. This step was repeated; the supernatant was transferred to fresh tubes, and nuclei were collected by centrifugation at 4,000x *g* for 5 min. Nuclear pellets were gently resuspended in at least 10 volumes of Buffer A with inhibitors. 1 ml of nuclear suspension was gently overlayed over 10 ml of sucrose cushion solution (10 mM Tris-HCl pH 8.0, 1M sucrose, 10 mM EDTA) in a 15 ml conical tube and the interphase was gently disrupted by swirling a pipette tip to create a gradient (no more than 1 cm into the sucrose solution). These tubes were centrifuged at 3,200x *g* for ∼1.5 hr, until a uniform white pellet corresponding to nuclei accumulated at the bottom of the tubes. The supernatant was aspirated and nuclei were carefully resuspended in 10-volume Buffer A with inhibitors. A second sucrose flotation step was carried out in 1.5 ml tubes by overlaying 150 µl of nuclei suspension onto 1 ml of sucrose solution, again gently disrupting the interphase. The tubes were spun at 20,000x *g* for 10 min, and the resulting pellets were resuspended in 10 volumes of lysis buffer with detergents and inhibitors (as above for inputs) and crushed with a douncer with 150 strokes to completely disrupt nuclei. This suspension was cleared at 8,000x *g* for 5 min and the supernatant was kept for protein quantification and use as a nuclear extract for IP. Pure chromatin is gray when pelleted. The presence of white patches indicates that intact nuclei remain. When this occurred, the pellet was resuspended in lysis buffer with detergent and crushed a further 100 times until the pellet was uniformly gray. The pellet was then washed once with 10 mM Tris-HCl pH 7.5 with 0.1% Triton X-100 followed by twice with 10 mM Tris-HCl pH 7.5 (with 8,000x *g* centrifugations in between and the last one at 20,000x *g* for 5 min). Pellets were weighed and resuspended in 9 volumes of SDS protein sample buffer with DTT for a 1X final concentration so that the chromatin concentration was between approximately 50-100 µg/µl.

#### Cloning, Expression and Purification of Recombinant PIR-1

Wild-type (WT) or mutant PIR-1 cDNA sequences lacking the first ATG was inserted between the *Nde*I site and *Bam*HI sites of pET-28a (Novagen) in fusion with the 6x Histidine tag N-terminally. The resulting plasmid was transformed into BL21 (DE3) RIL *E. coli* cells, which were grown in 1 liter of LB medium at 37°C to an OD_600_ of 0.4, and induced for 4 hr with 1 mM IPTG at room temperature. Cells were pelleted at 5,000x *g* for 10 min at 4°C and lysed by sonication in 25 ml of lysis/binding buffer (50 mM Tris-HCl pH 7.5, 700 mM NaCl, 5 mM β-mercaptoethanol, 5% glycerol, 15 mM imidazole, 0.01% NP-40). S100 fractions were prepared by ultracentrifugation at 100,000x *g* at 4°C for 1 hr. In a 15 ml conical tube, 2 ml of HisPur beads (Thermo Scientific) were washed 3 times with the lysis/binding buffer and centrifuged at 3,000x *g* between washes. The beads were mixed with the S100 supernatant, transferred to a 50 ml conical tube for rotation at 4°C for 1 hr. Beads were transferred to an empty Poly-Prep chromatography column (Bio-Rad) and washed at 4°C with at least 200 bead volumes of the lysis/binding buffer. Elution was performed at 4°C with 500 µl of imidazole buffer per fraction (50 mM Tris-HCl pH 7.5, 100 mM NaCl, 5 mM β-mercaptoethanol, 5% glycerol, 400 mM imidazole, 0.01% NP-40). Peak fractions were analyzed by 10% SDS-PAGE followed by Coomassie Blue staining. Proteins were dialyzed using 50 mM Tris-HCl pH 7.5, 100 mM NaCl, 1 mM EDTA, 1 mM DTT, 50% glycerol, 0.01% Triton X-100.

#### PIR-1 Activity Assays

To examine the dephosphorylation activity of recombinant PIR-1, a 26-nt ppp-RNA1 (ppp-GGAUCCUUGAAAUGGAACAUCUGAAU) was transcribed *in vitro* with T7 RNA polymerase followed by gel-purification using 15% PAGE/6M urea (two bands of desired size were co-purified). In **Figure1B**, 1 µM of ppp-RNA was co-digested with ∼0.25 µM of recombinant WT or mutant PIR-1 and 0.25 U of Terminator exonuclease (Epicentre) in 10 µl 1X PIR-1 reaction buffer containing 50 mM Tris-HCl (pH 8.0) and 0.1 M NaCl, 2 mM DTT, and 2 mM MgCl_2_ at 30°C for 1 hr. The reaction was stopped by adding formamide gel loading buffer II (Ambion), and run on a 15% PAGE/6 M urea with 0.5X TBE buffer. The RNA was visualized with UV light after staining with SYBR Gold (Thermo Fisher Scientific).

The above *in vitro* transcription predominantly generates byproduct RNAs of much bigger size likely due to template switching when T7 RNA polymerase runs off a template. This prompted us to generate a precursor RNA ppp-GUCAUUCAG AUGUUCCAUUUCAAGGA<u>GGGUCGGCAUGGCAUCUCCACCUCCUCGCGGUCCGA CCUGGGCUACUUCGGUAGGCUAAGGGAGAAG</u>, which contains a Hepatitis delta virus (HDV) ribozyme (underlined) to self-cleave the precursor, generating ppp-RNA2 co-transcriptionally (ppp-GUCAUUCAGAUGUUCCAUUUCAAGGA; Schürer et al. 2002). In **Figure1C**, ppp-RNA2 alone, ppp-dsRNA generated using ppp-RNA2 annealed with an RNA oligo 5’OH-UUGAAAUGGAACAUCUGAAUGAC (the oligo is smaller than ppp-RNA2 and thus can be separated from ppp-RNA2 in gel purification) and ppp-RNA/DNA hybrid generated using ppp-RNA2 annealed with a DNA oligo 5’OH-TTGAAATGGAAC ATCTGAATGAC in 1X PIR-1 reaction buffer (the annealing rate is close to 100% as shown in **Figure S1B**), digested with recombinant PIR-1 using the above reaction condition, and gel-purified to obtain processed ppp-RNA2. Then these processed RNAs were subjected to digestion with 0.05 U of Terminator in a 10 µl PIR-1 reaction buffer at 30°C for 30 minutes, resolved on a 15% PAGE/6M urea, and visualized using SYBR Gold staining.

In the binding assay, recombinant PIR-1 (no Terminator) was incubated with ppp-RNA1 (**Figure 1D and 1E**) or double stranded nucleic acids including ppp-RNA2/RNA oligo or ppp-RNA2/DNA oligo (**Figure S1C and S1D**) using 1X PIR-1 reaction buffer at 20°C for 40 minutes. The reaction was resolved using a 10% native PAGE gel containing 50 mM Tris-HCl (pH 8.0) at room temperature and visualized using SYBR Gold staining.

#### Preparation of Tissues for Microscopy

For visualization of live animals, washed worms were mounted on slides with a 2% agarose pad with M9 buffer containing 0.4% levamisole to paralyze the animals. For live embryos, gravid adults were placed on the agarose pad and cut around the vulva with a fine hypodermic needle. For preparation of tissues for DAPI staining and immunofluorescence were carried out largely according to (Claycomb et al, 2009). For gonad dissection 40 to 50 L4 to young adult worms were picked from plates and washed extensively with 1x Egg Buffer (25 mM HEPES-NaOH pH 7.4, 118 mM NaCl, 2 mM EDTA, 0.5 mM EGTA, 0.1% Tween-20) to eliminate bacteria. Then the buffer was replaced with Egg Buffer containing 0.4 mM levamisole (15-30 µl) and transferred onto an 18×18 mm coverslip. By cutting the animals with the tip of a fine hypodermic needle at either the head below the pharynx or at the tail, gonads (and intestines) were released. An equal volume of fixative solution (3.7% formaldehyde in 1x Egg Buffer without Tween-20) was added and pipetted up and down to further extrude and dissociate germline tissue from the rest of the animals. Fixation was allowed to occur for 5 min at room temperature. All but about 10 µl of solution were removed from the coverslip. The coverslip was picked up by touching the drop at the center of a positively charged slide (VWR VistaVision HistoBond) leaving a small corner of the coverslip protruding from the edge of the slide. Excess solution was wicked away from the edge of the coverslip using torn strips of absorbent filter paper to promote adherence of the tissues to the slide. The sample was then freeze-cracked by placing it on a pre-cooled aluminum block on dry-ice for at least 10 min and quickly flicking the coverslip from the slide using the protruding corner. The slide was immediately dipped in cold (−20°C) methanol in a Coplin jar for 1 min, and then transferred to 1X PBS buffer (10 mM Phosphate pH 7.4, 137 mM NaCl, 2.7 mM KCl) containing 0.1% Tween-20 (PBST) at room temperature. For DAPI staining only, the slide was washed in PBST for 10 min, followed by another 10-min wash with PBST with 0.5 µg/ml DAPI, and a final 30-min wash in PBST, all at room temperature. Slides were mounted by first removing excess buffer from the slides without letting the sample dry completely and then inverting the slide and touching the sample on a drop of 10 µl of Vectashield mounting medium placed at the center of a 22×22 mm coverslip. Excess medium was removed by pressing the inverted mounted slide on a paper towel, and the edges were sealed with transparent nail polish.

For immunofluorescence, the wash step after methanol was followed by 3x 10 min washes in PBST, followed by a blocking step with 0.5% BSA in PBST. For this 100 µl of the solution were added onto the worms and covered with a square Parafilm coverslip to hold the liquid in the sample area. Slides were incubated in a humid chamber at room temperature for at least 30 min. The Parafilm slides were removed by dipping the slide in PBST. 100 µl of primary antibody diluted in blocking solution were placed on the sample and covered with a Parafilm coverslip, and incubated in a humid chamber for 2 hr at room temperature or overnight at 4°C. After 3x 10-min washes in PBST at room temperature the slide was incubated with secondary antibody as described for the primary, followed by the wash steps with DAPI staining and mounting of the slide as described above.

For enhancement of PIR-1::GFP signal, worms were incubated with a 1:100 dilution of anti-GFP mouse monoclonal antibody (WAKO) overnight at 4°C. Secondary antibody incubation was performed for 2 hr at room temperature with a 1:500 dilution of FITC-conjugated donkey anti-mouse (Jackson). For histone H3 a rabbit polyclonal anti-H3 (Cell Signaling) was used at a 1:100 dilution and incubated overnight at 4°C, followed by a 2-hour, room-temperature incubation with 1:500 TRITC-conjugated anti-rabbit antibody (Jackson). Images were acquired with a Zeiss Axioplan 2 microscope using Zeiss AxioVision software.

## SUPPLEMENTARY FIGURE LEGEND

**Figure S1.**
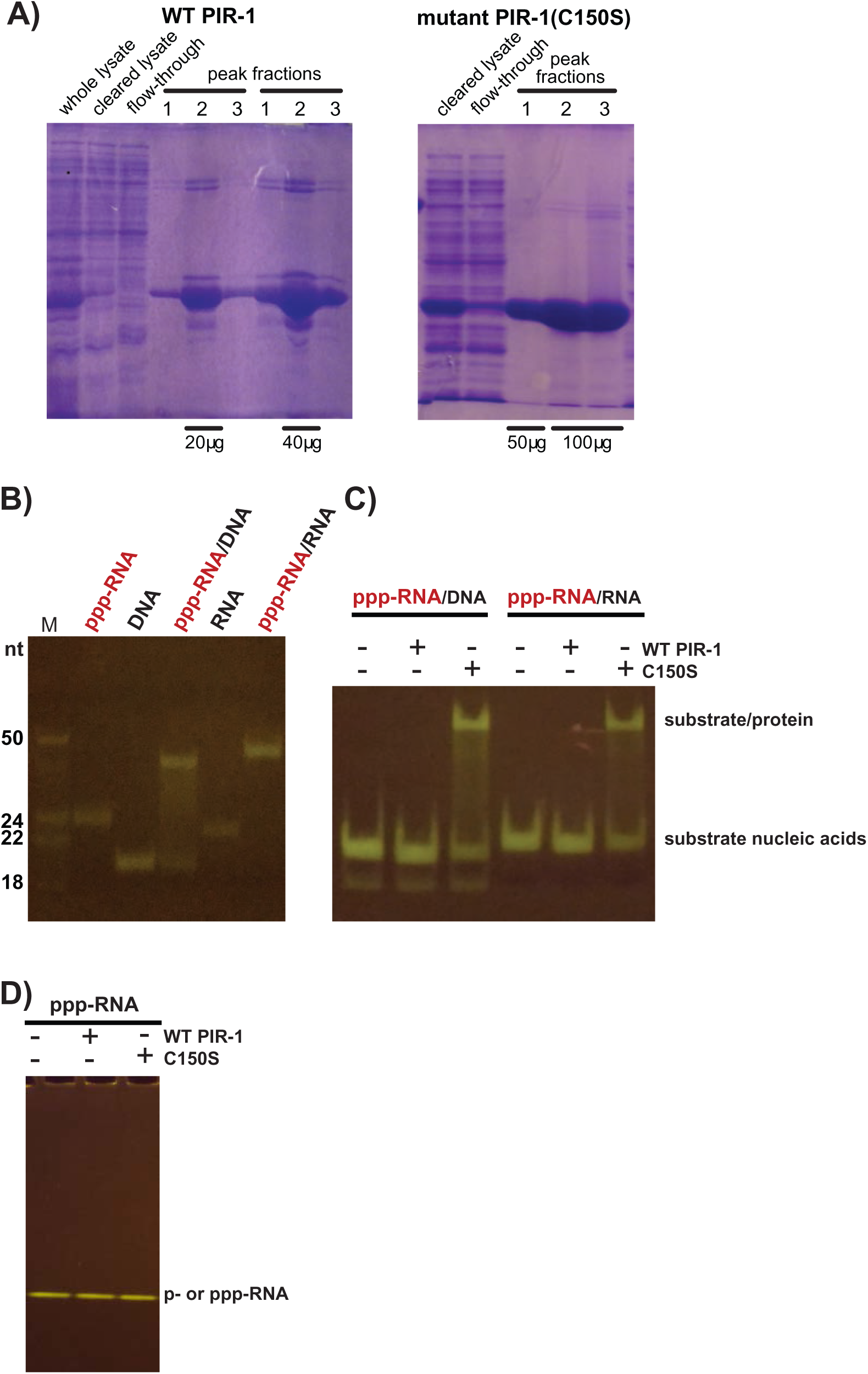
C150 PIR-1 binds ppp-RNAs in double-stranded structures. Related to Figure 1. **A)** The cleared lysates, flow-throughs and elution fractions during recombinant His_6_-tagged PIR-1 purification were resolved on a 12% denaturing protein PAGE gel and visualized with Coomassie Blue staining. **B)** Single-stranded ppp-RNAs were annealed with a complementary RNA or DNA oligo, and resolved on a 15% native PAGE gel to check the annealing efficiency. **C)** WT and C150S PIR-1 were incubated with double-stranded nucleic acids including ppp-RNA/RNA oligo or ppp-RNA/DNA oligo, and then resolved on a native 12% PAGE gel.

**Figure S2.**
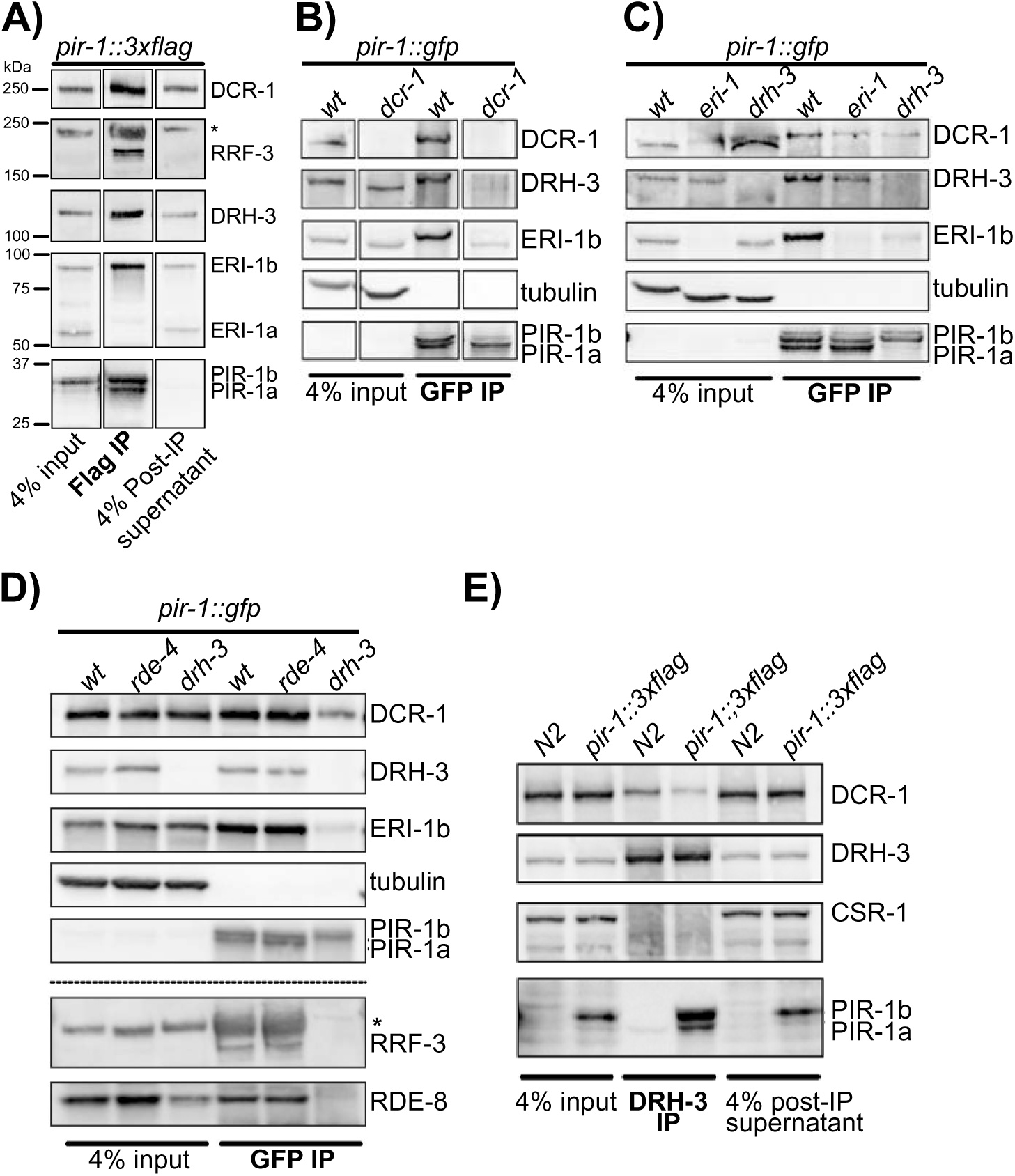
Analysis of PIR-1 isoforms and interacting proteins. Related to Table 1, Table S1 and Figure 2. **A)** Western blot analyses of PIR-1 IP from *pir-1::3xflag*-rescued young adult worms identified PIR-1-interacting proteins including DCR-1, RRF-3, DRH-3 and ERI-1b. **B-D)** Western blot analyses of PIR-1 IPs from WT (*avr3x*) and single-copy *pir-1::gfp-*rescued young adults in *dcr-1, eri-1, drh-3*, *and rde-4* mutant backgrounds. Tubulin was used as a control. **E)** Western blot analyses of DRH-3 IP using N2 and *pir-1::3xflag*-rescued young adult worms identified PIR-1 and DCR-1. Asterisks in panel A and D mark unspecified bands.

**Figure S3.**
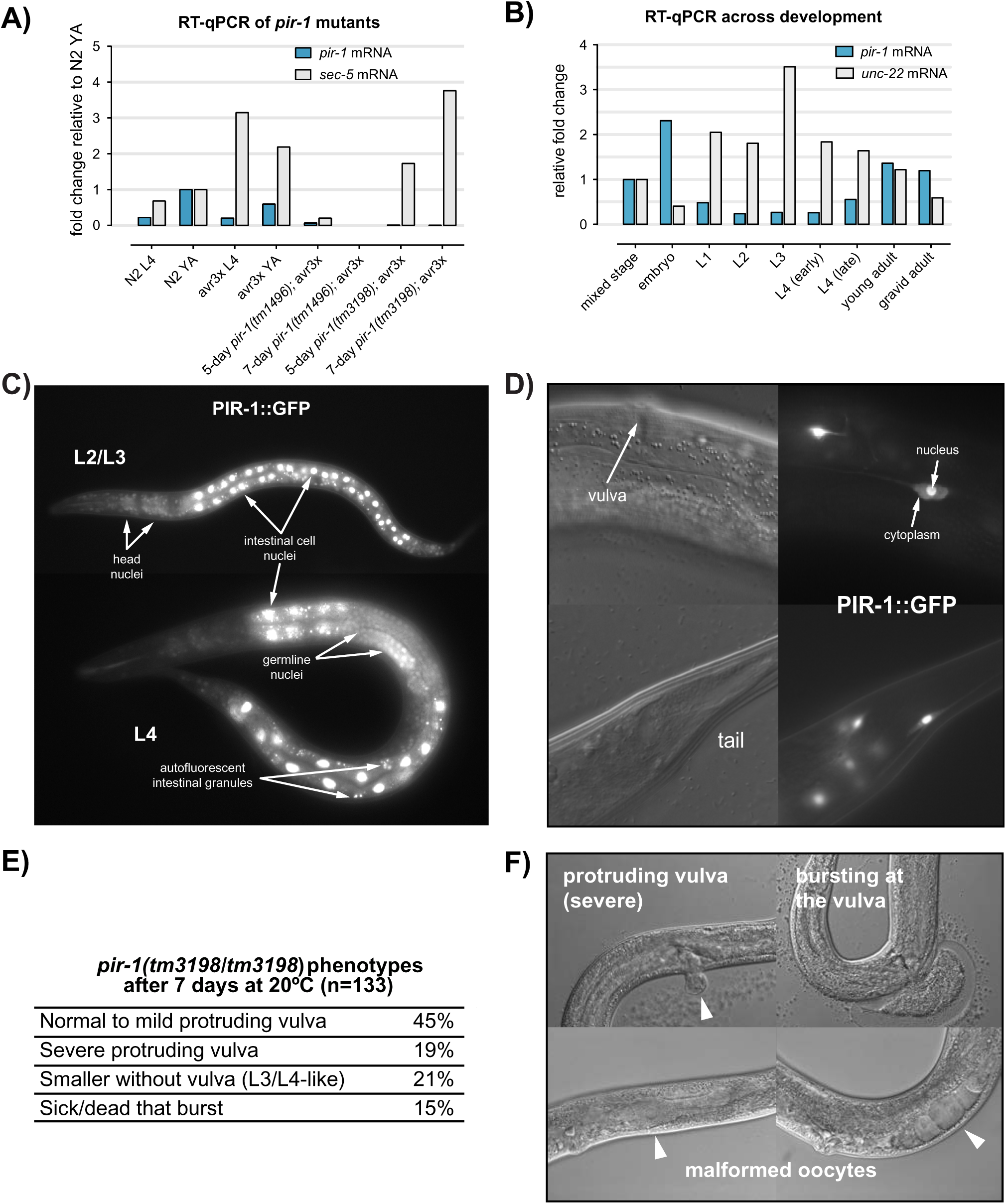
Characterization of *pir-1* mRNA, loss-of-function phenotypes and expression patterns. Related to Figure 3. **A)** RT-qPCR analysis of *pir-1* and *sec-5* mRNA levels normalized to *gpd-2* mRNA levels in the same samples. The fold changes are calculated based on the RNA levels in the young adult (YA) N2 worms. The *pir-1* mutants were counter-selected with ivermectin for five or seven days at 20 °C. **B)** RT-qPCR analysis of *pir-1* and muscle-specific *unc-22* mRNA across developmental stages using N2 worms. 18S rRNA was used for normalization and the fold changes were calculated based on the RNA levels in the mixed stage N2. **C)** Live rescued *pir-1* mutant larvae with an integrated *pir-1::gfp* transgene which was introduced using bombardment, reveal a nearly ubiquitous protein expression pattern. **D)** Images of live non-integrated bombardment lines exhibit high PIR-1::GFP expression in only a few somatic cells with both nuclear and cytoplasmic localization. **E)** Quantification of visible phenotypes exhibited by 133 *tm3198* homozygotes grown for seven days at 20 °C. **F)** Images of live *pir-1* mutant animals exhibiting major phenotypes scored in **E)**.

**Figure S4.**
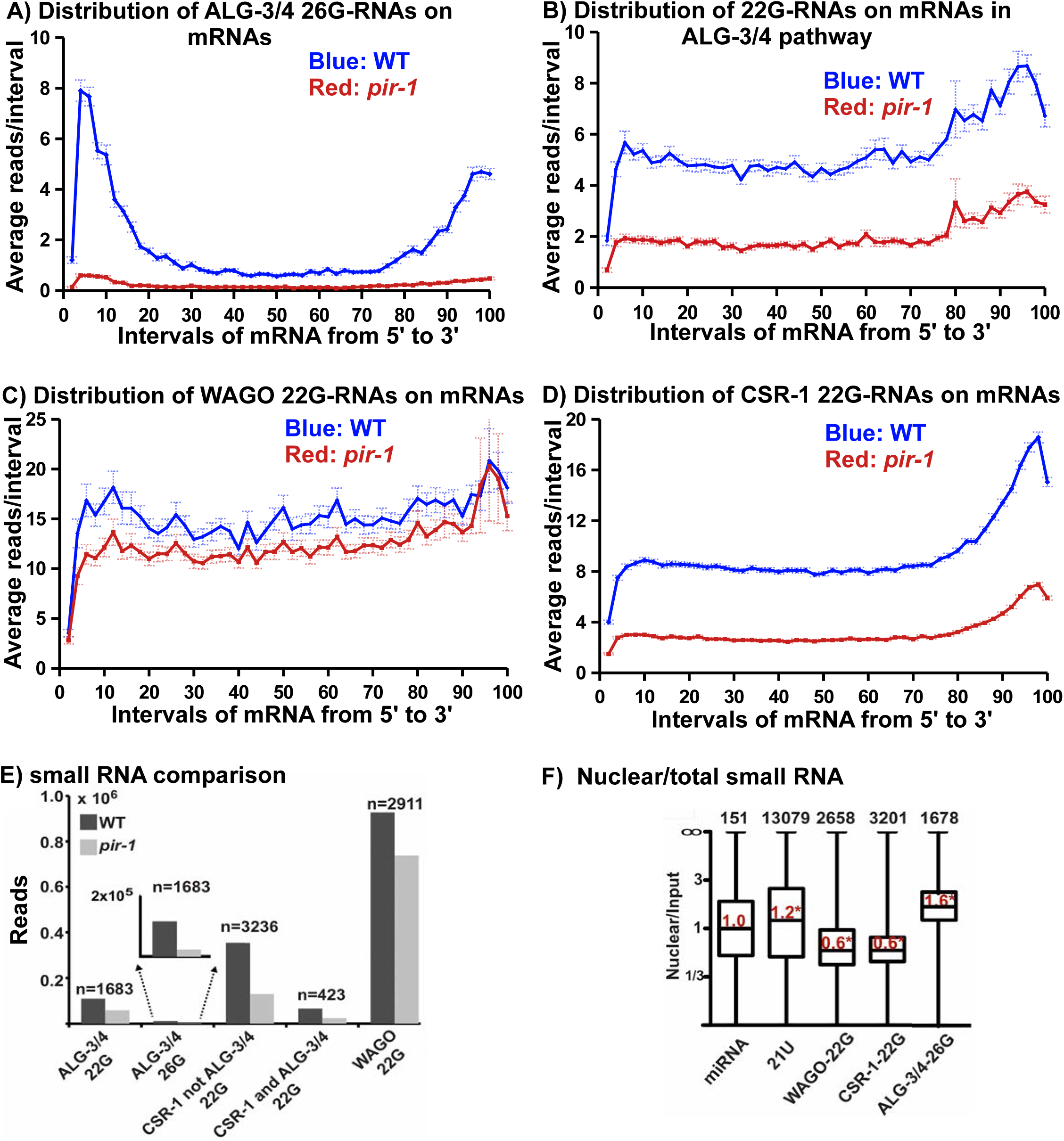
Analyses of small RNAs in *pir-1* null mutants. Related to Figure 4. **A-D)** distribution of small RNAs along mRNA templates: each mRNA is evenly divided into 50 intervals from 5’ to 3’ (’X’ axis); 22 or 26G-RNAs are assigned to each interval after normalization to total 21U-RNAs; a cumulative number is first obtained for each interval using all mRNAs and divided by the total mRNA number to obtain the average (’Y’ axis) in WT (*avr3x*,blue) and *pir-1* mutant (red). **E)** Total 26G-RNAs and/or 22G-RNAs derived from the ‘n’ number of genes in each small RNA pathway in WT (*avr3x*) or *pir-1* catalytic mutant were normalized to total 21U-RNAs for comparison. An enlarged figure was shown for ALG-3/4 26G comparison. **F)** Box-and-whisker plot of small RNA enrichment or depletion in nuclear extract vs. input extract. For each gene, a ratio of small RNAs in the nuclear extract to those in the input was obtained, and then a box and whisker plot was generated using the ratios for the indicated number of genes in each small RNA group. * indicates significant changes from ratio 1 using Wilcoxon signed-rank test.

**Figure S5.**
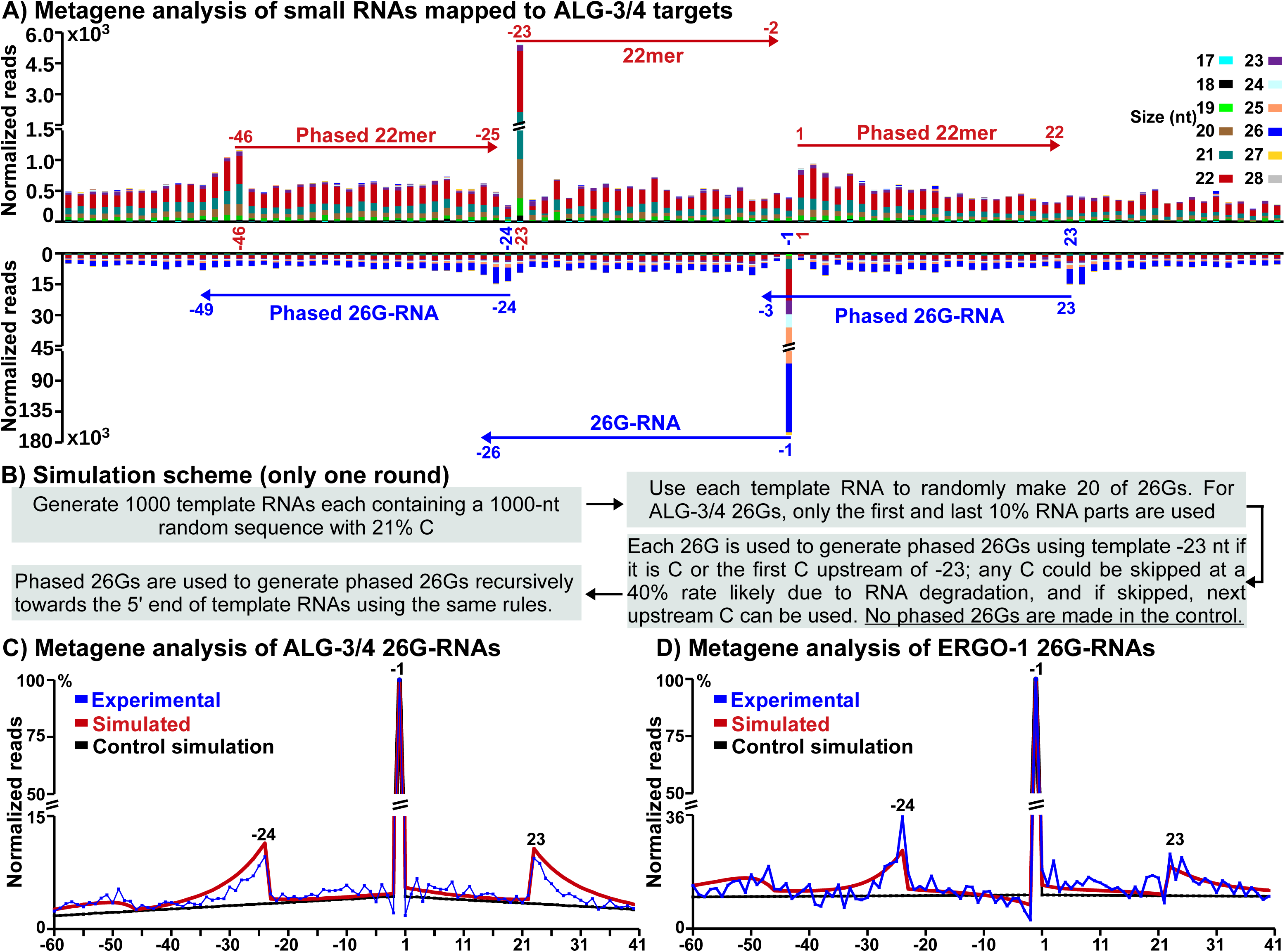
Metagene analysis of small RNAs around 26G-RNAs. Related to Figure 5 and see Supplementary Discussion. **A)** distribution of small RNAs in the ALG-3/4 pathway in L4-stage hermaphrodites: each 26G-RNA is represented by its very 5’ G, defined as -1 using a template mRNA C nt; small RNA reads (’Y’ axis) of various sizes mapped to each position of the -60 (59 nts upstream of -1) to 40 (40 nts downstream of -1) region (’X’ axis) are obtained for each 26G-RNA after normalization to total 21U-RNA reads, and accumulated for all 26G-RNAs. Top panel represents sense RNAs derived from mRNAs and bottom represents antisense RNAs made by RdRPs. **B)** A mathematical simulation scheme for modeling the distribution of 26G-RNAs with all parameters based on the experimental data but a 40% failure rate of each template C usage empirically determined for obtaining the best fit. **C-D)** distribution of 26G-RNA reads around specified 26G-RNA loci in the ALG-3/4 pathway based on the experimental and simulation data using the same method as in A); ‘Y’ axis represents the ratio of the 26G-RNA reads at each position to those at mRNA -1; the simulation control uses the same parameters but does not allow for generating phased 26G-RNAs.

**Figure S6.**
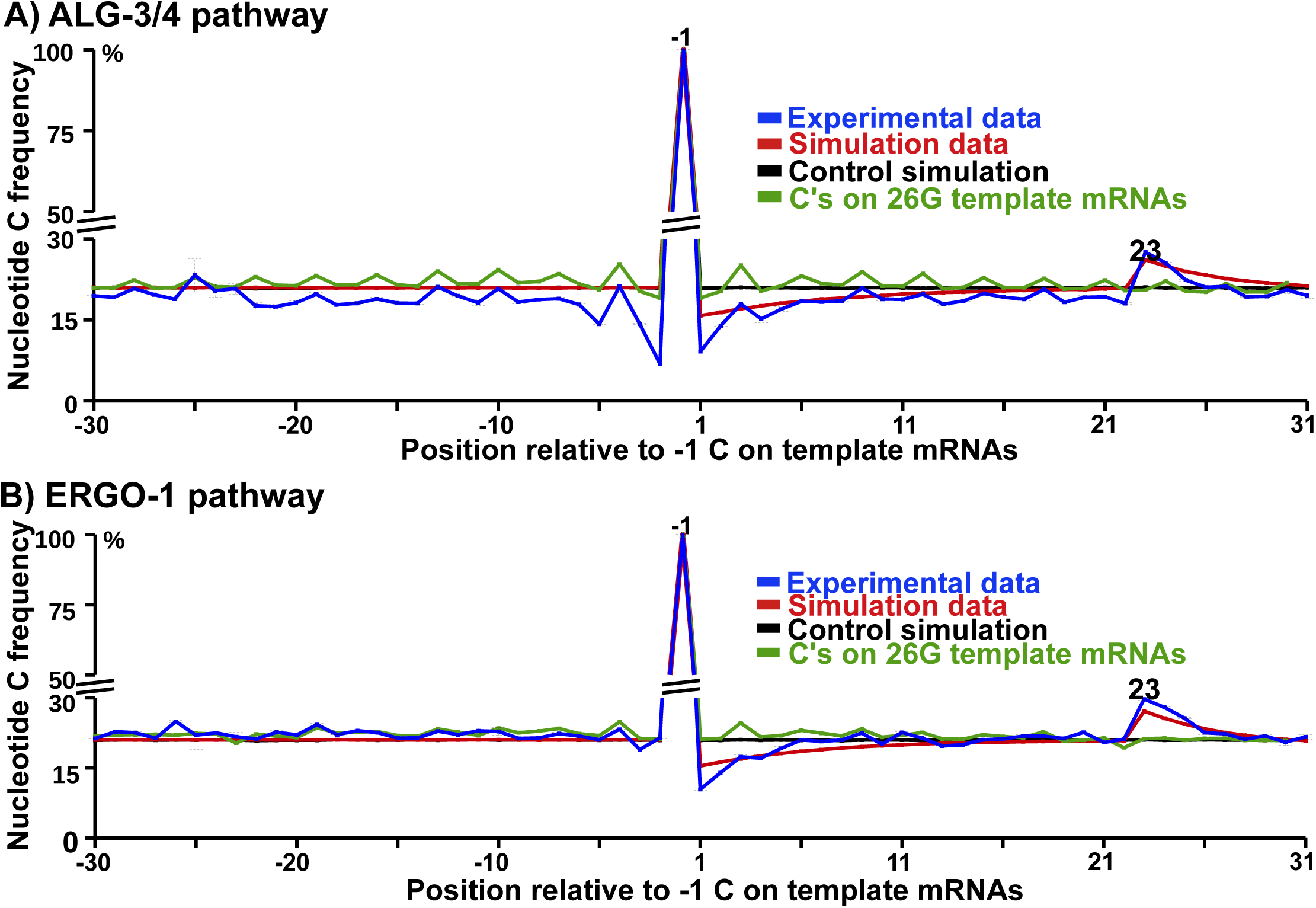
Metagene analysis of 26G-RNA distribution. Related to Figure 6 and see Supplementary Discussion. **A-B)** Analysis of C nt distribution around -1 C on template RNAs. All the -1 C encoding the first nt of 26G-RNAs in the ALG-3/4 (**A**) and ERGO-1 (**B**) pathways are selected from the experimental (blue) and simulation data (red and black), and as a genome sequence control, all C’s on template mRNAs are selected (green). The positions of these C’s are defined as -1, and the nt C frequency at each upstream (-1 to -30) and downstream (1 to 30) position was obtained using the selected loci. Unlike those metagene analyses in Figure 5 and S5, each C locus bears a weight of 1 instead of the RNA read number. The simulation data and control are the same as those used in Figure S5. Dotted lines represent one-standard-error bars in the experimental data, and those bars in the simulation data are too small to draw. On the left part of the figures, the black lines are hidden underneath the read lines.

**Table S1.** PIR-1-interacting proteins.

| **PIR-1 Interactors identified in PIR-1::GFP IP Using Young Adult Worms** |  |  |  |
| --- | --- | --- | --- |
| **Protein** | **Amino Acid Number** | **Spectral Counts<sup>a</sup>** | **Protein Coverage** |
| PIR-1 | 233 | 142 | 35.9% |
| DCR-1 | 1845 | 118 | 22.8% |
| RRF-3 | 1780 | 54 | 17.9% |
| DRH-3 | 1119 | 50 | 24.8% |
| ERI-3 | 578 | 23 | 14.9% |
| RDE-4 | 385 | 15 | 17.1% |
| ERI-1 | 582 | 11 | 11.3% |
| ERI-5<sup>b</sup> | 531 | 9 | 8.7% |

| **PIR-1 Interactors identified in PIR-1::3xFlag IP Using Gravid Adult Worms** |  |  |  |
| --- | --- | --- | --- |
| **Protein** | **Amino Acid Number** | **Spectral Counts<sup>a</sup>** | **Protein Coverage** |
| PIR-1 | 233 | 12 | 22.3% |
| DCR-1 | 1845 | 27 | 9.0% |
| RRF-3 | 1780 | 15 | 6.1% |
| DRH-3 | 1119 | 19 | 7.1% |
| ERI-3<sup>b</sup> | 578 | 2 | 5.7% |
| RDE-4<sup>b</sup> | 385 | 8 | 15.1% |

<sup>a</sup>The number of tandem mass spectra matching peptides derived from each protein
<sup>b</sup>The spectral count is less than 10.

**Table S2.**
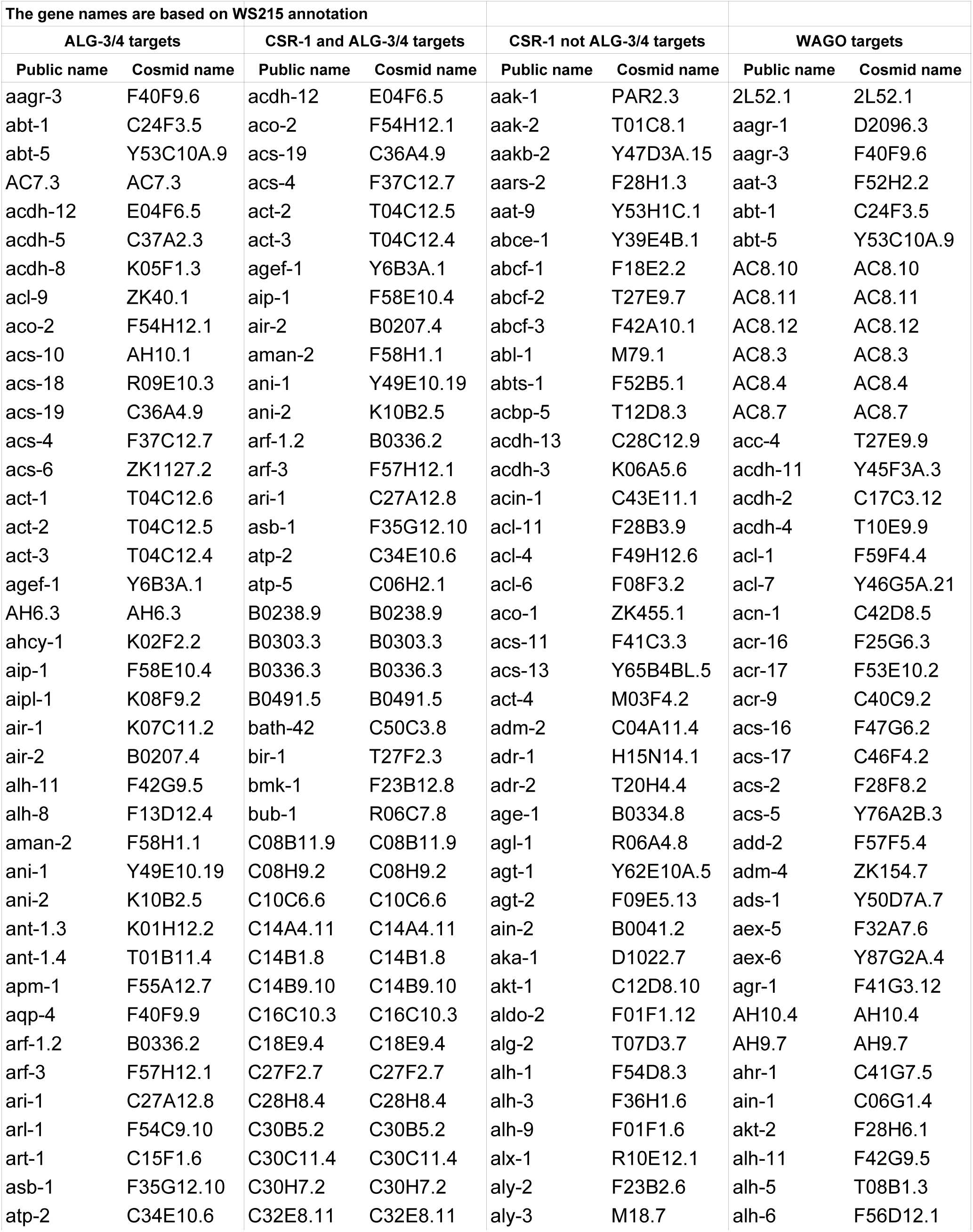
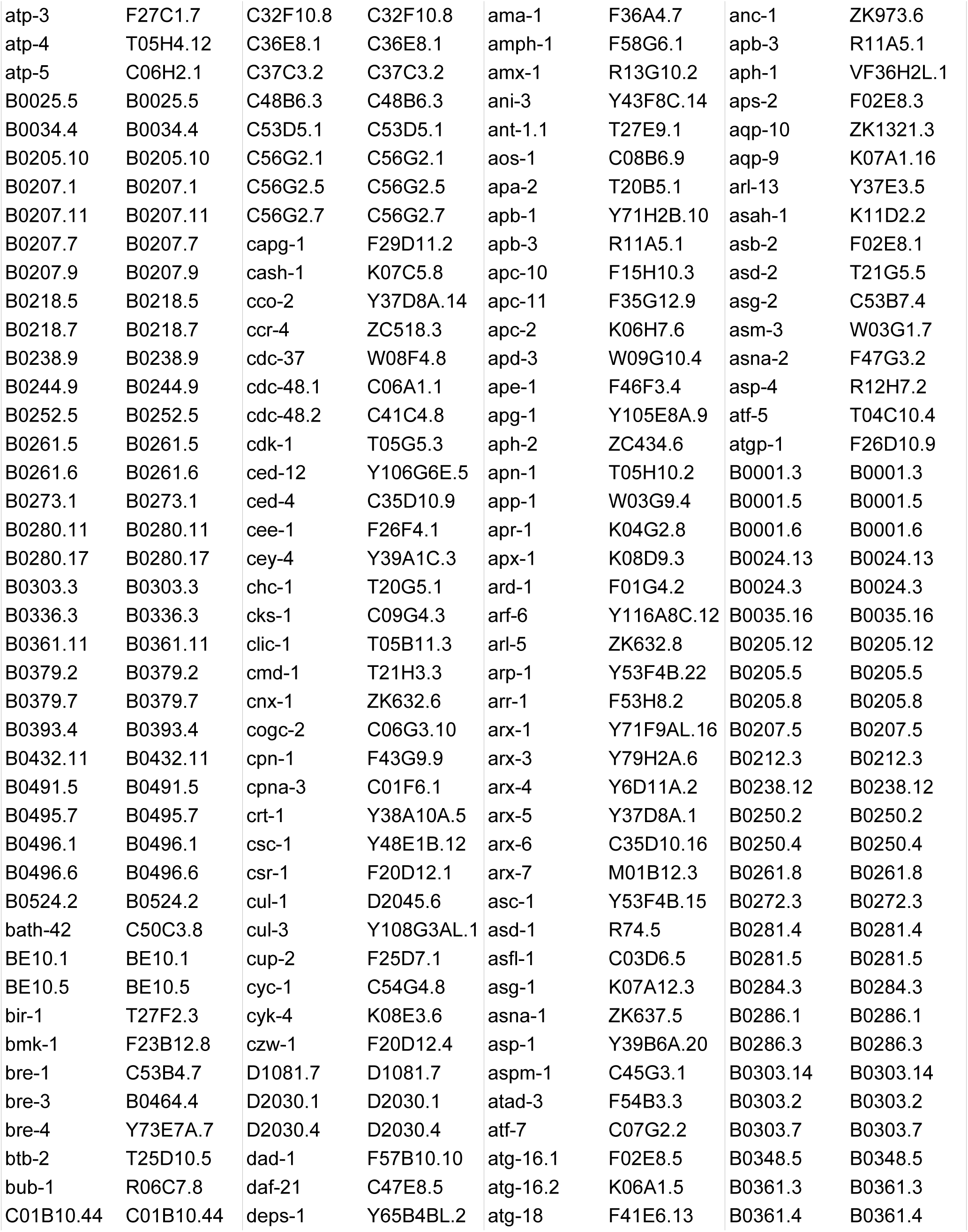
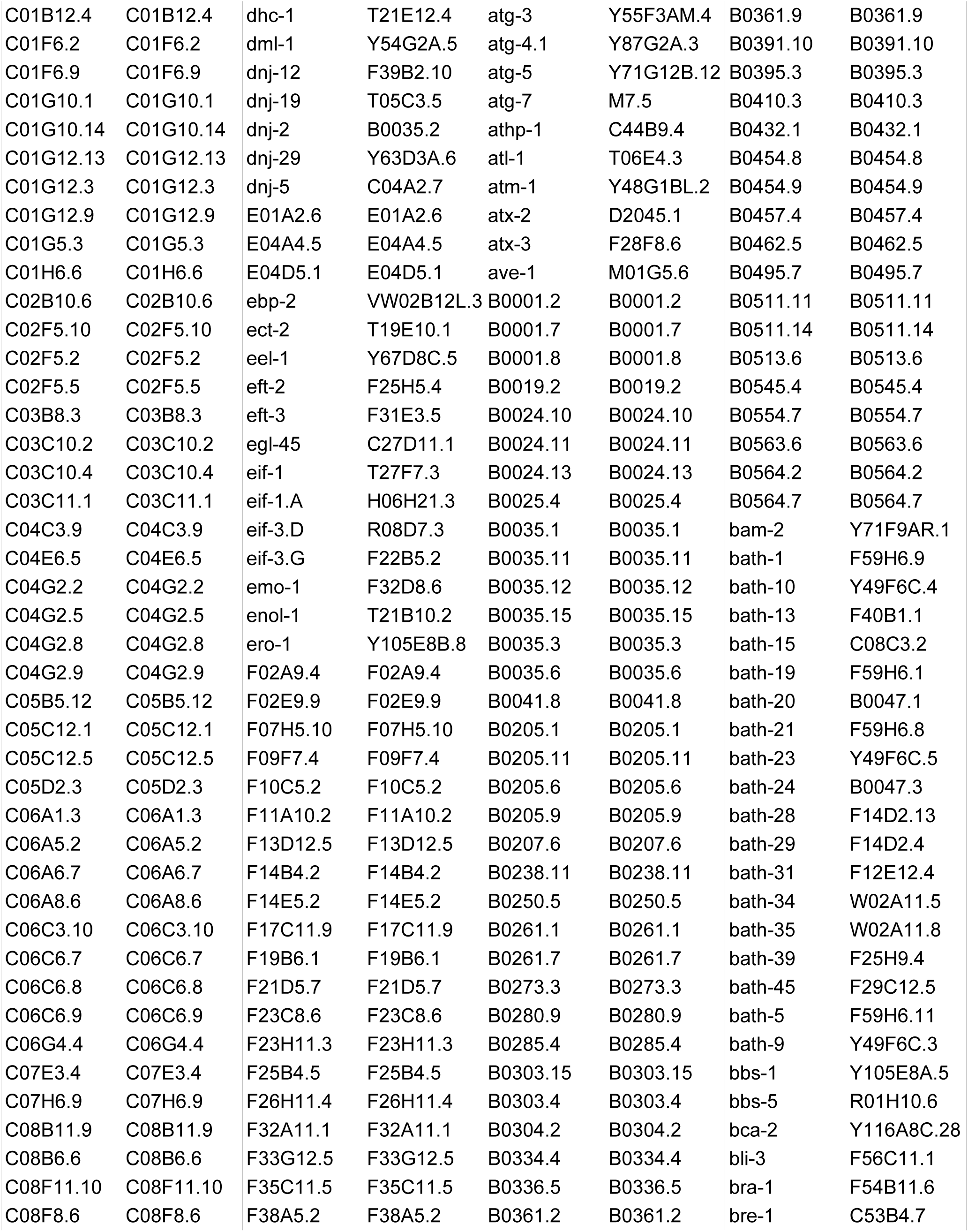
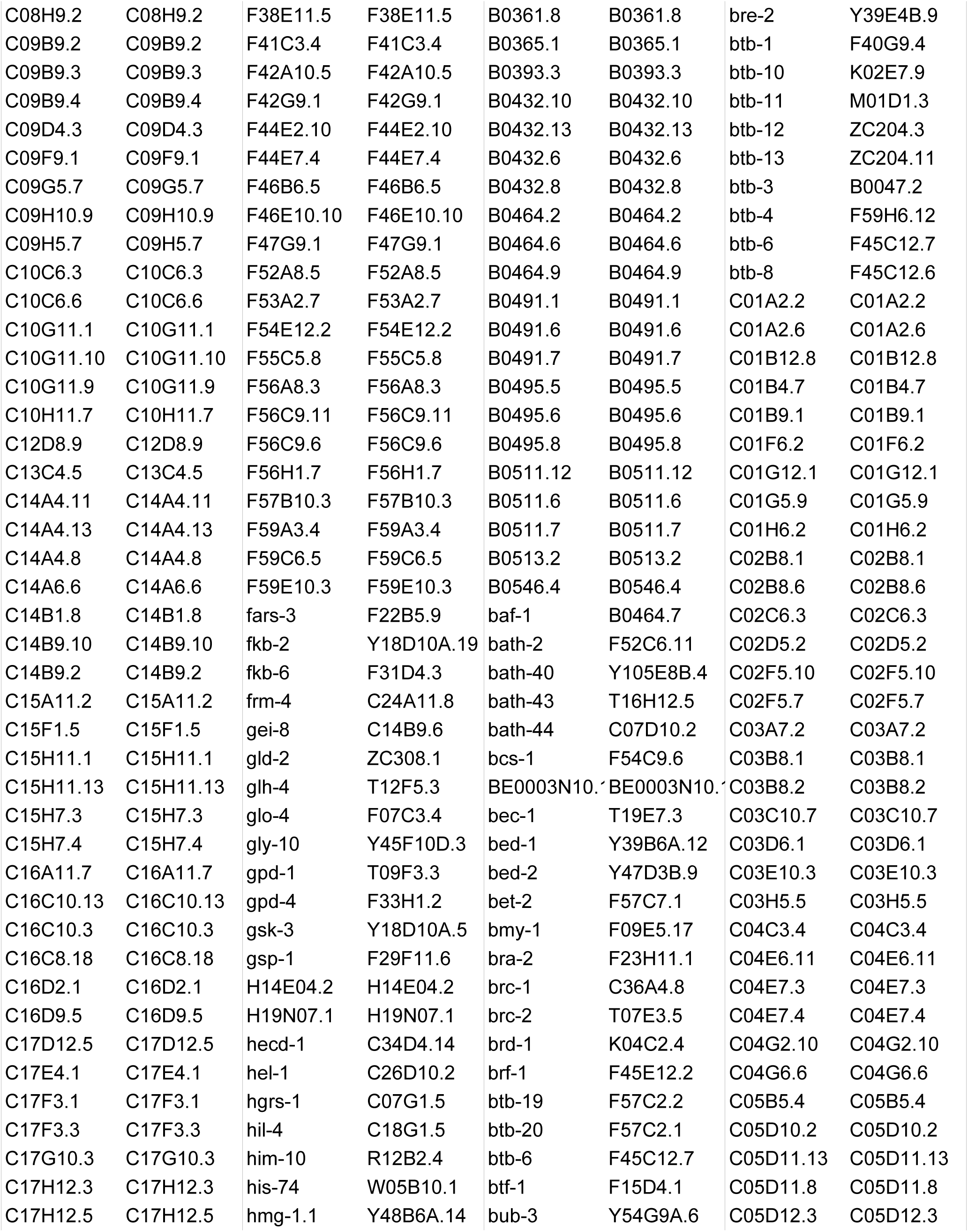
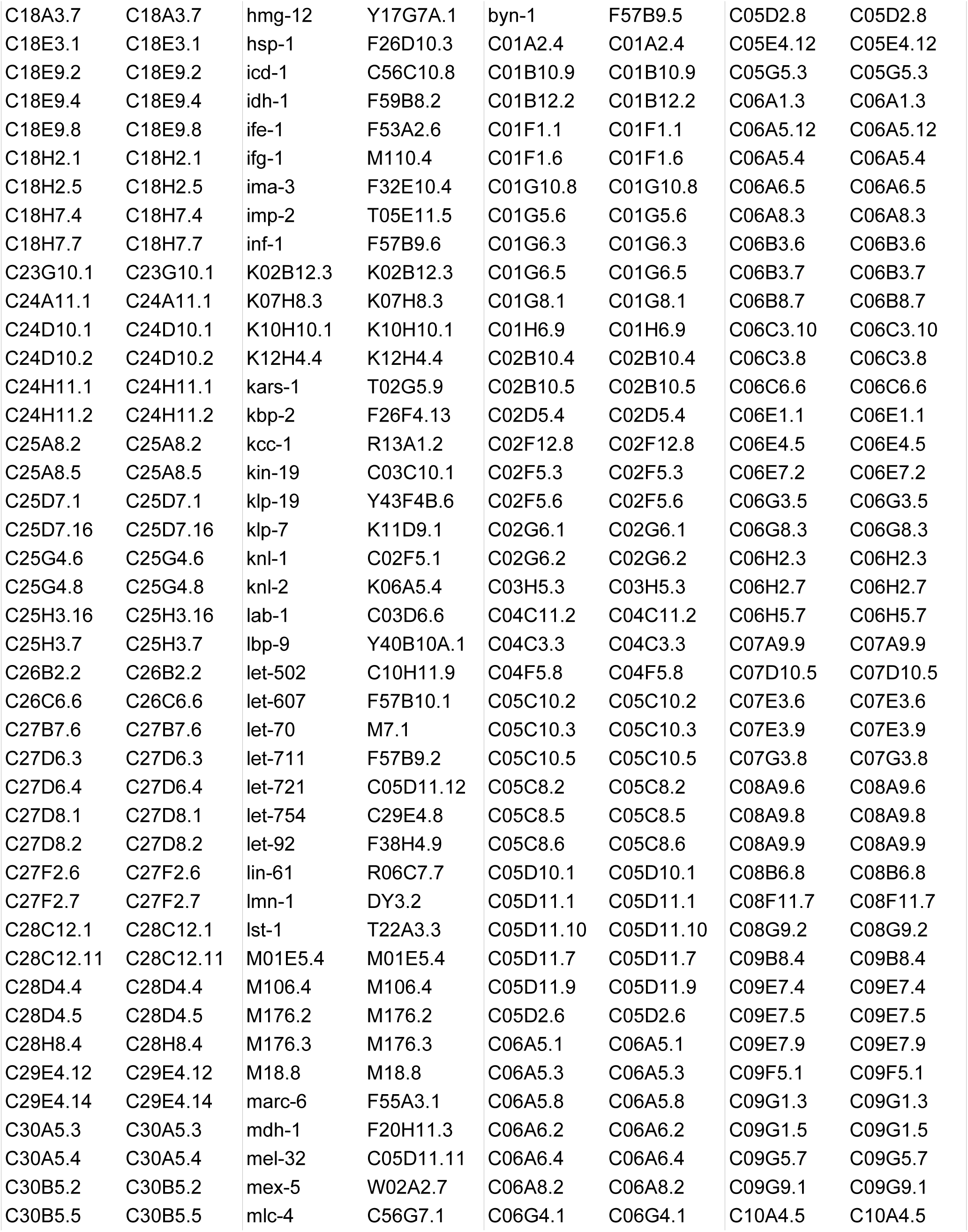
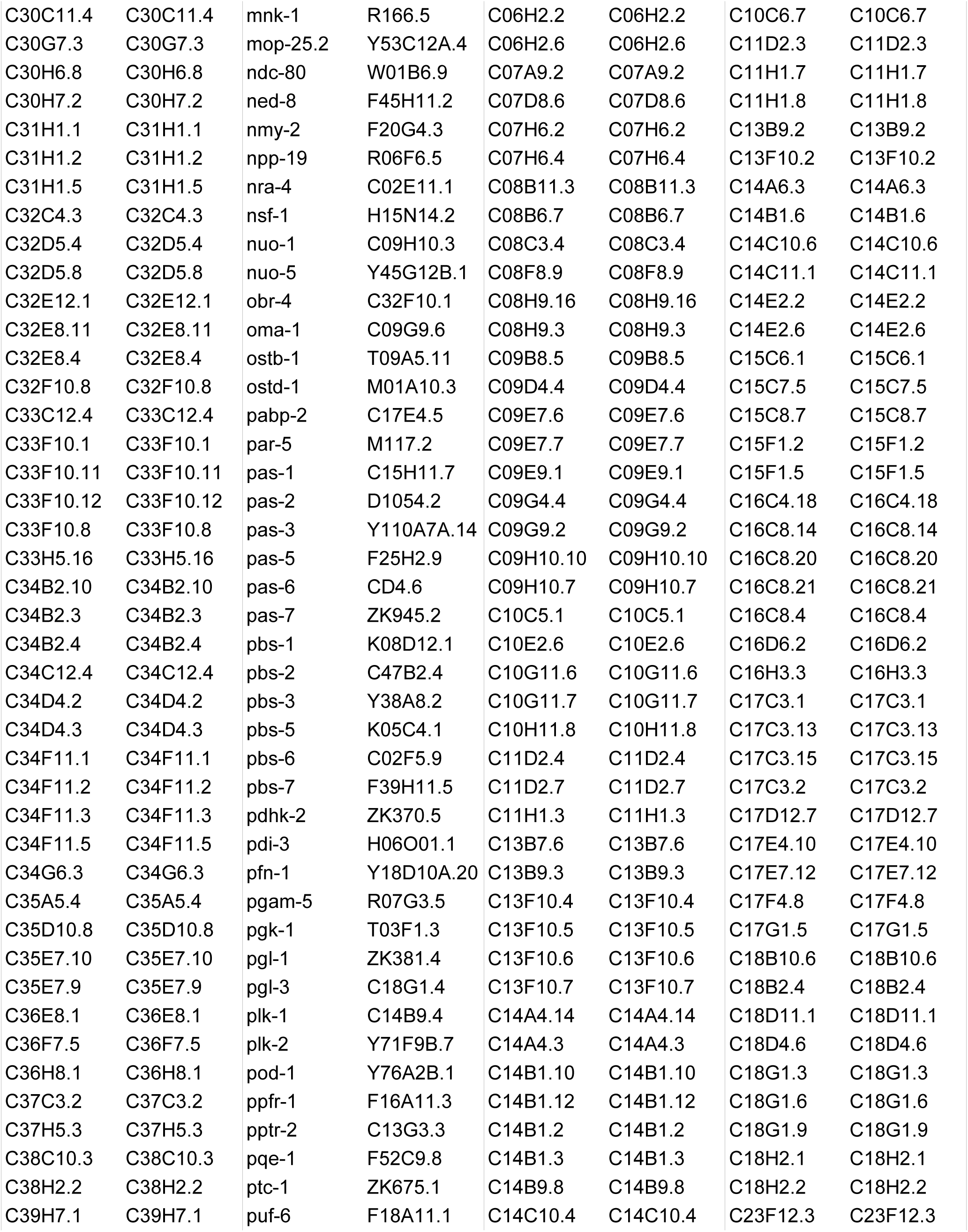
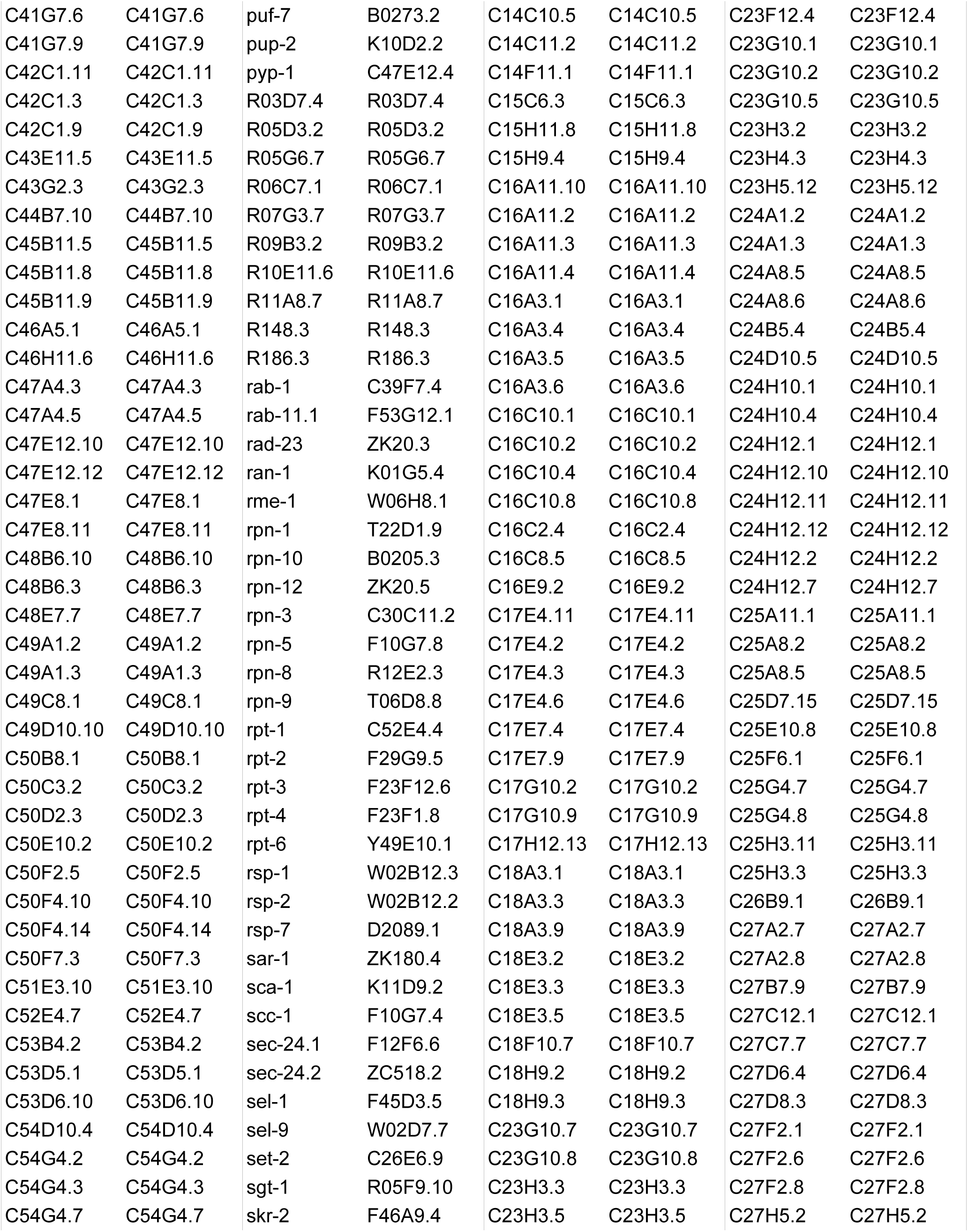
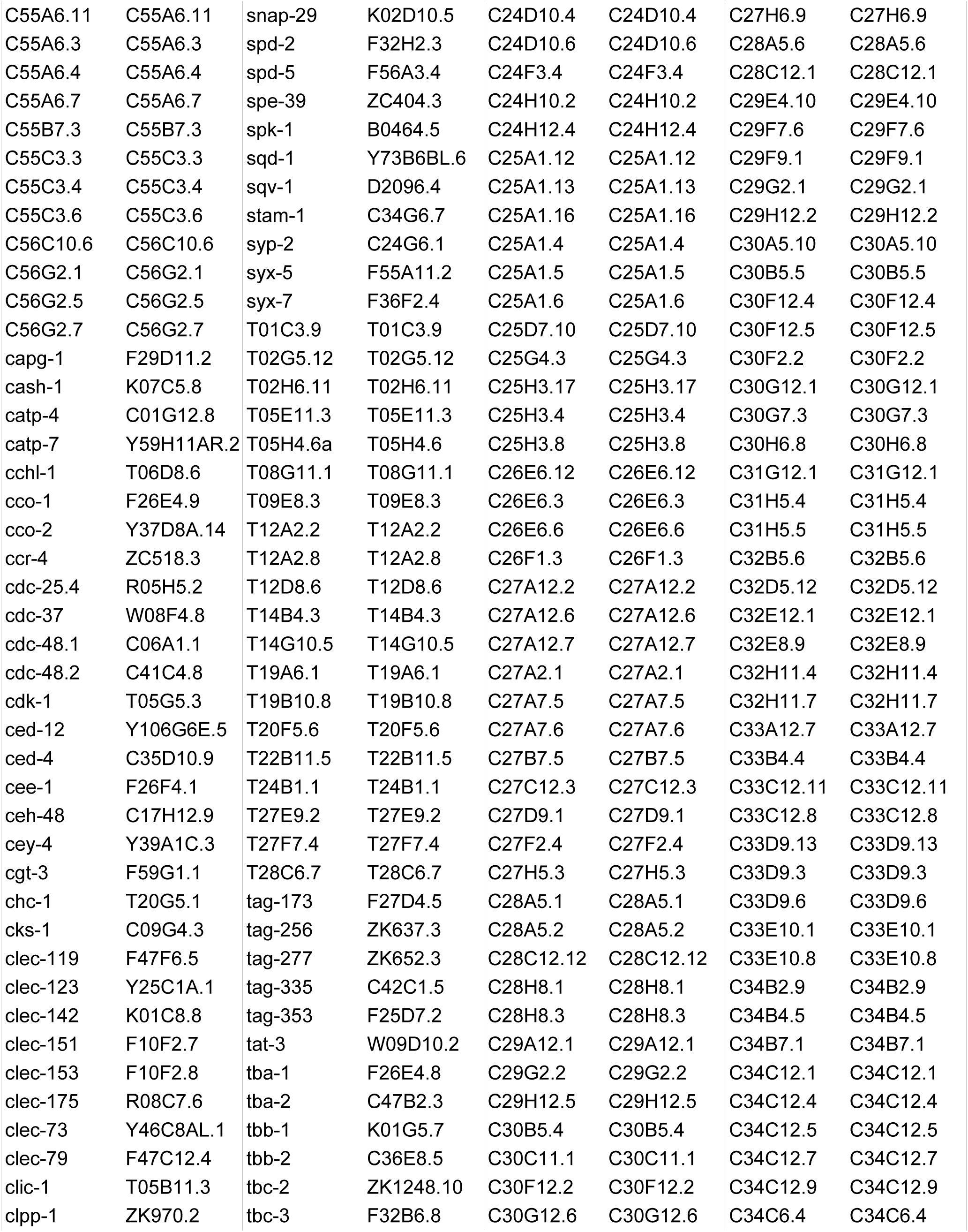
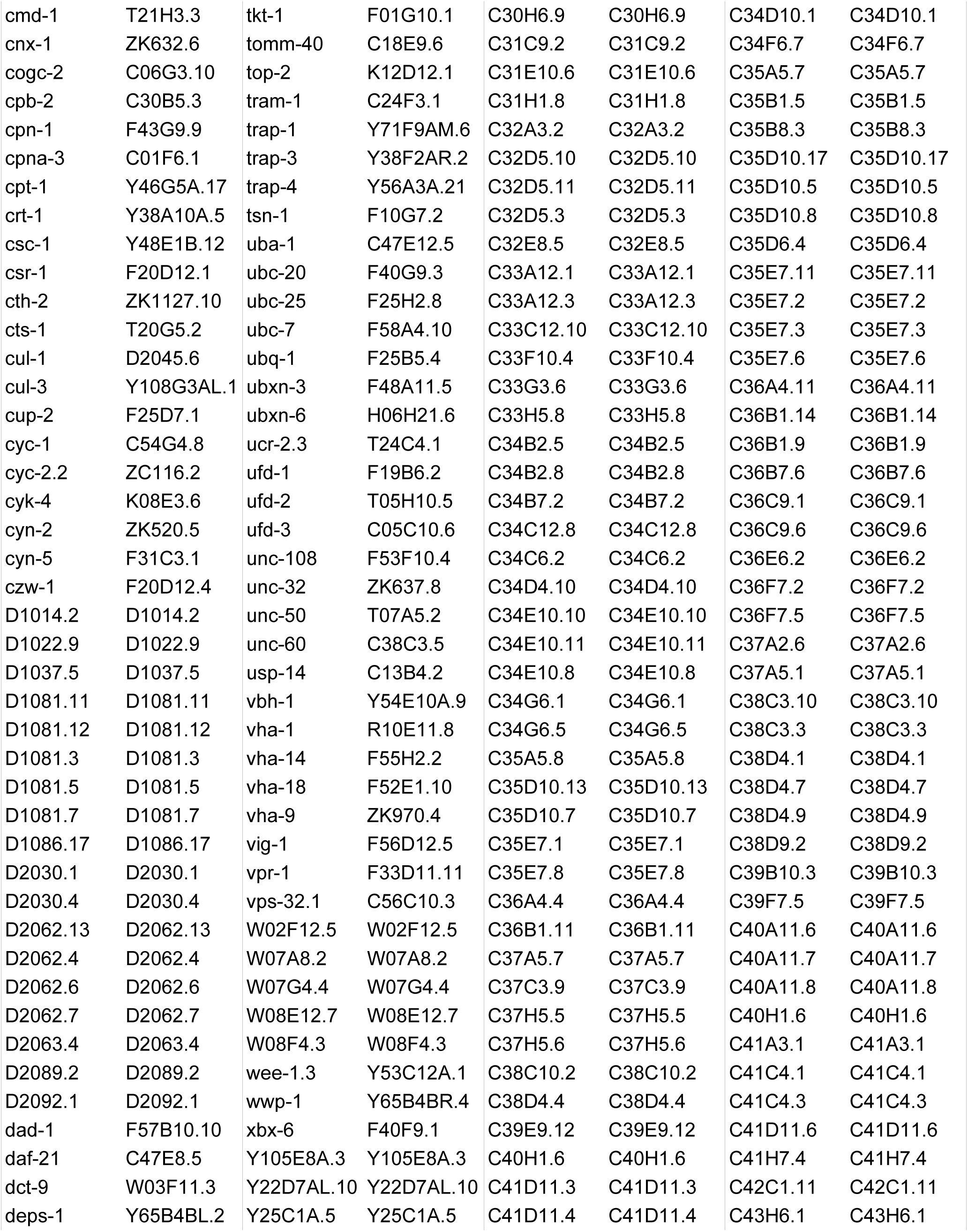
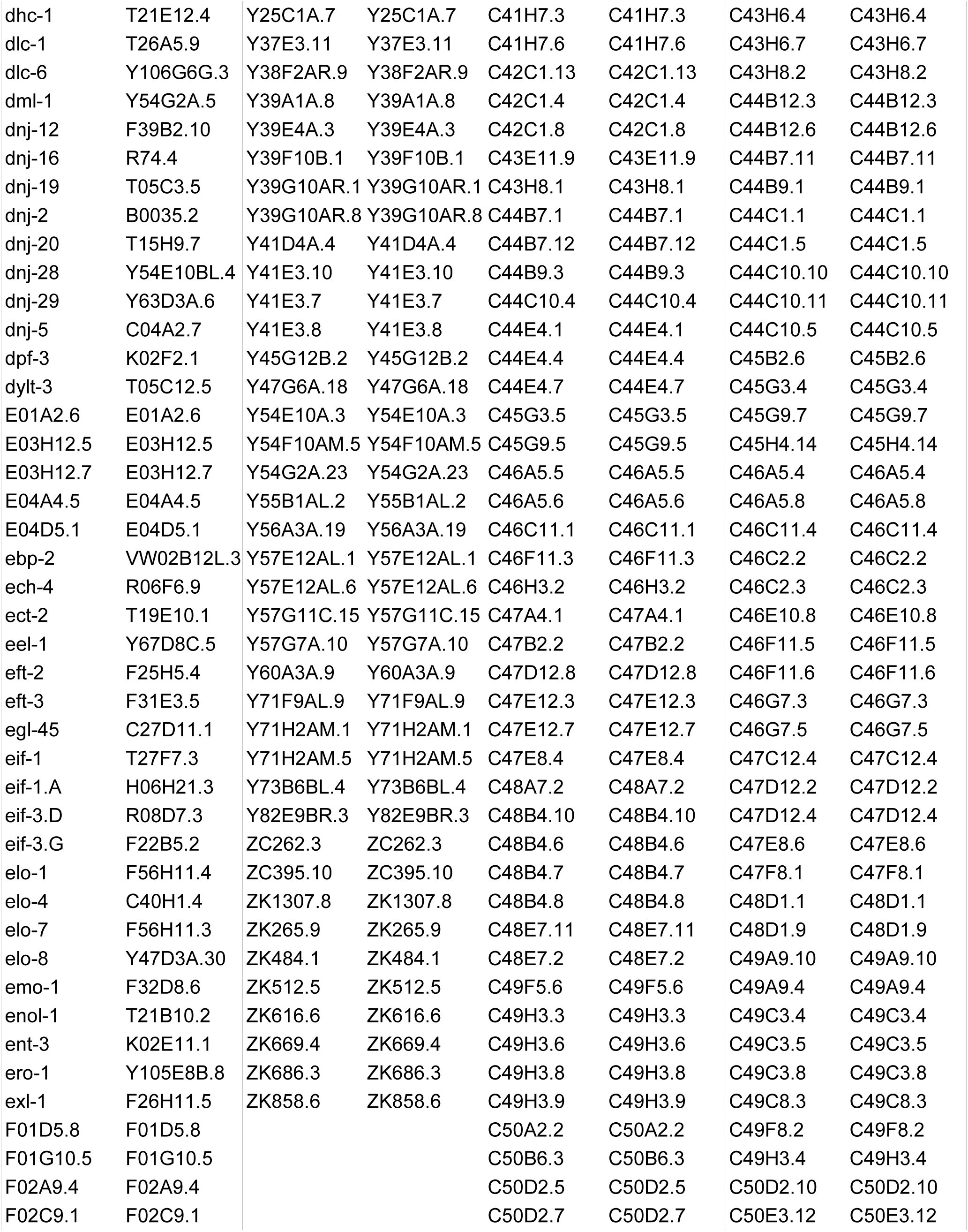
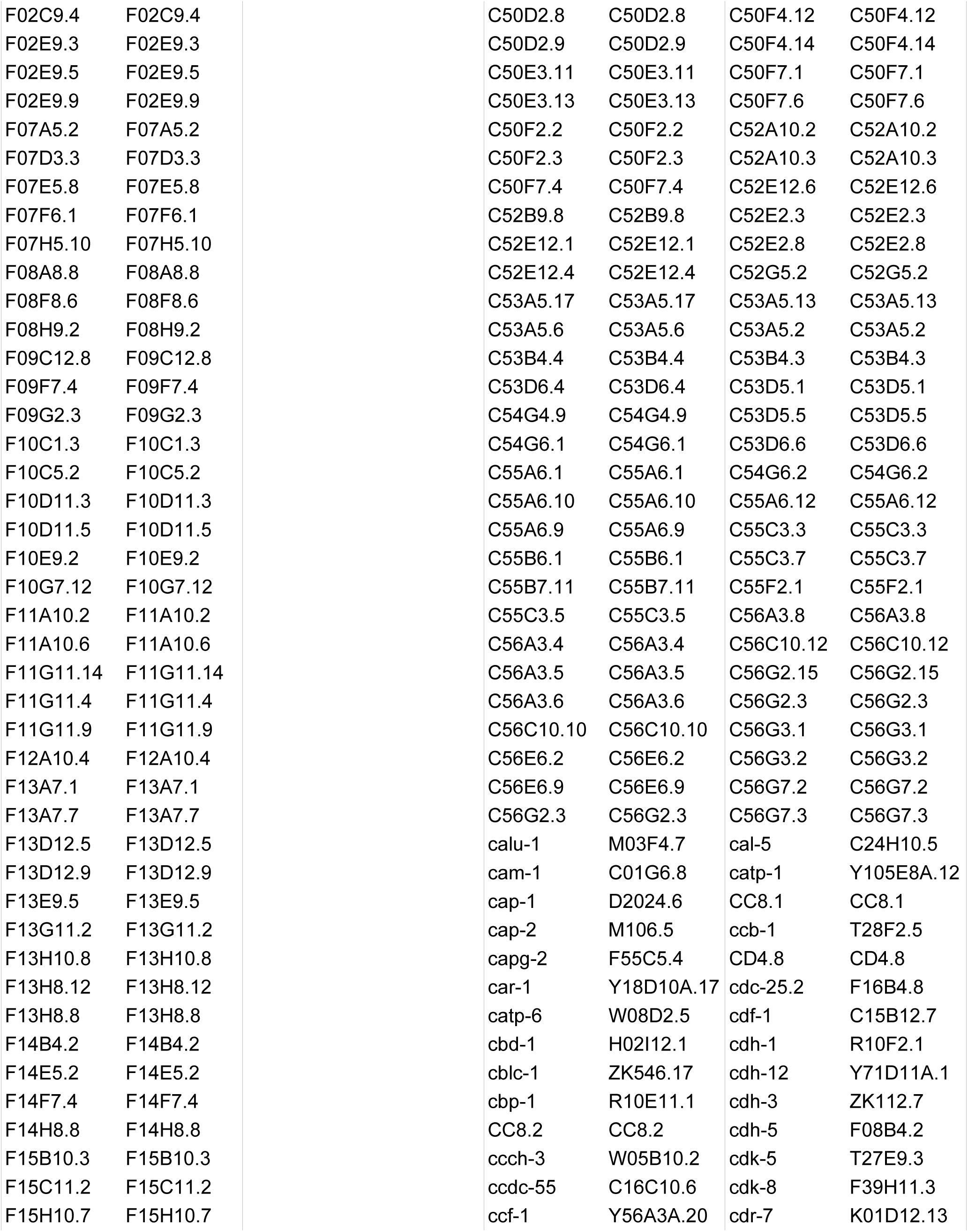
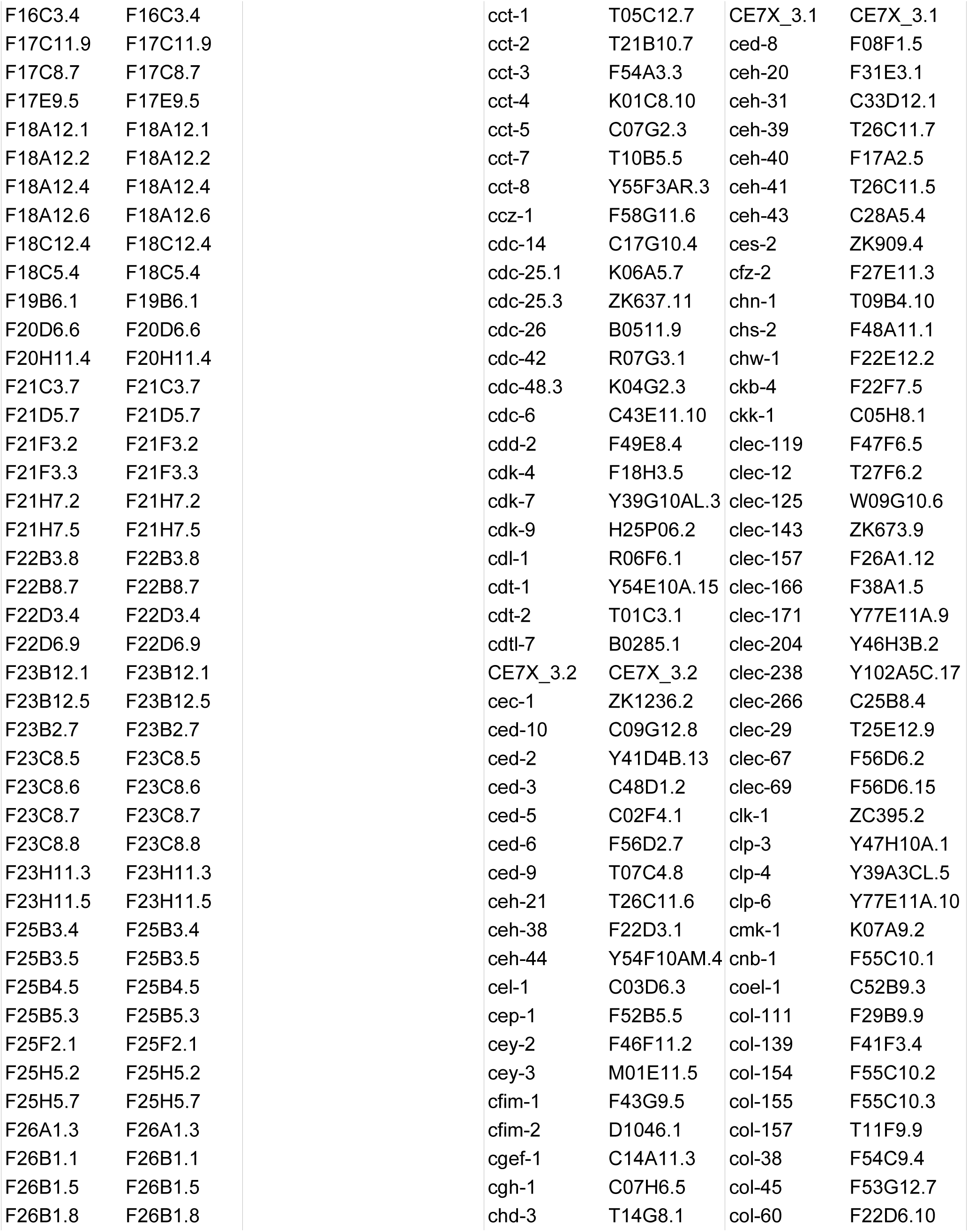
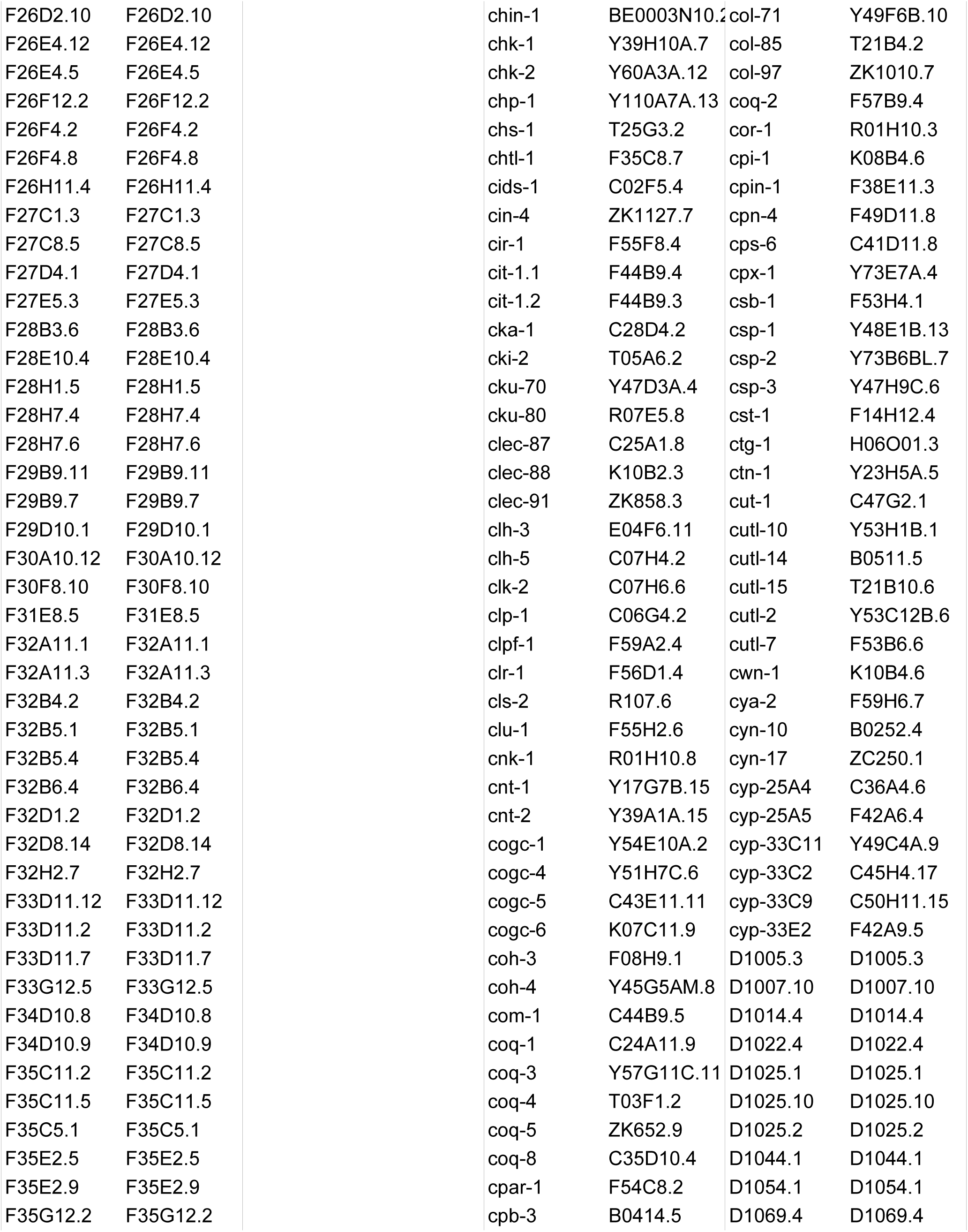
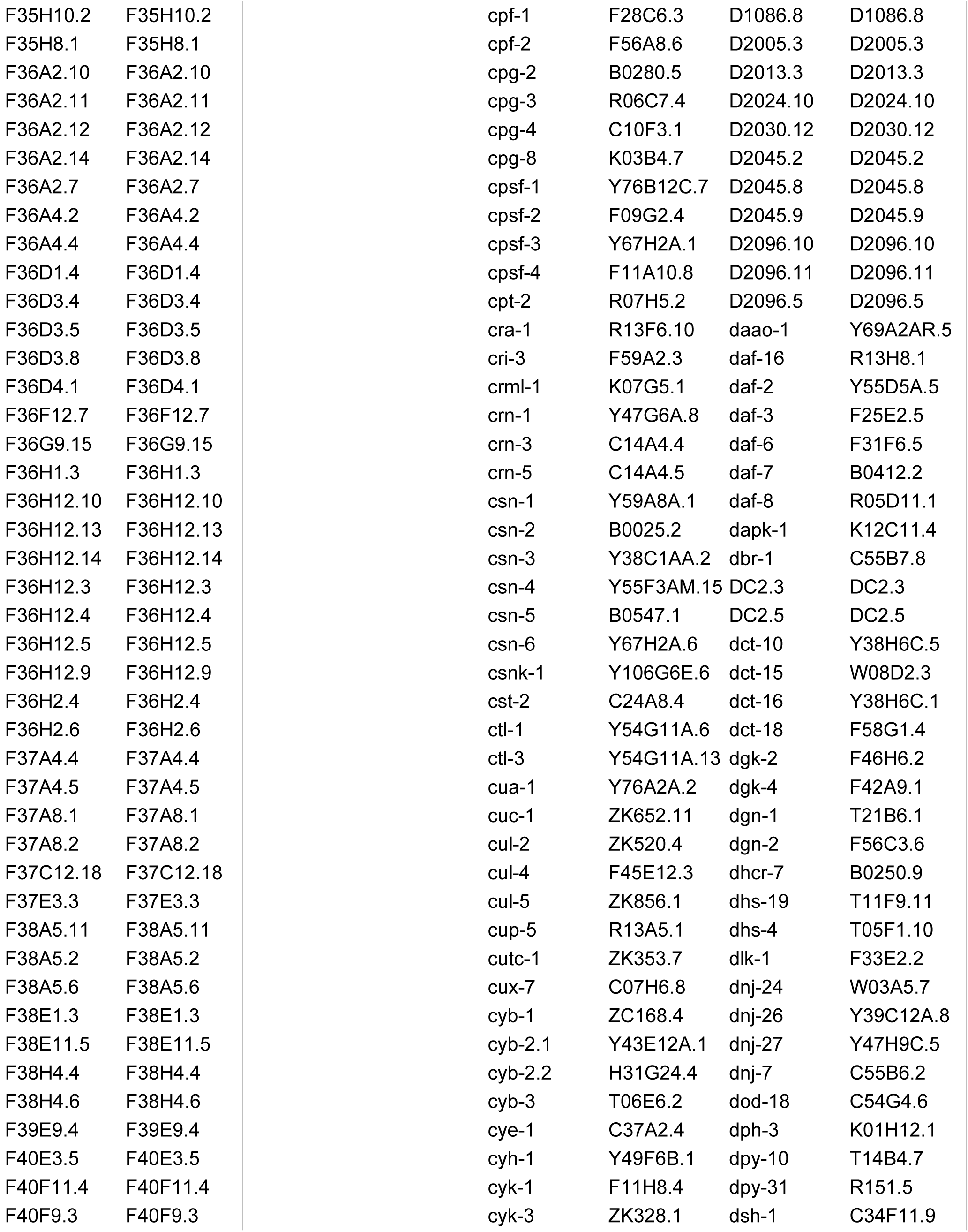
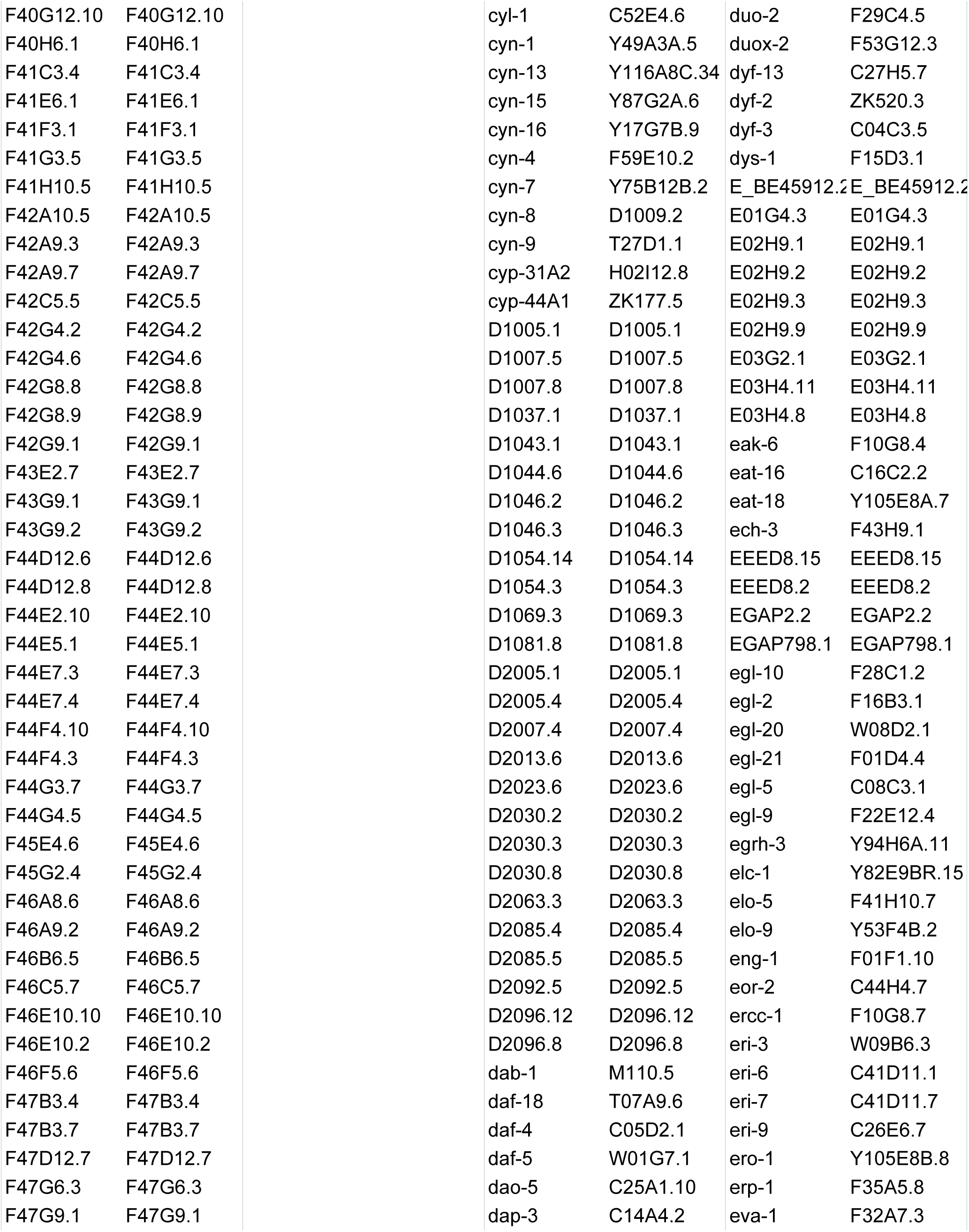
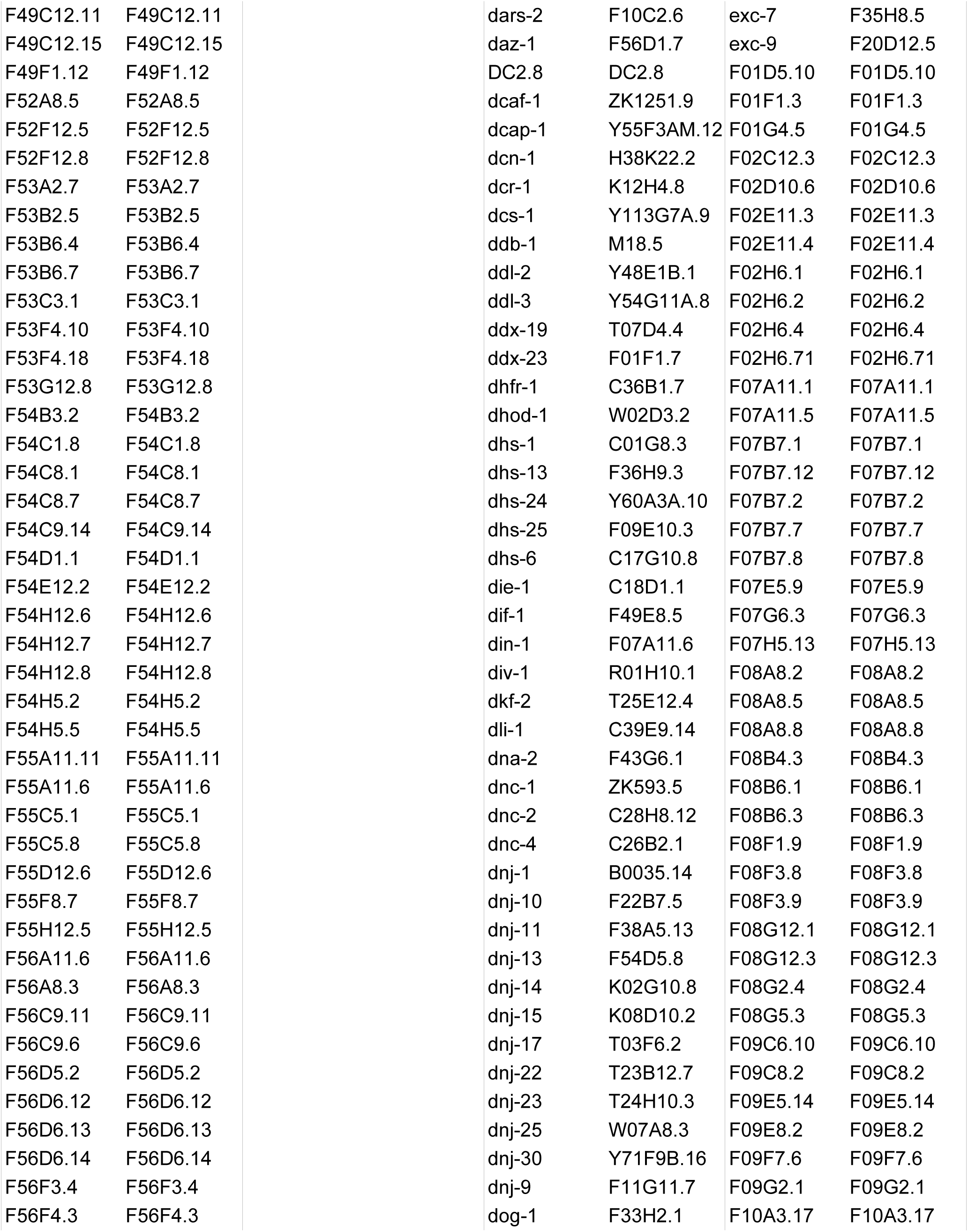
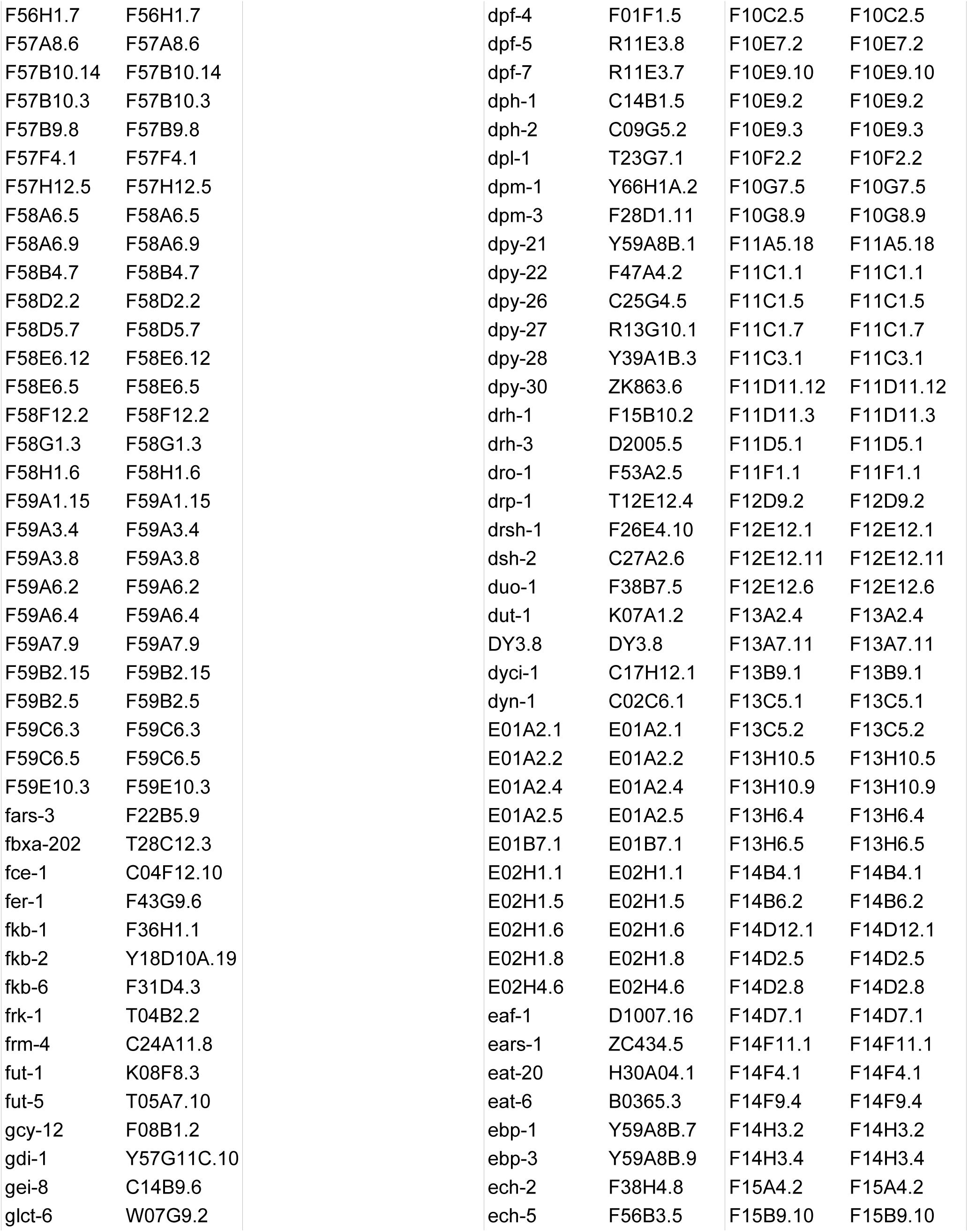
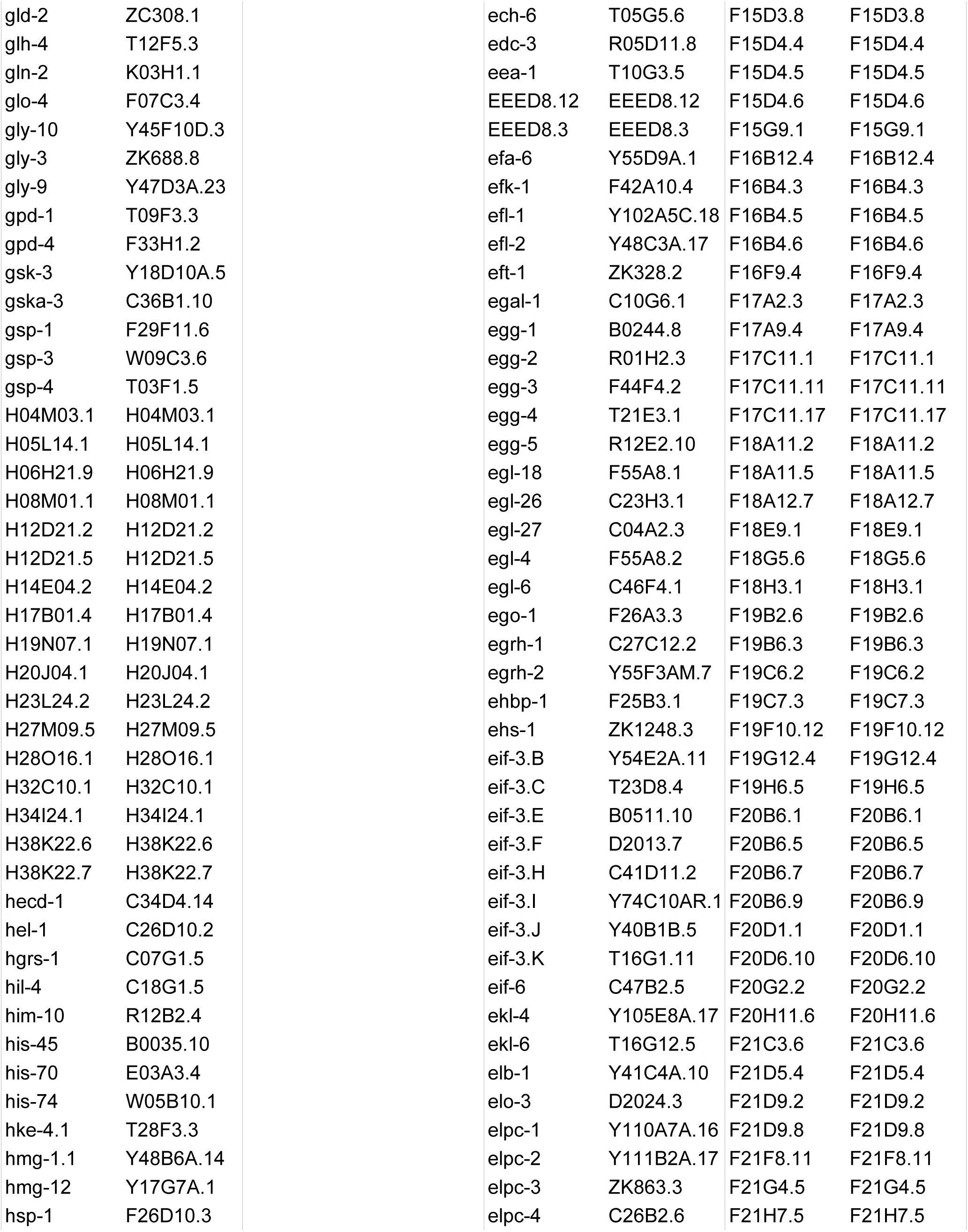
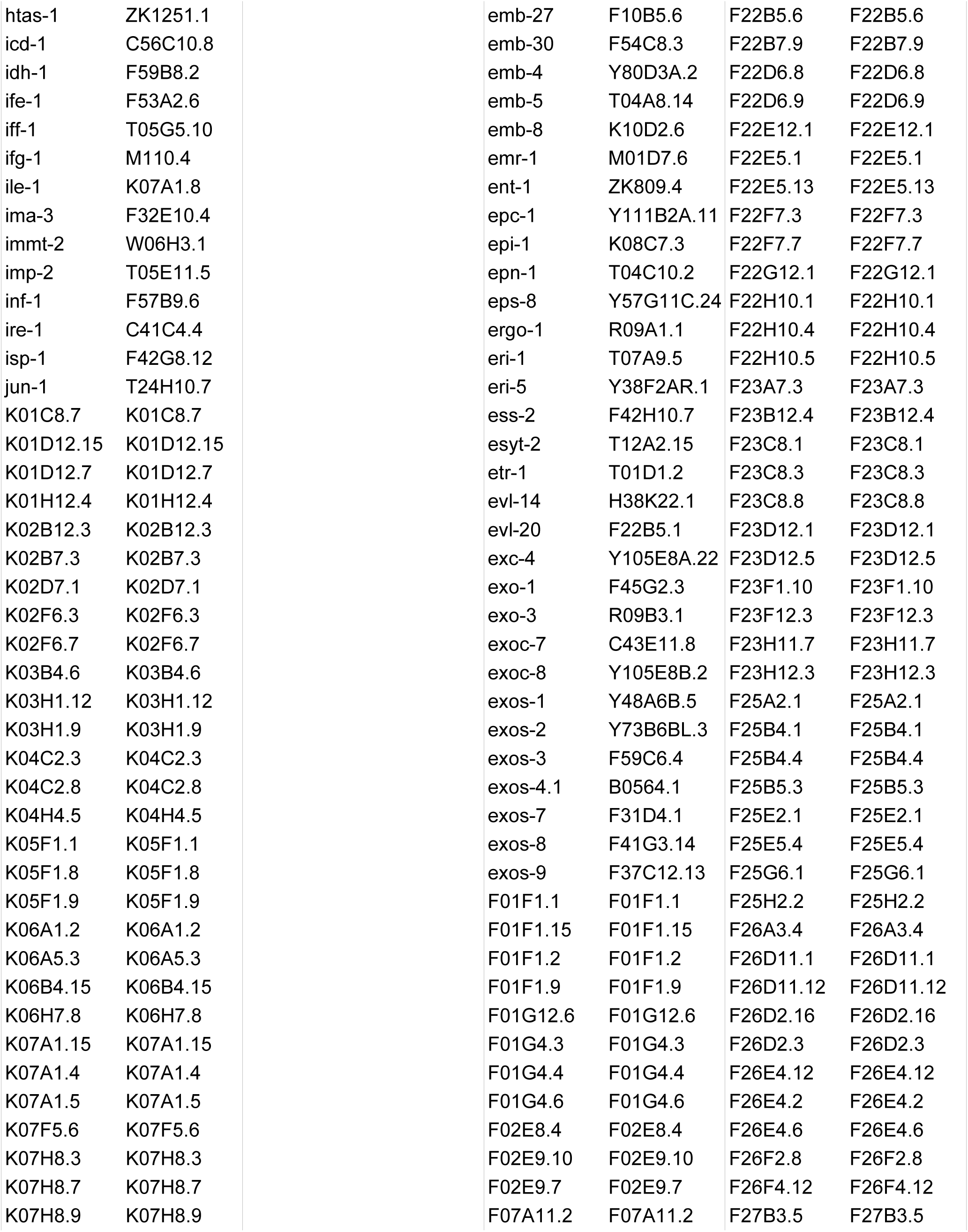
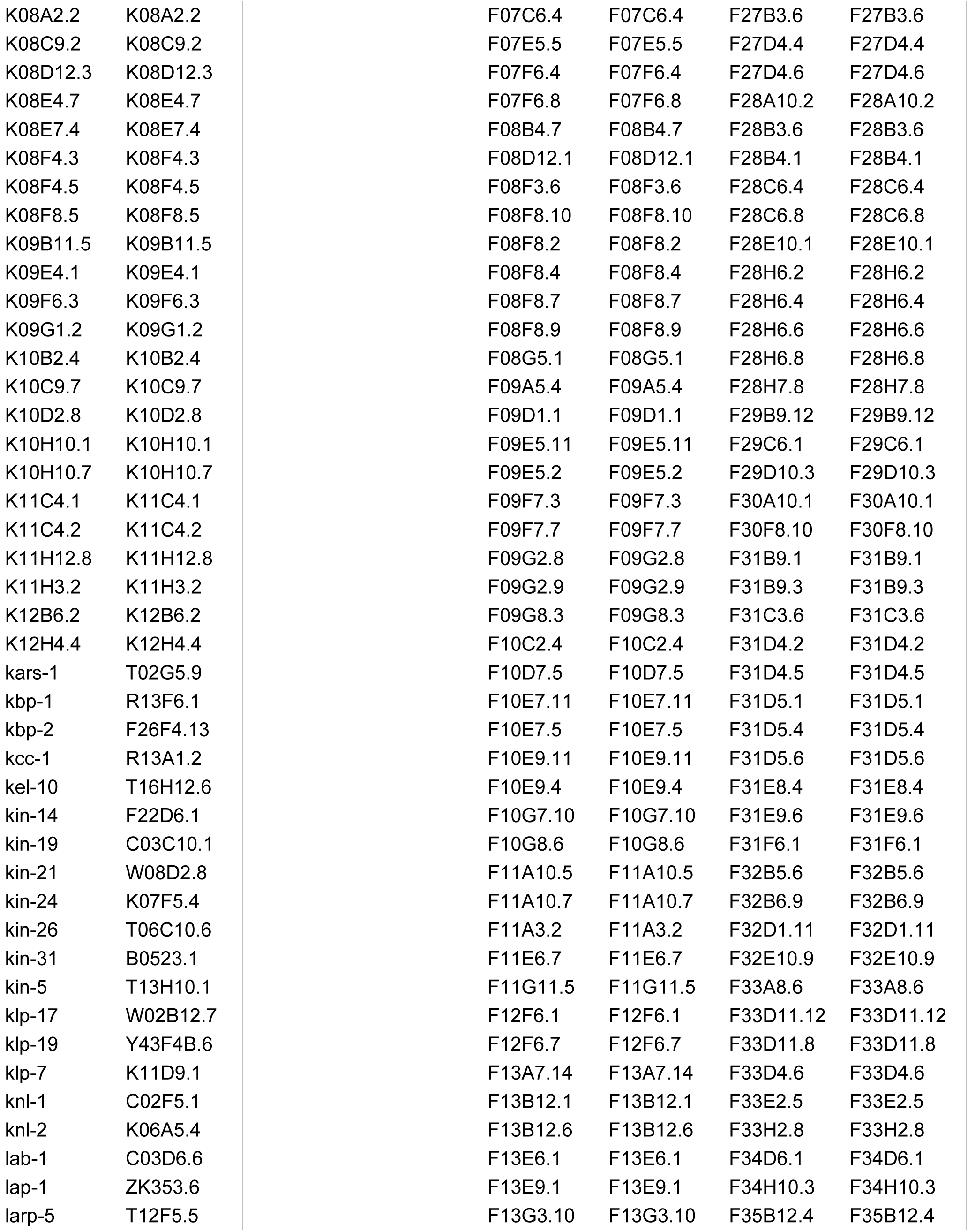
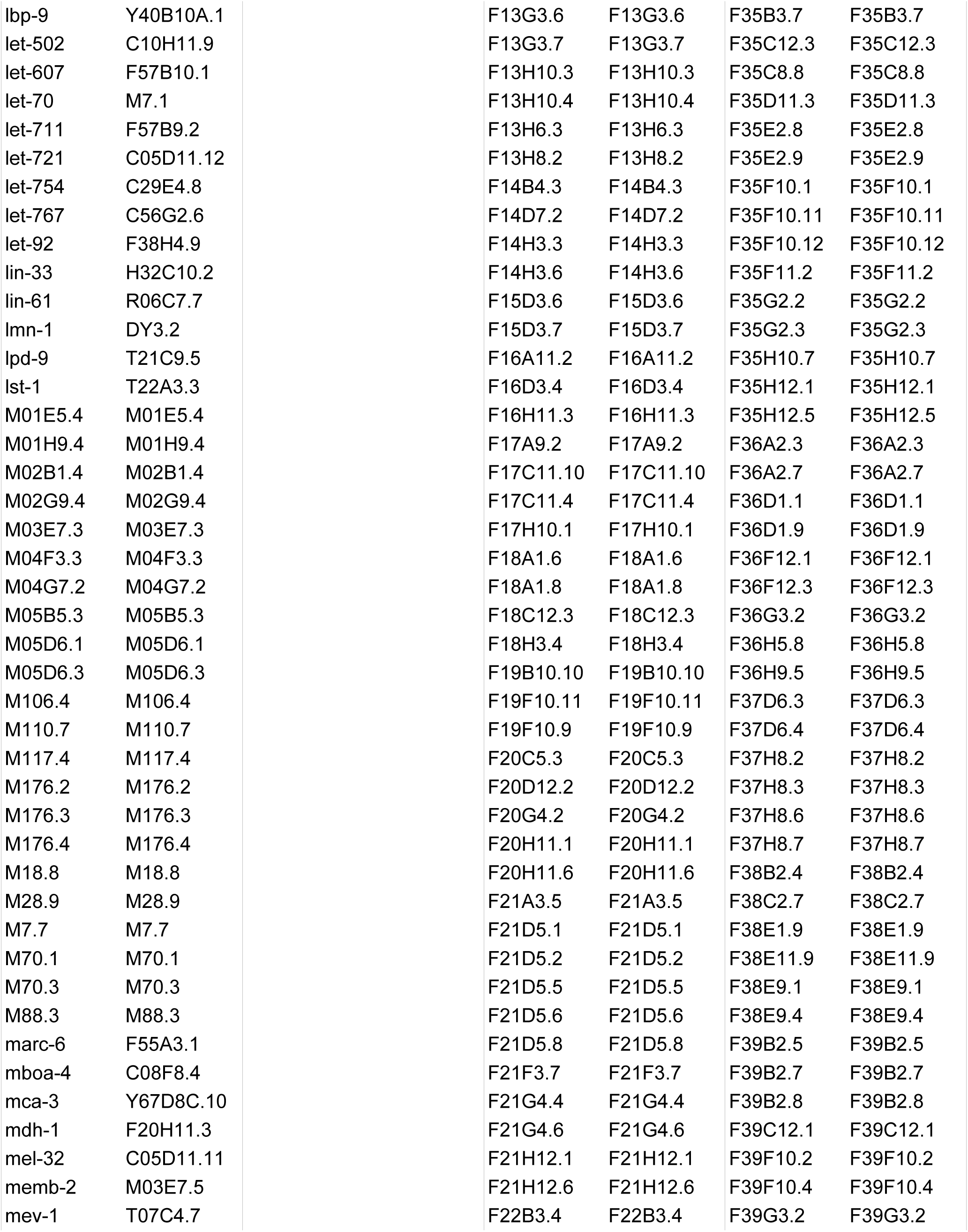
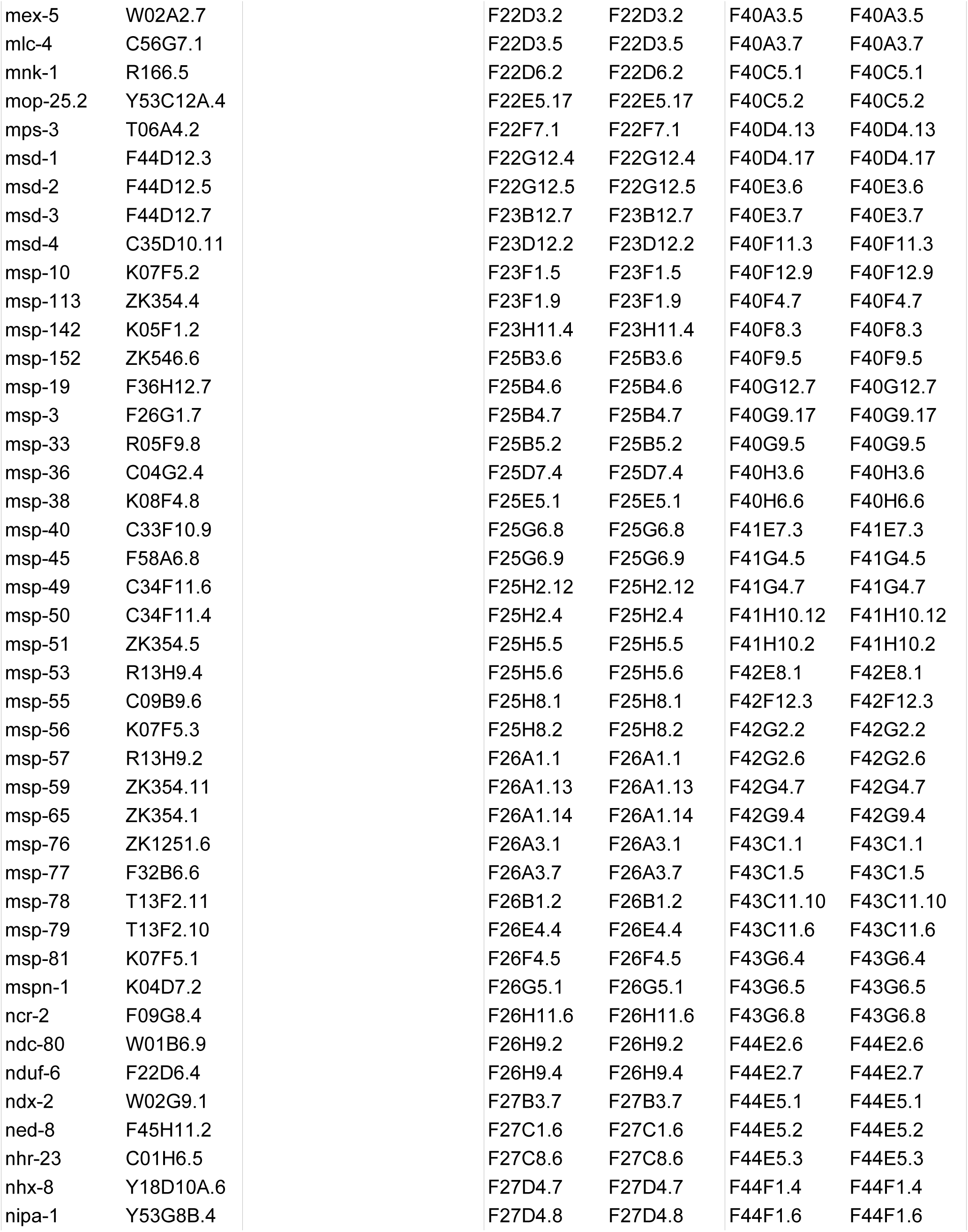
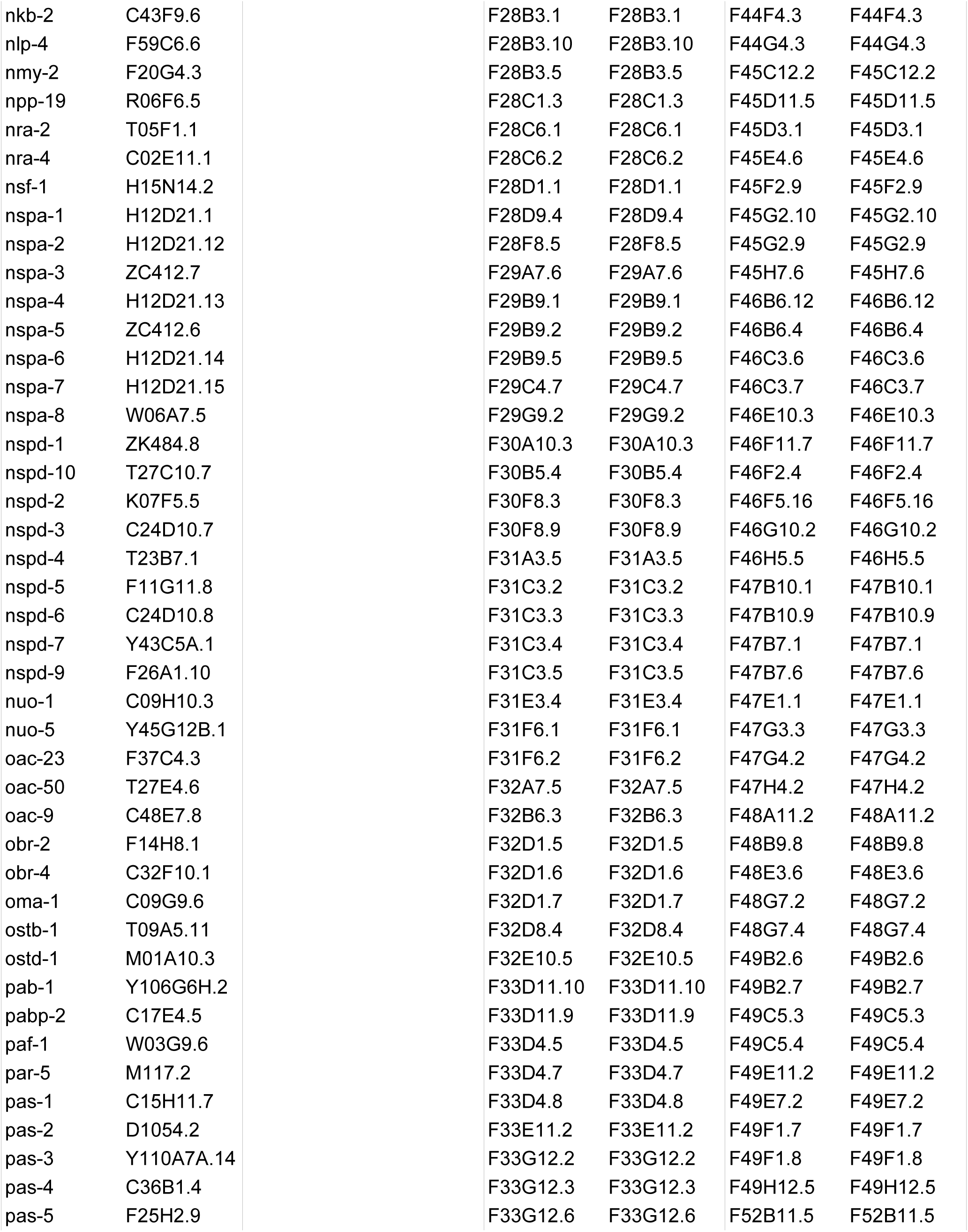
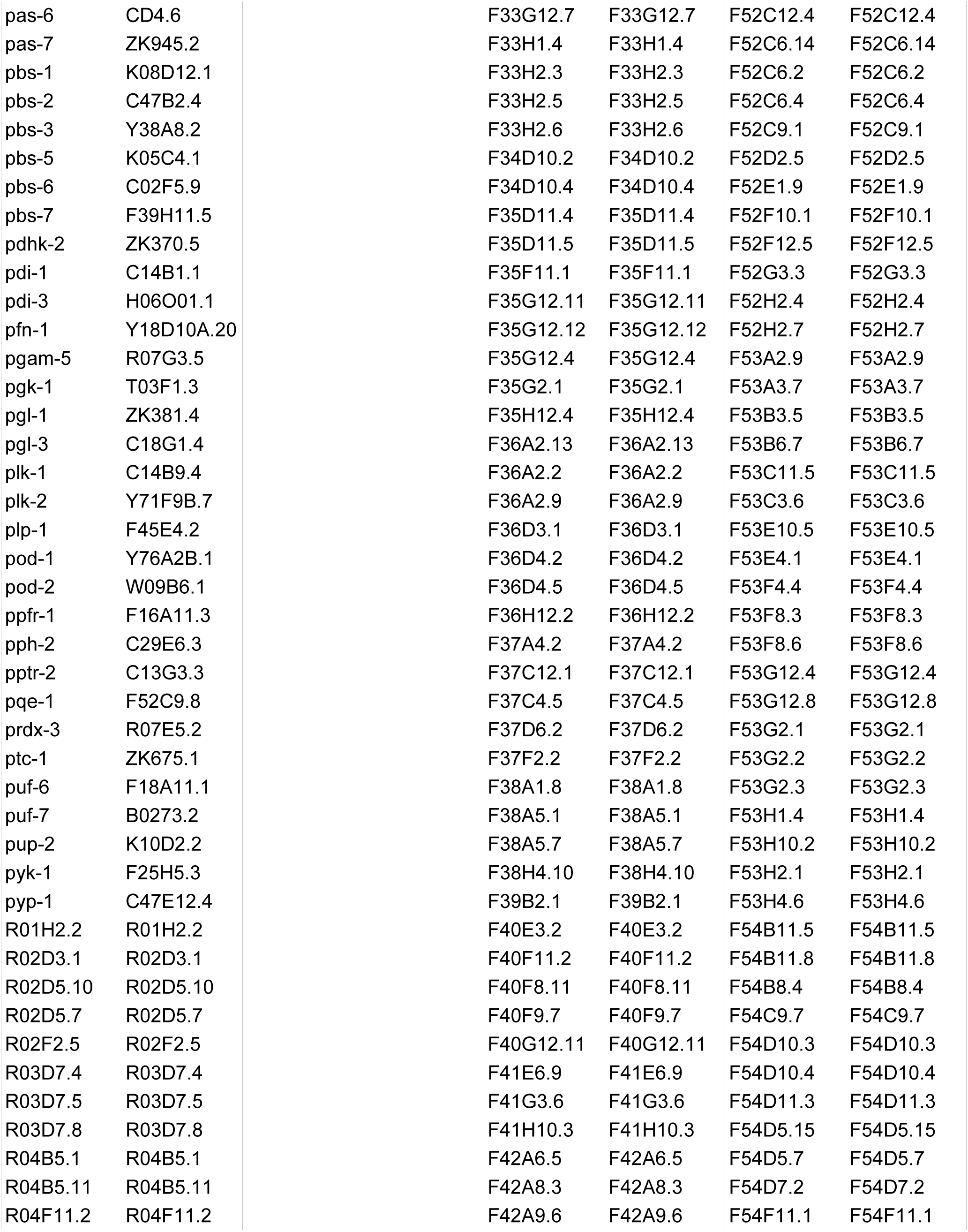
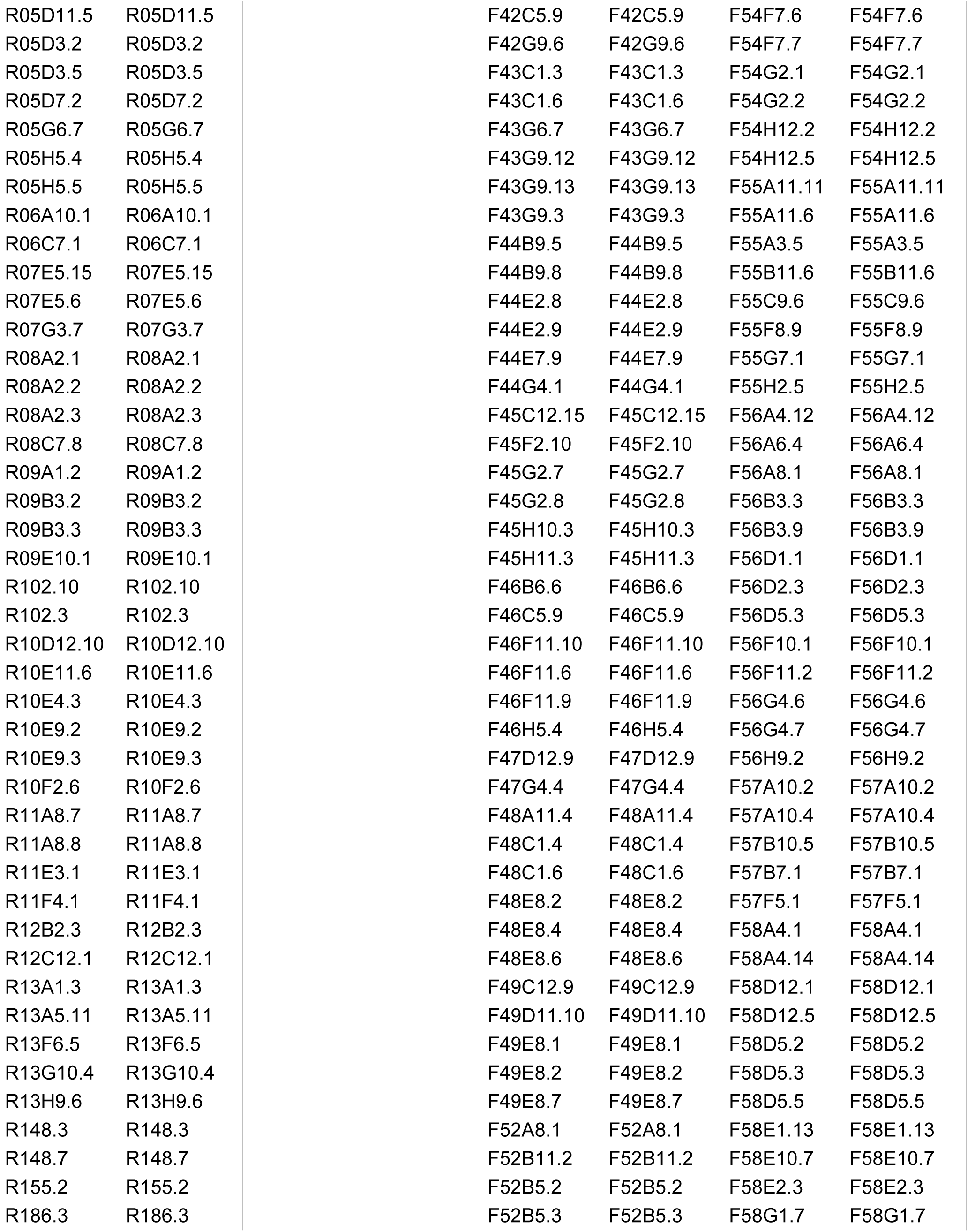
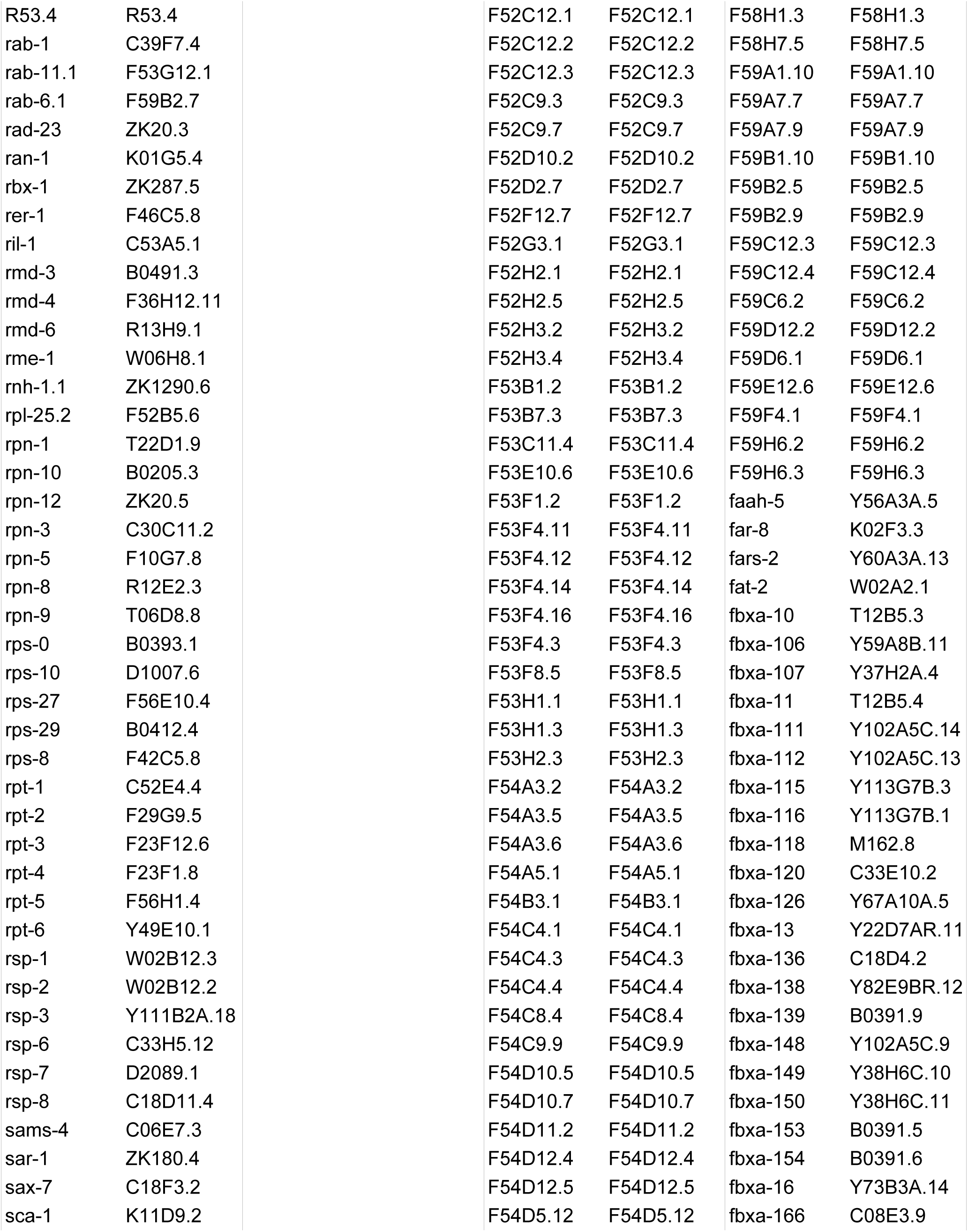
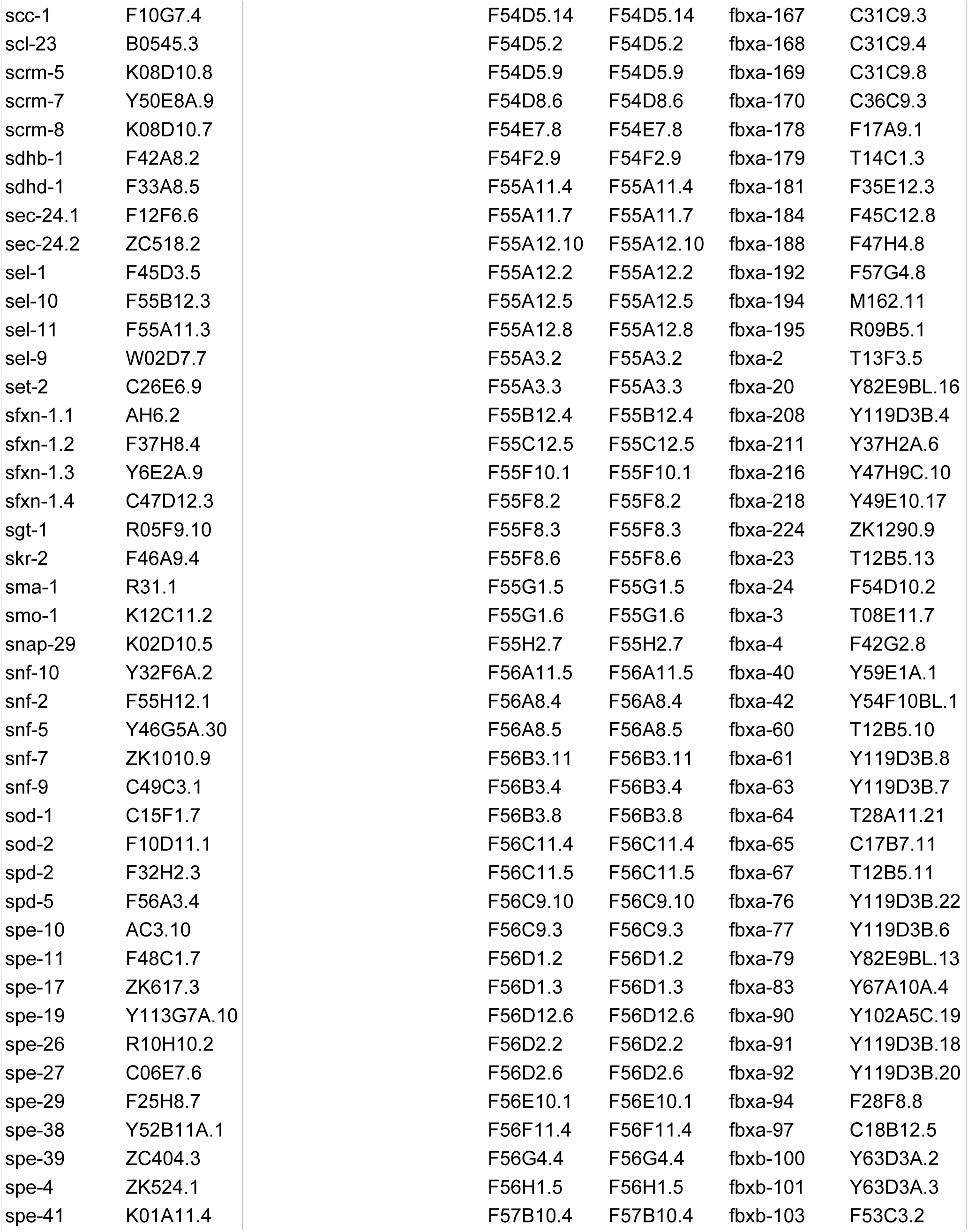
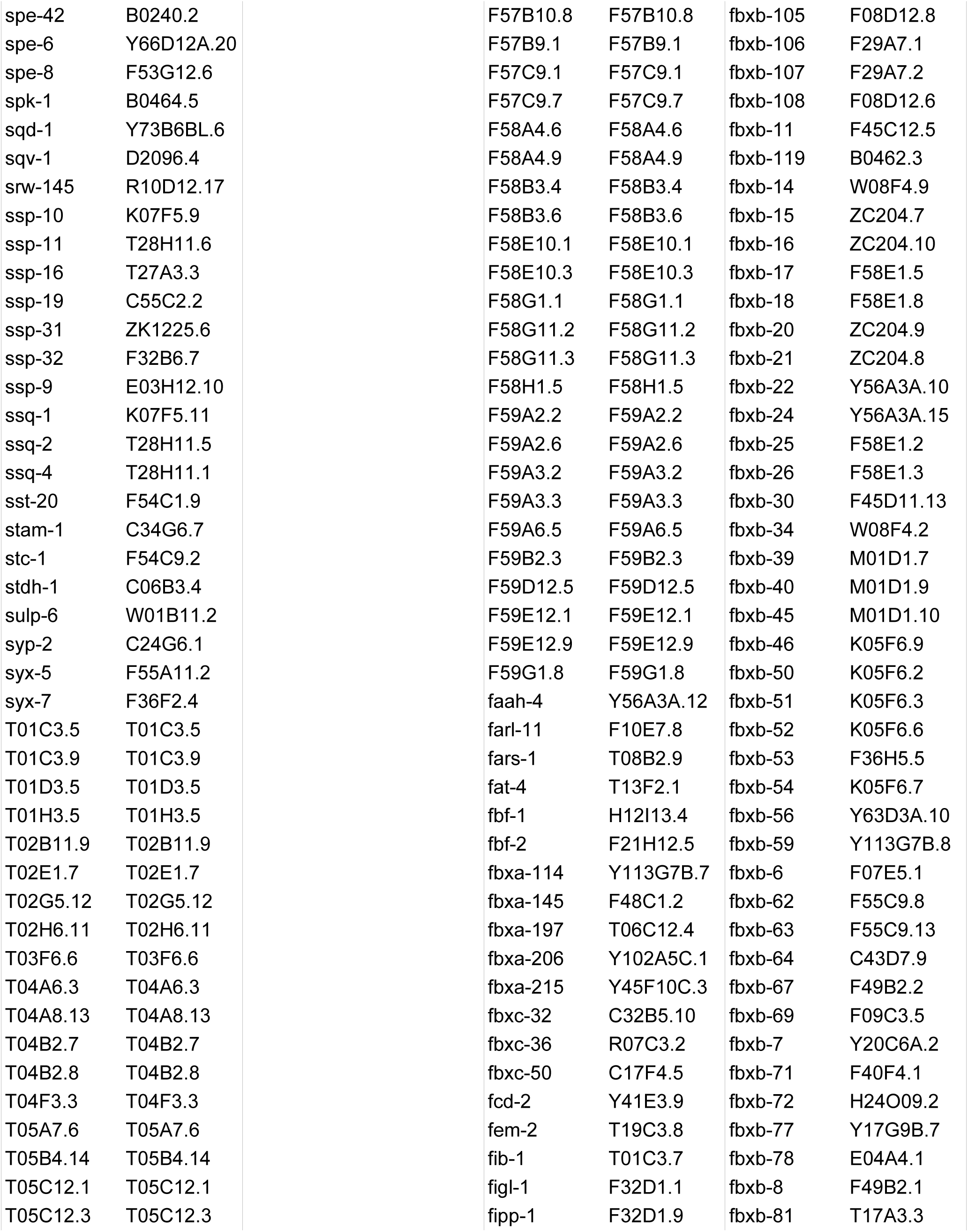
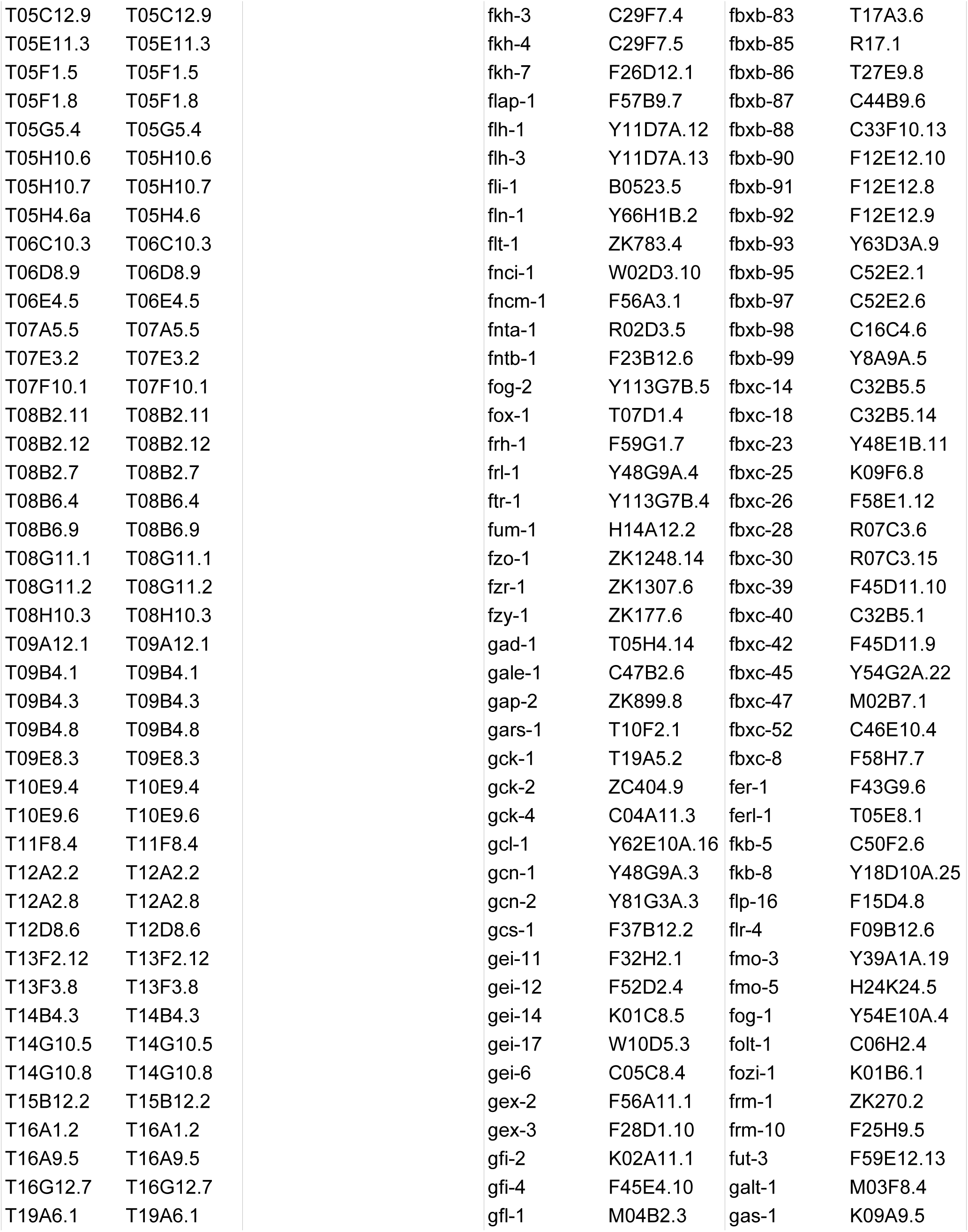
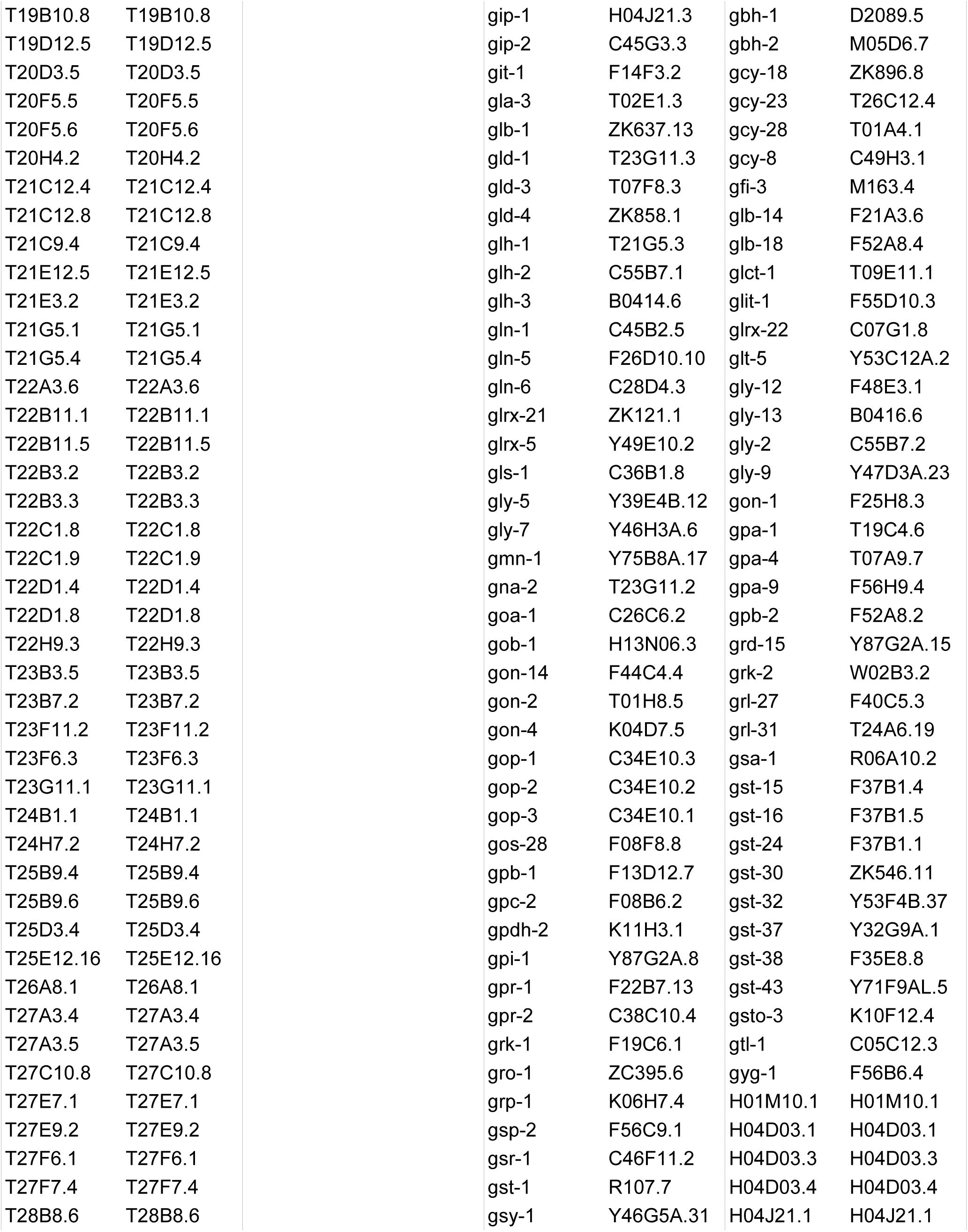
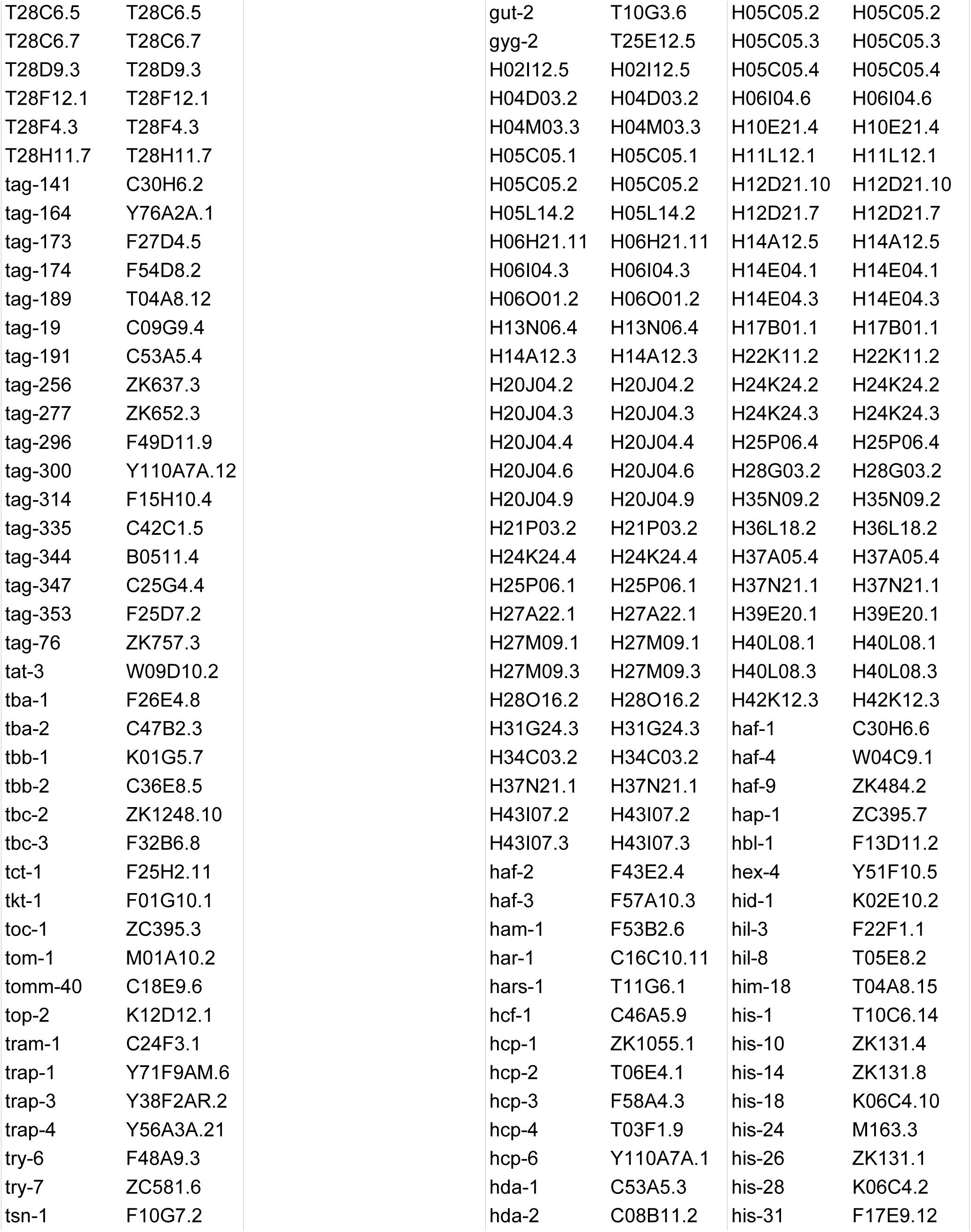
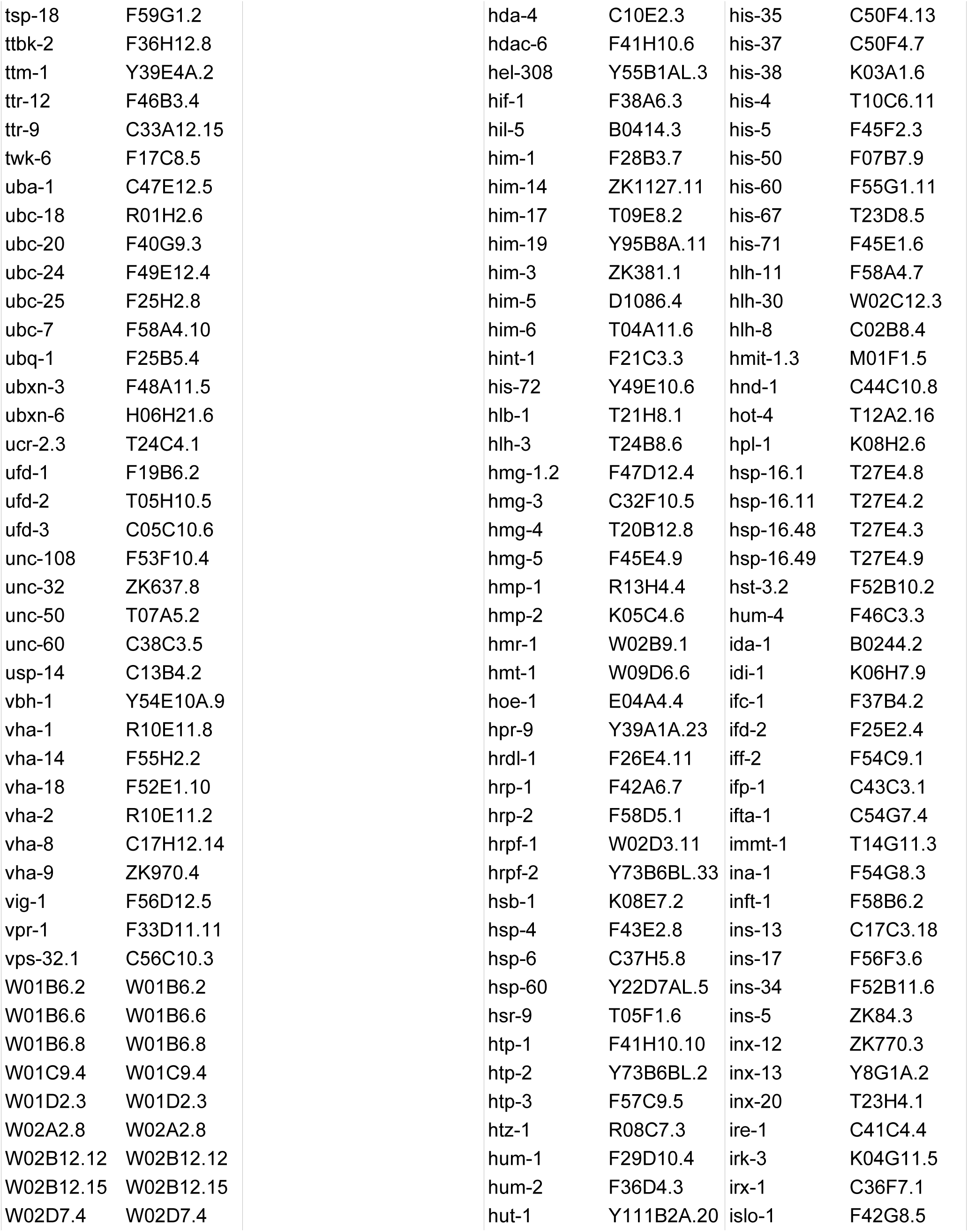
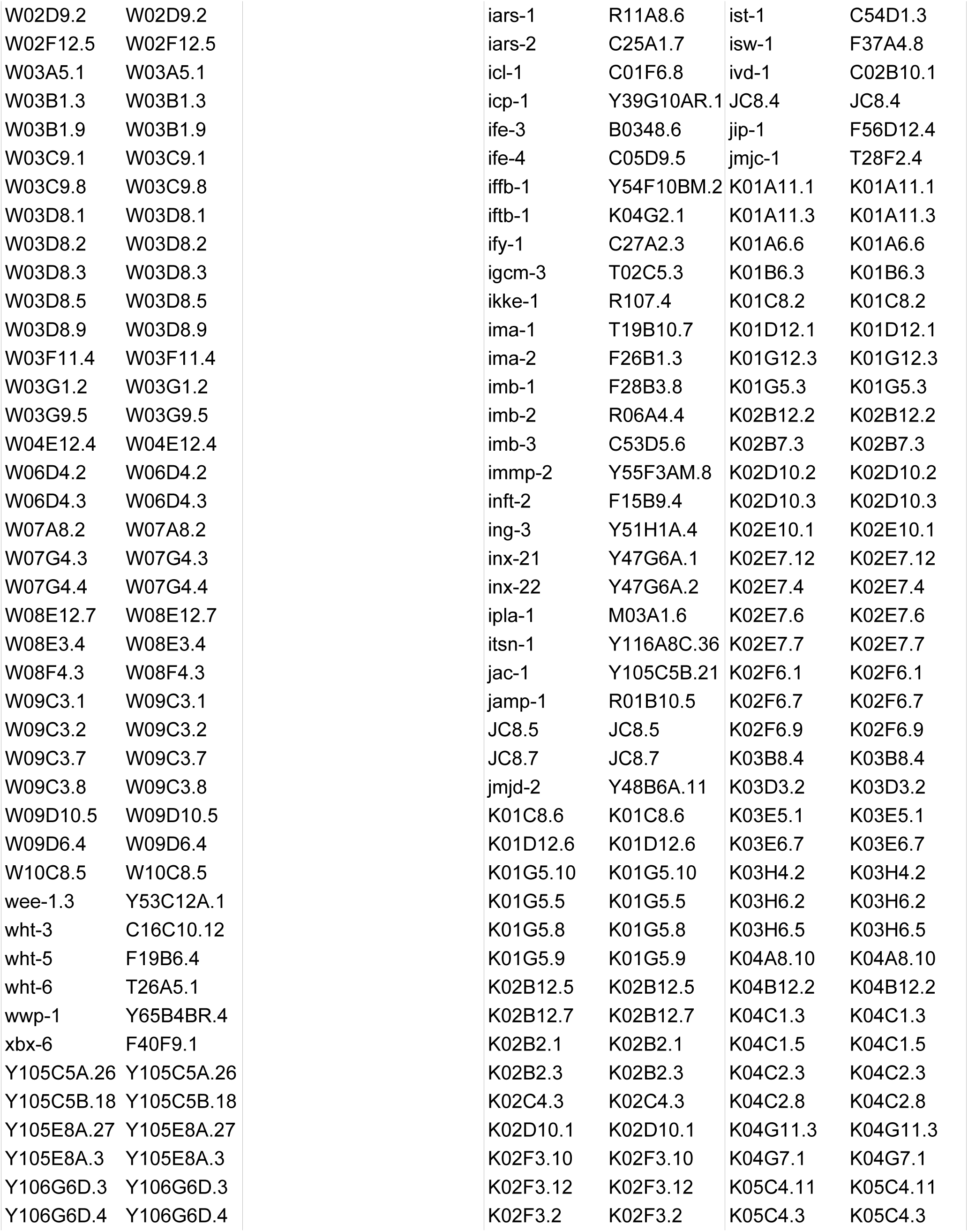
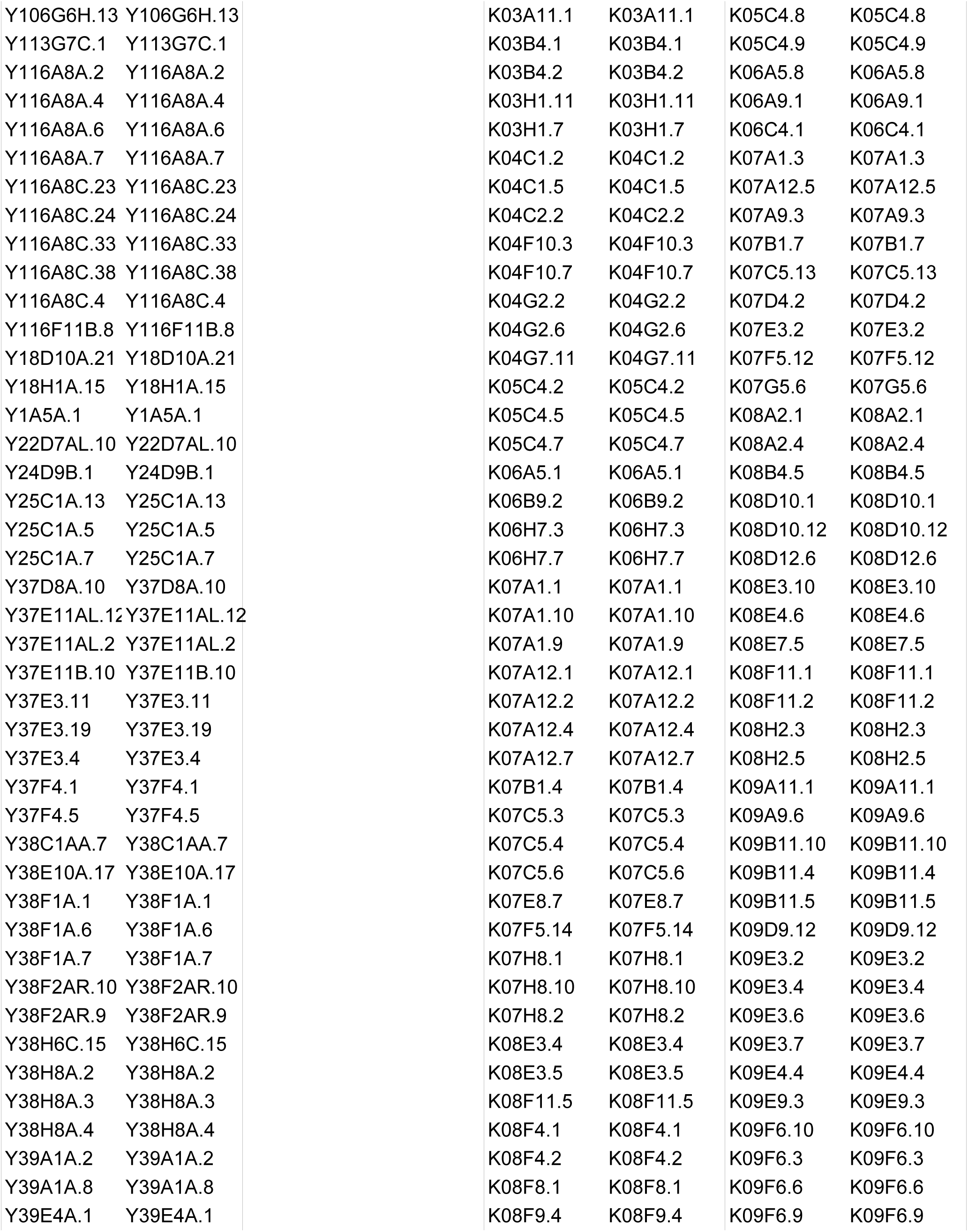
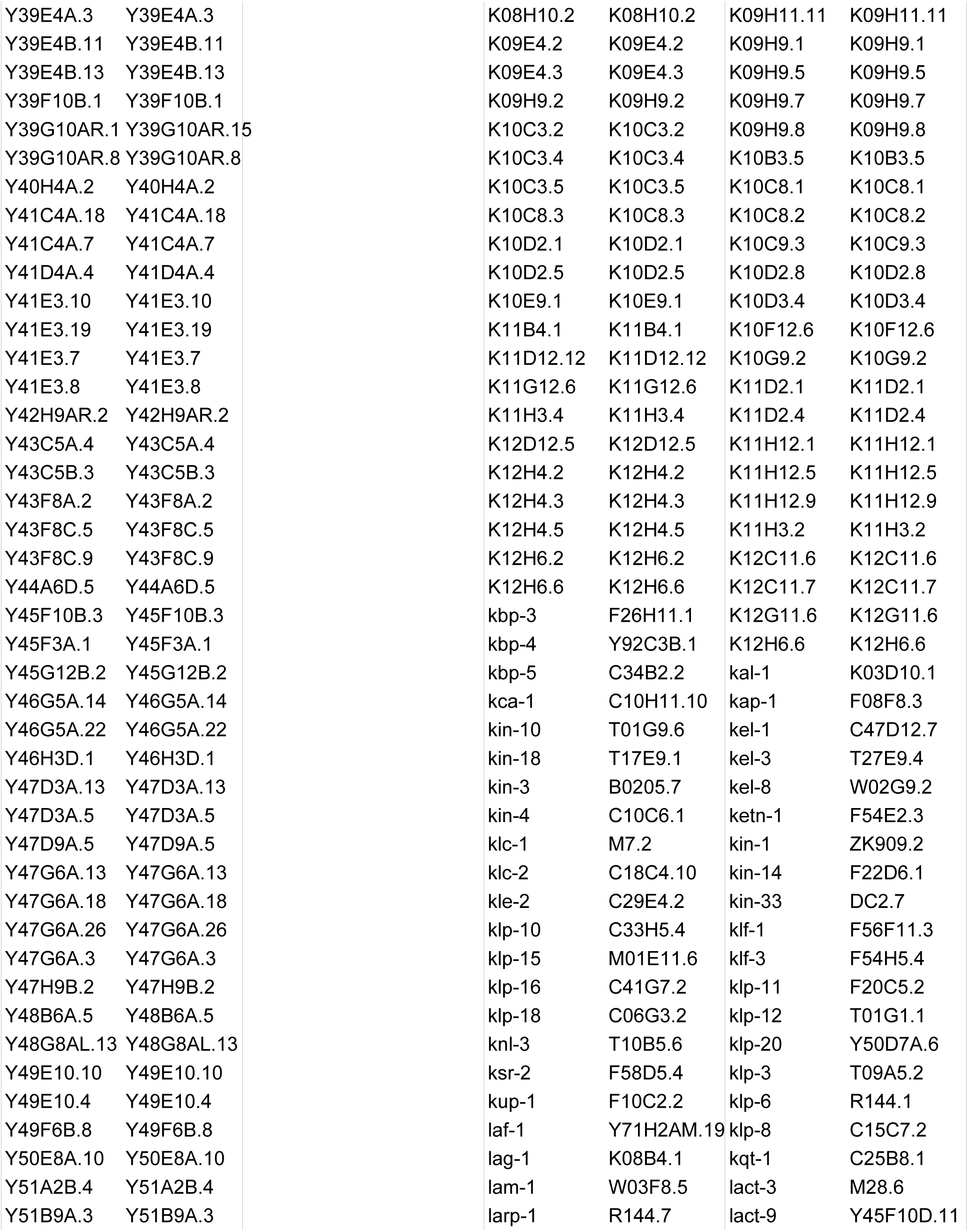
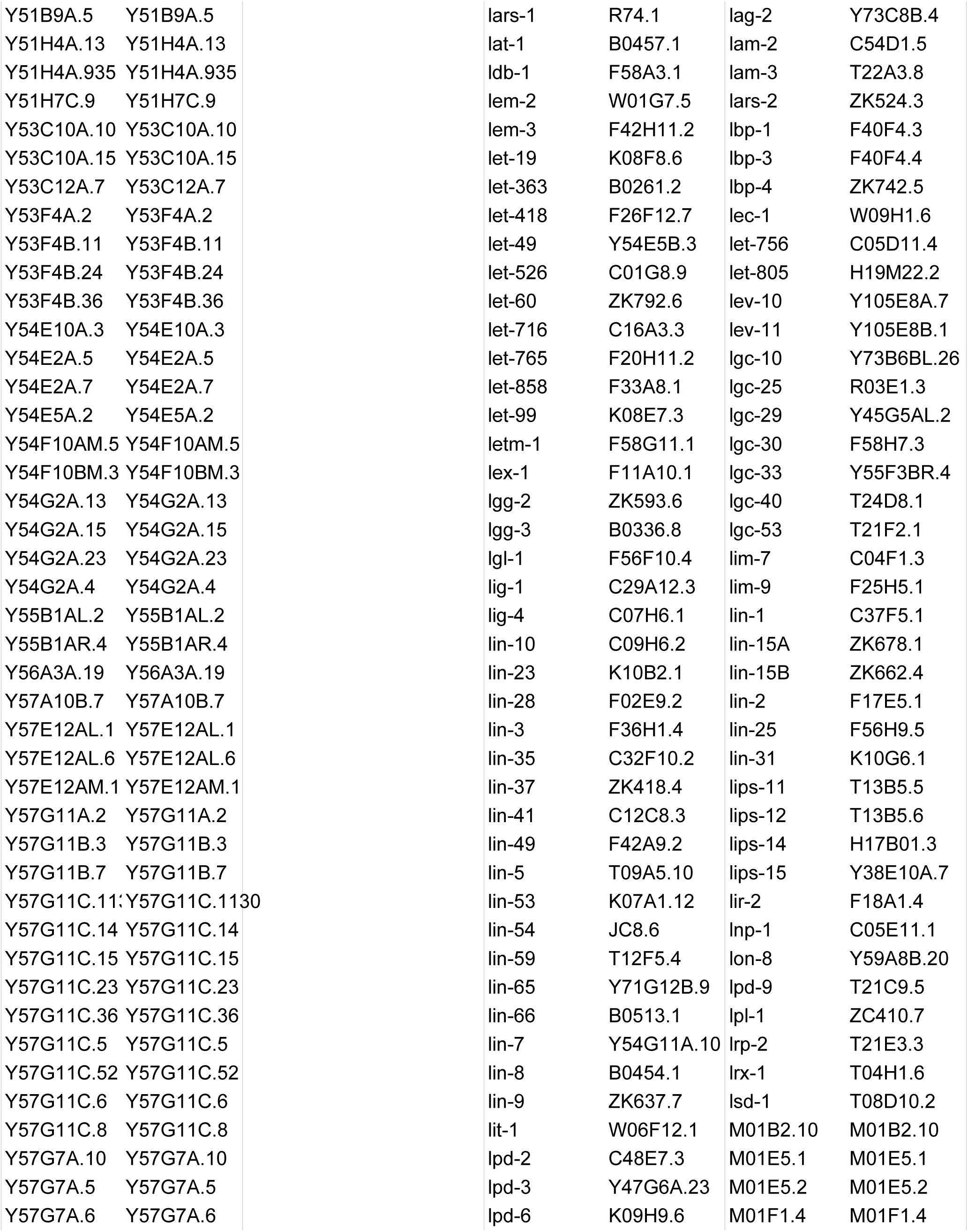
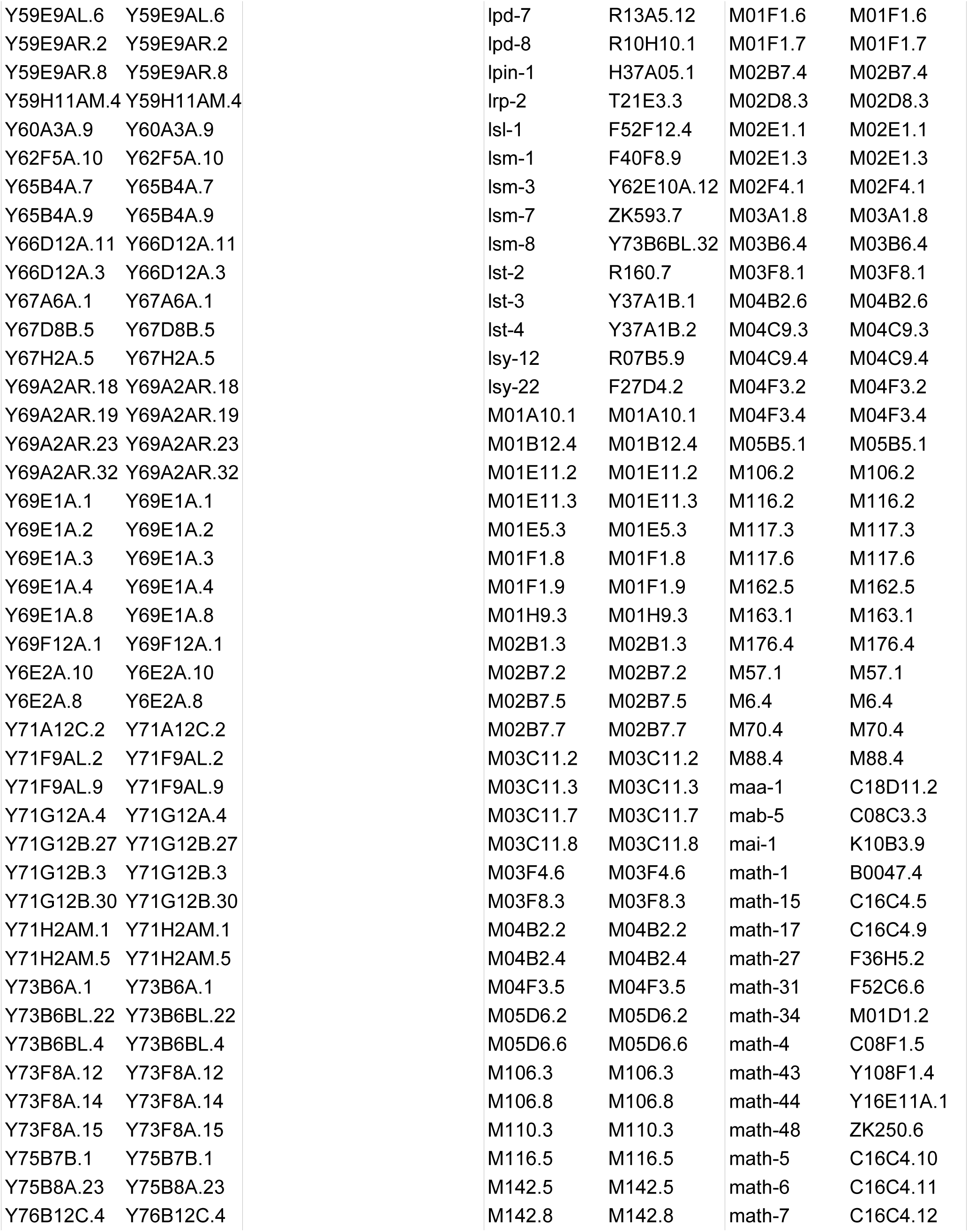
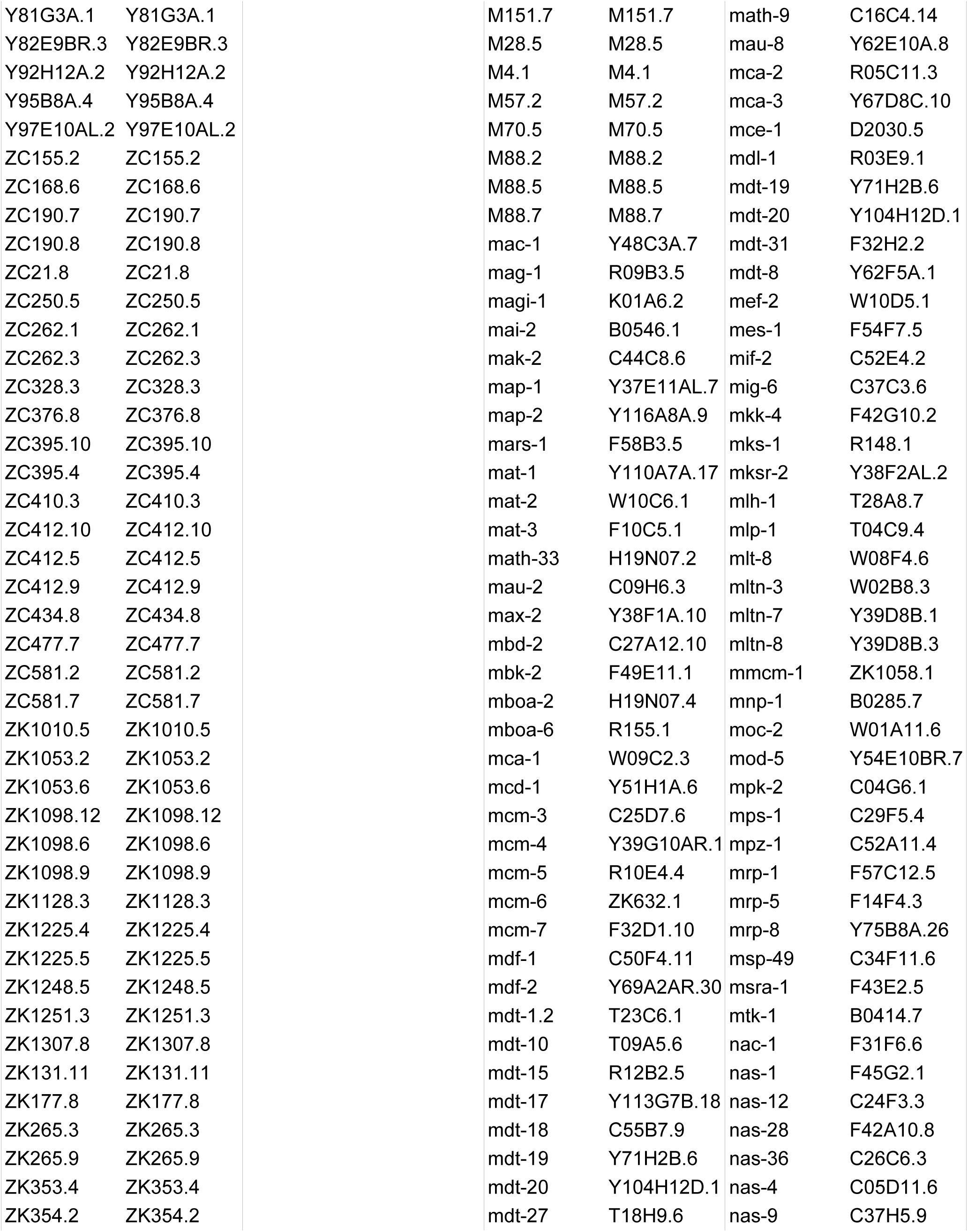
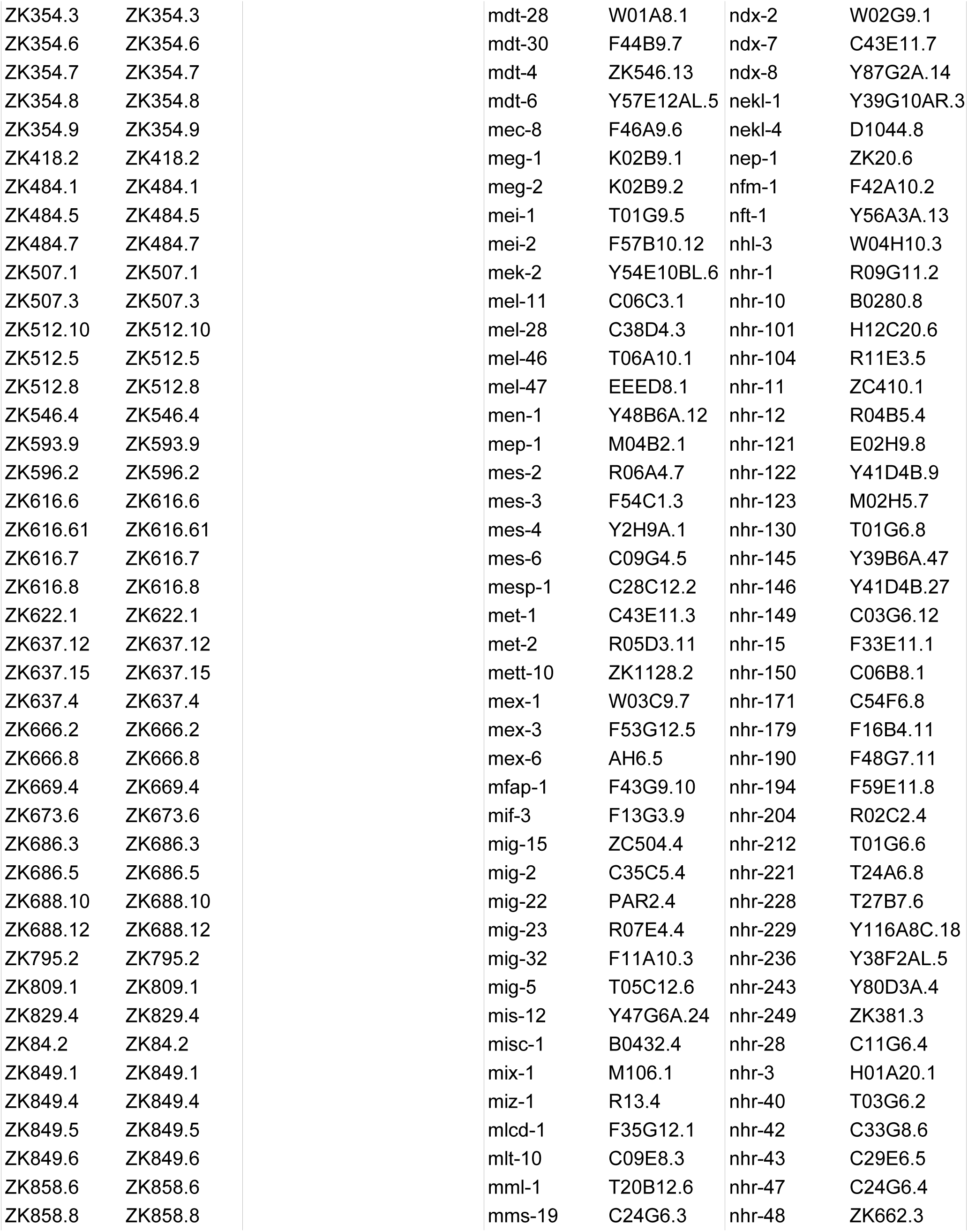
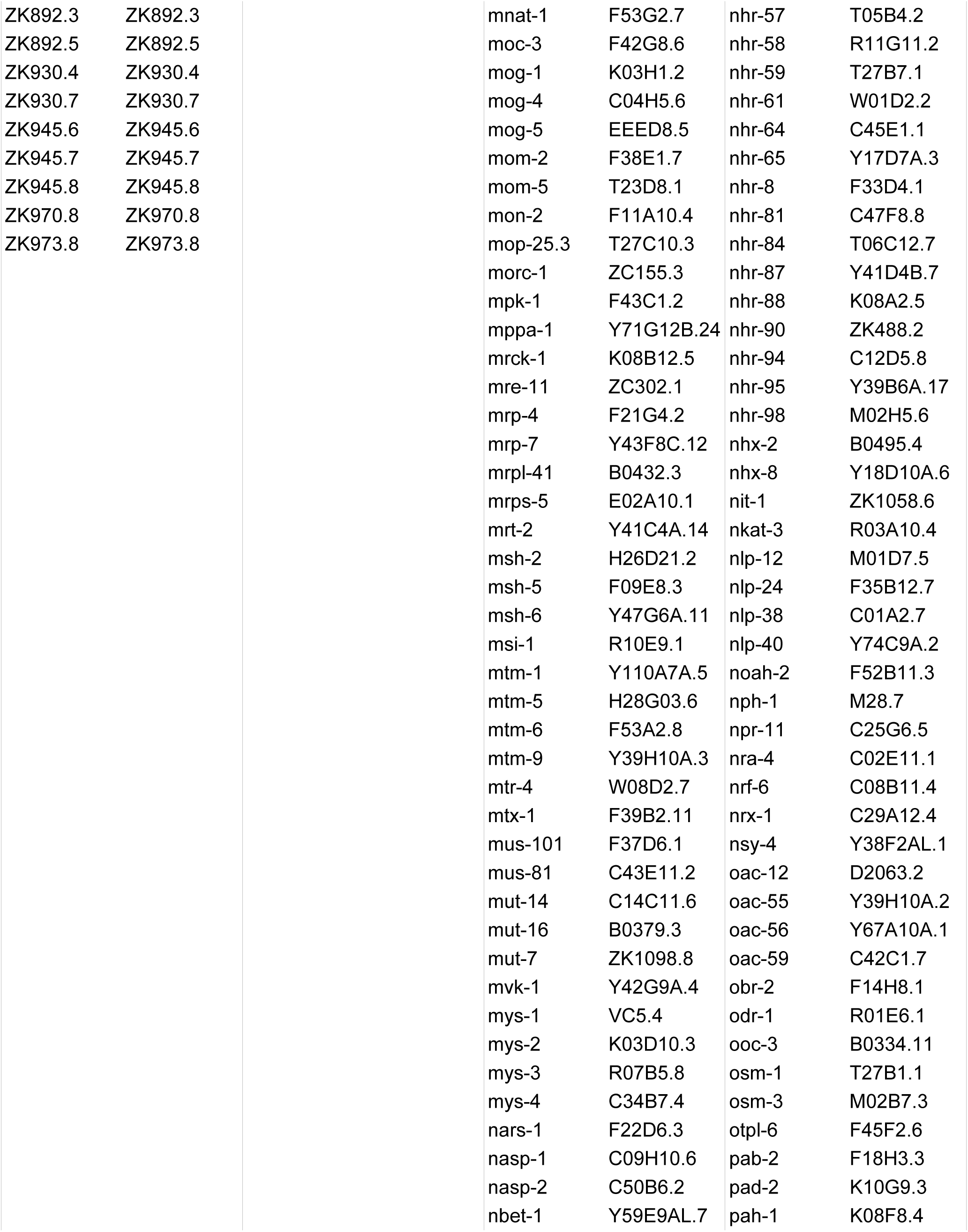
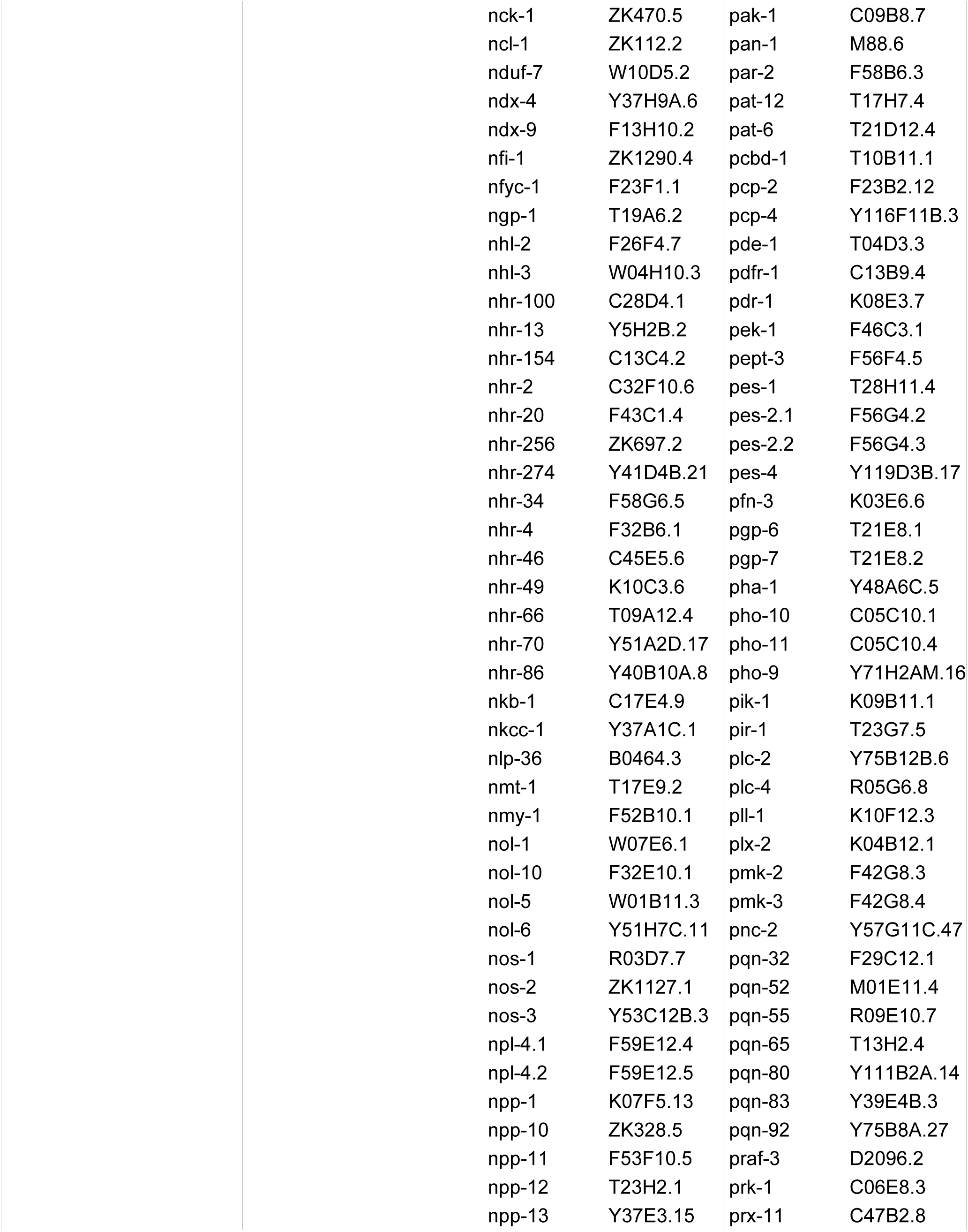
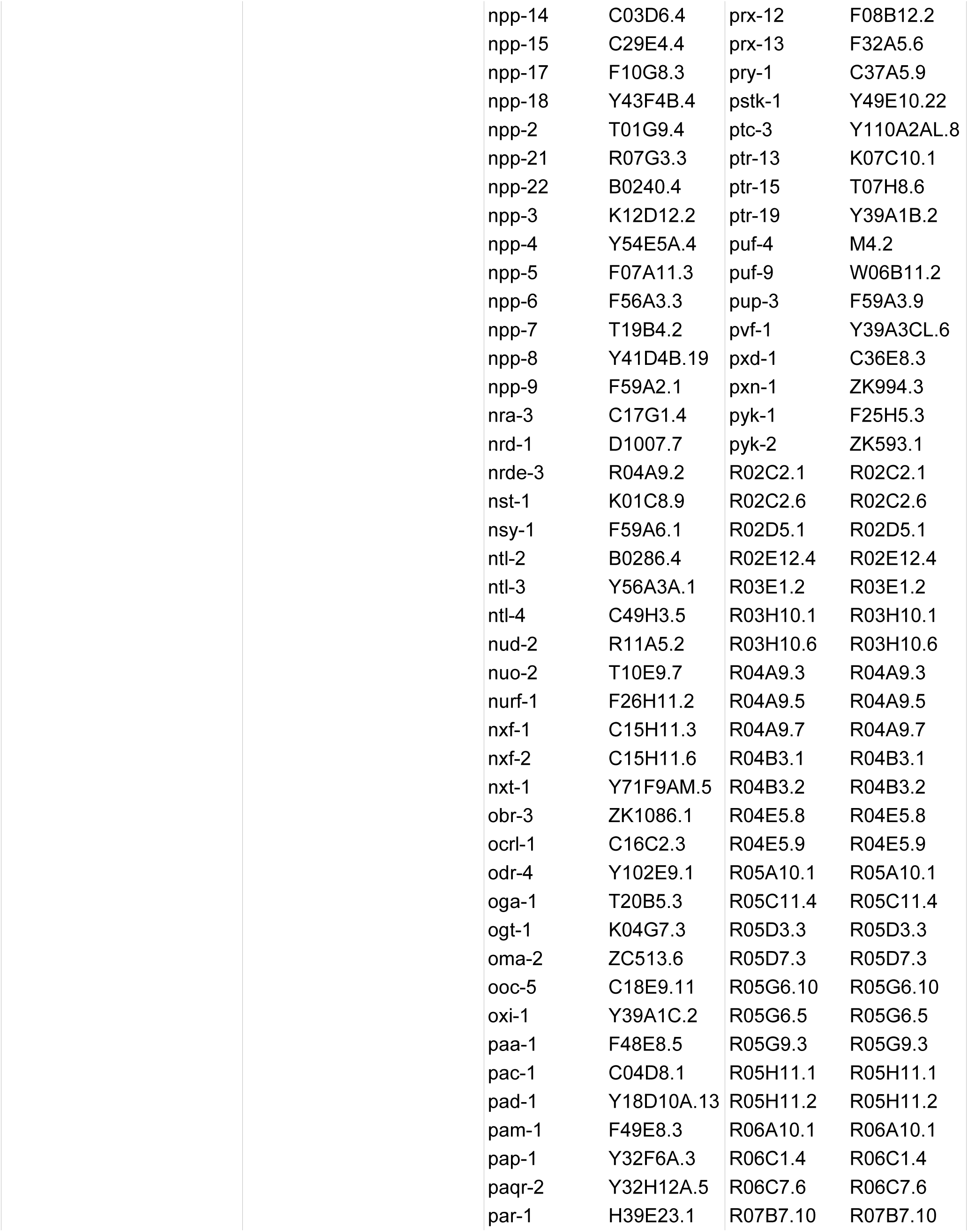
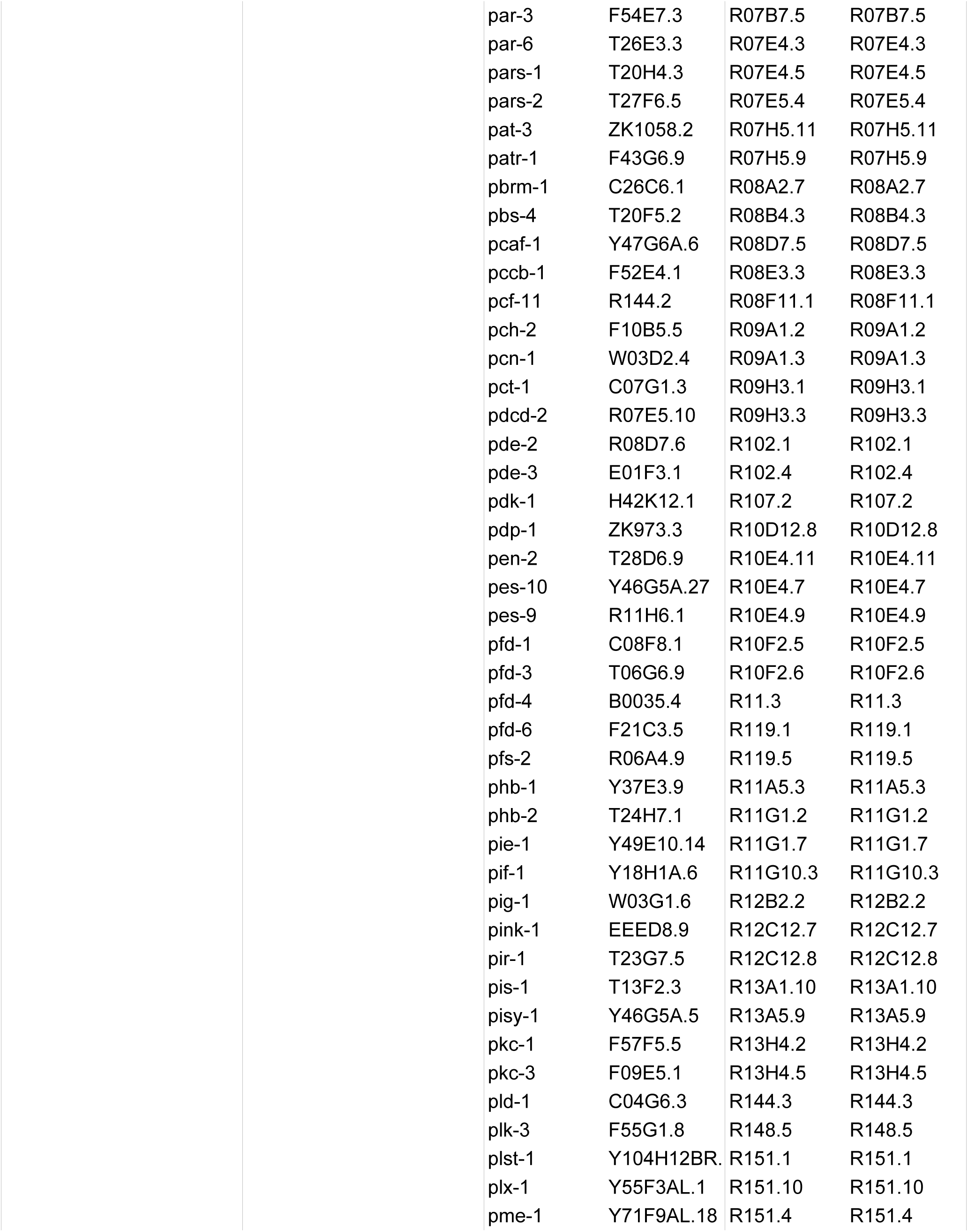
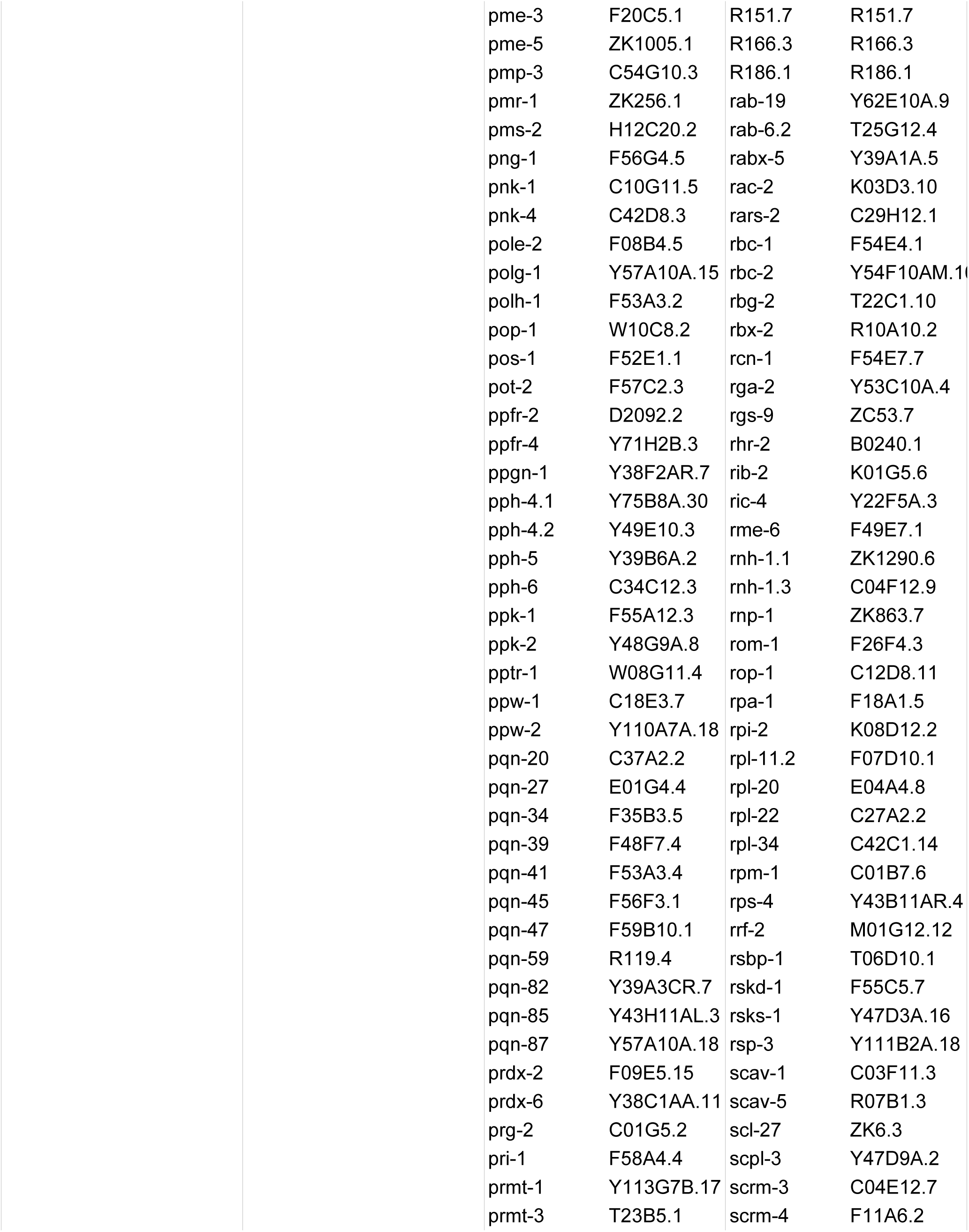
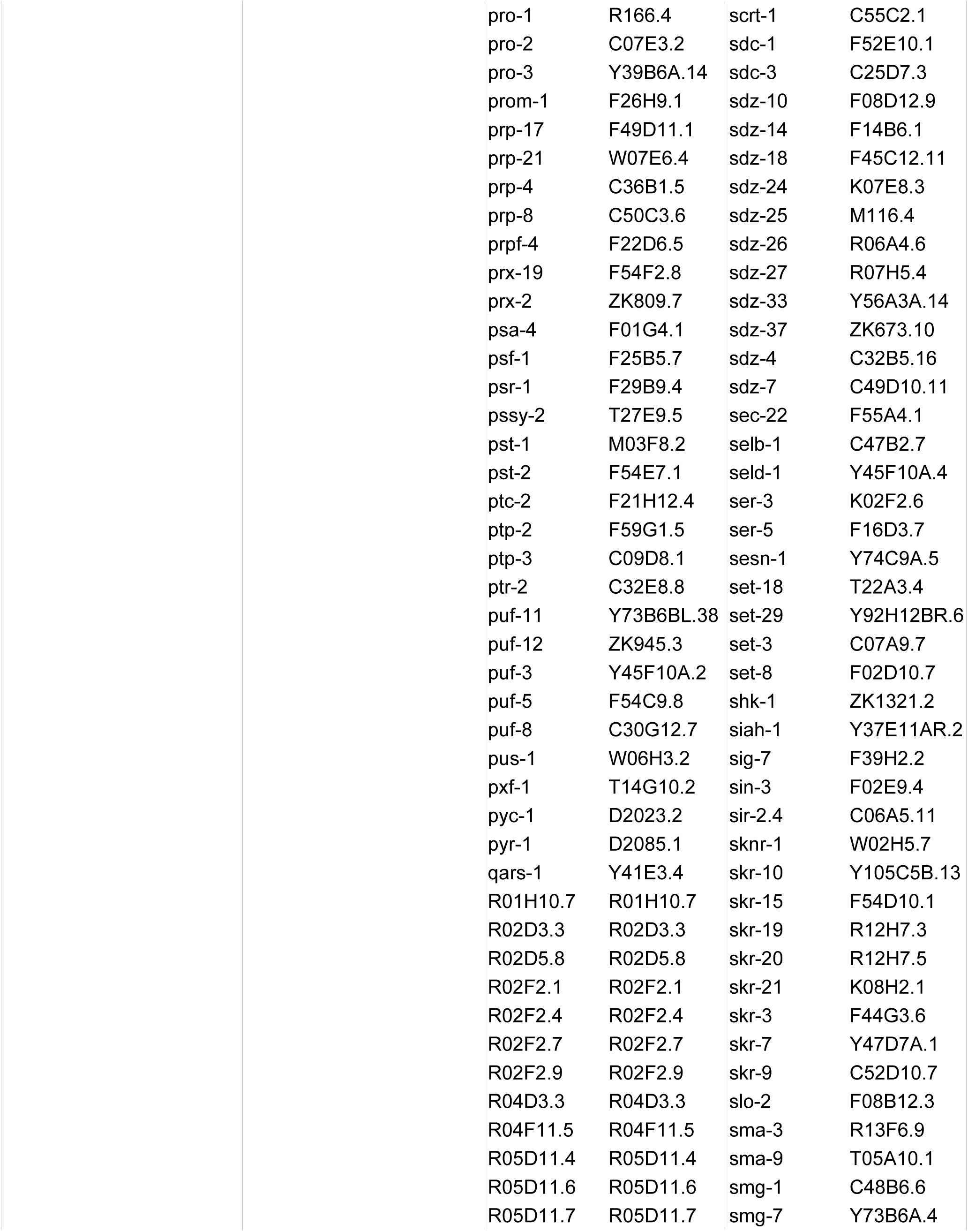
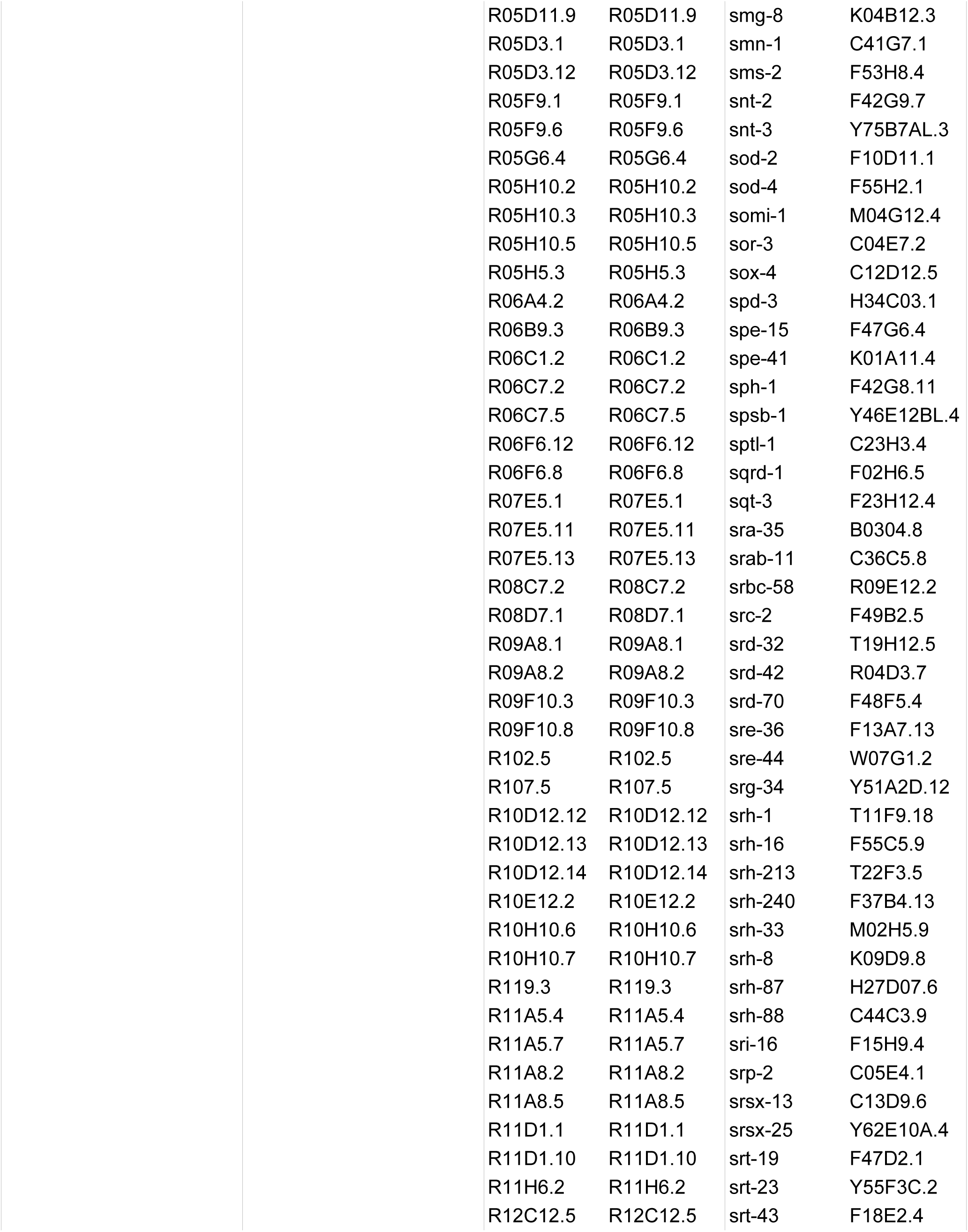
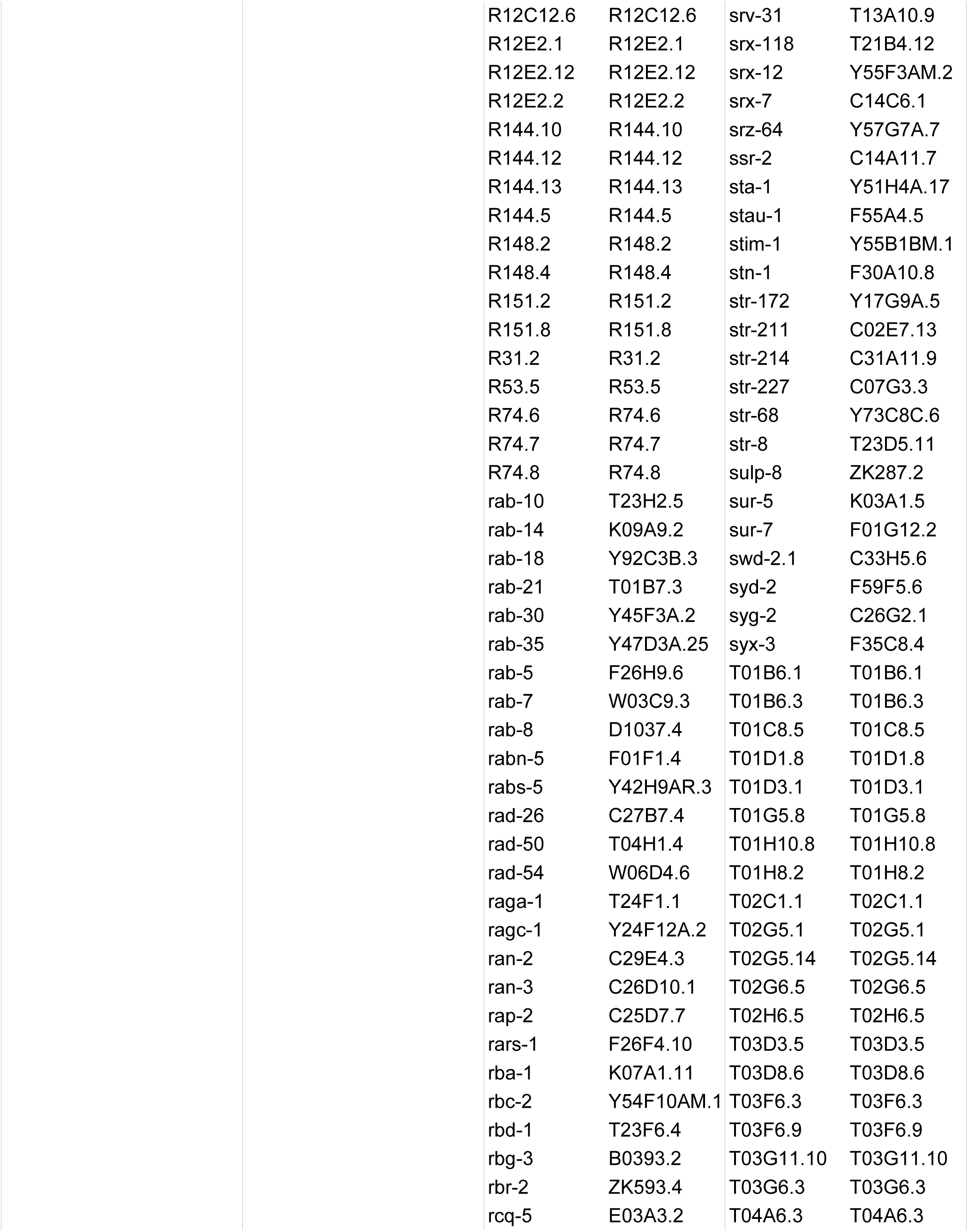
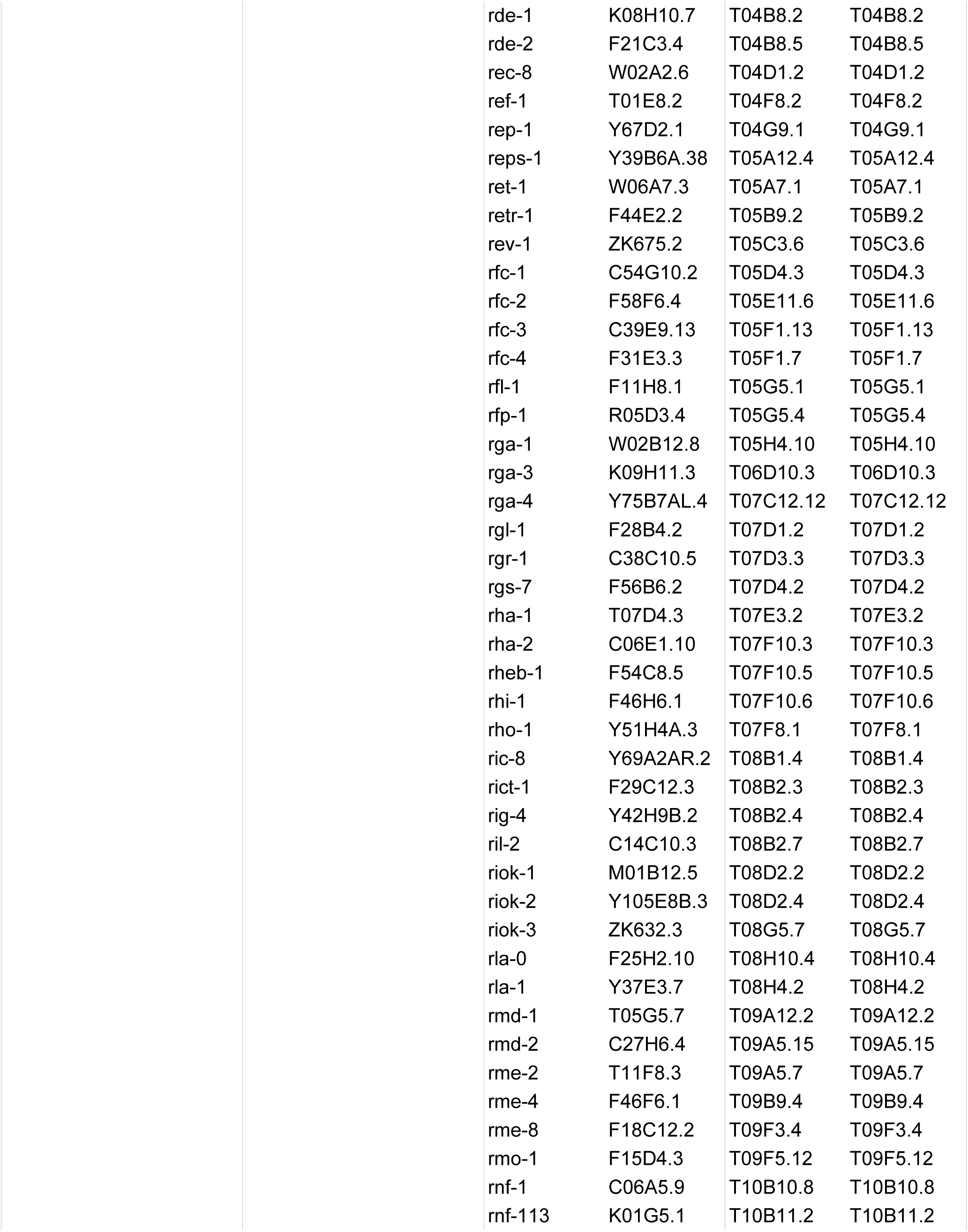
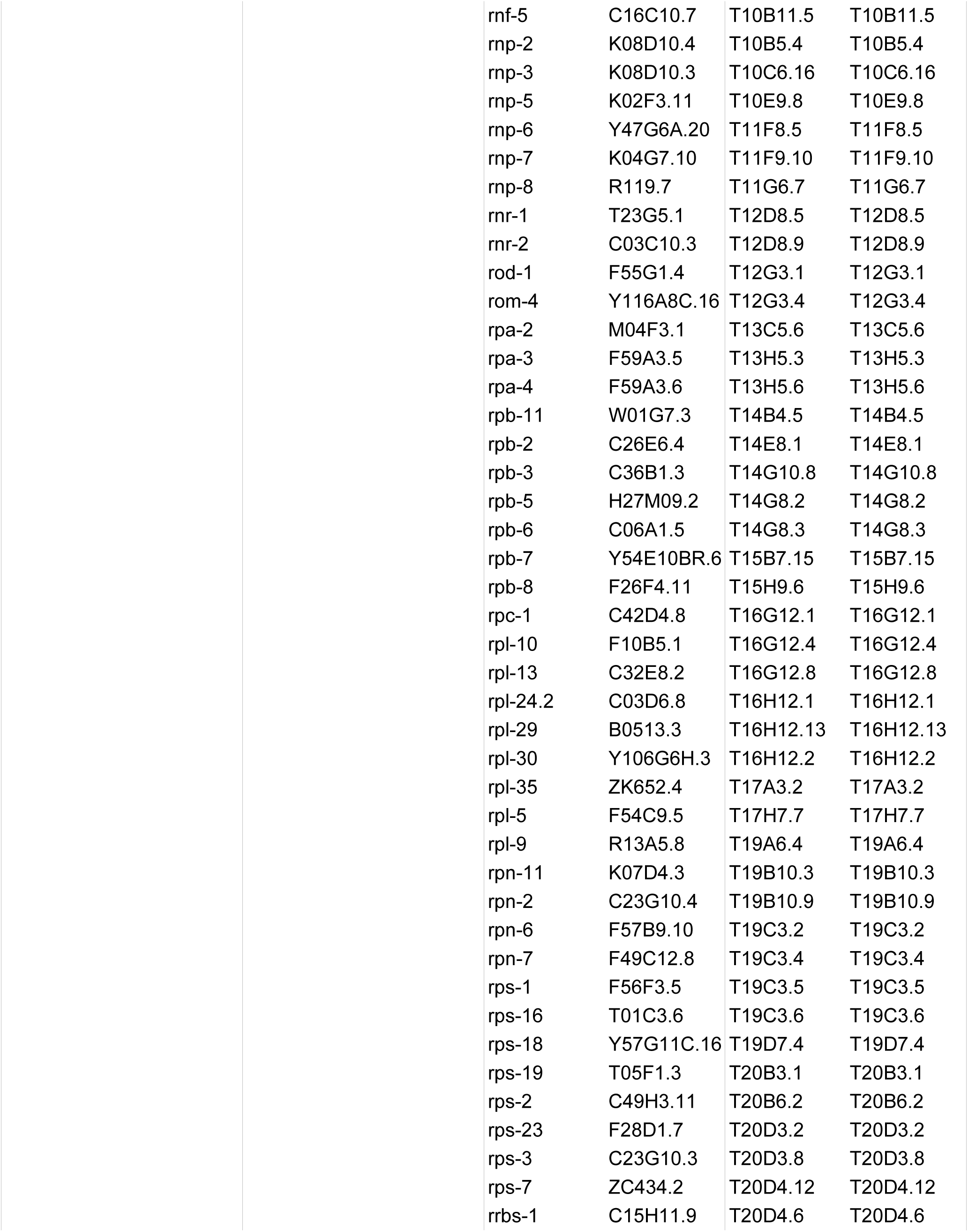
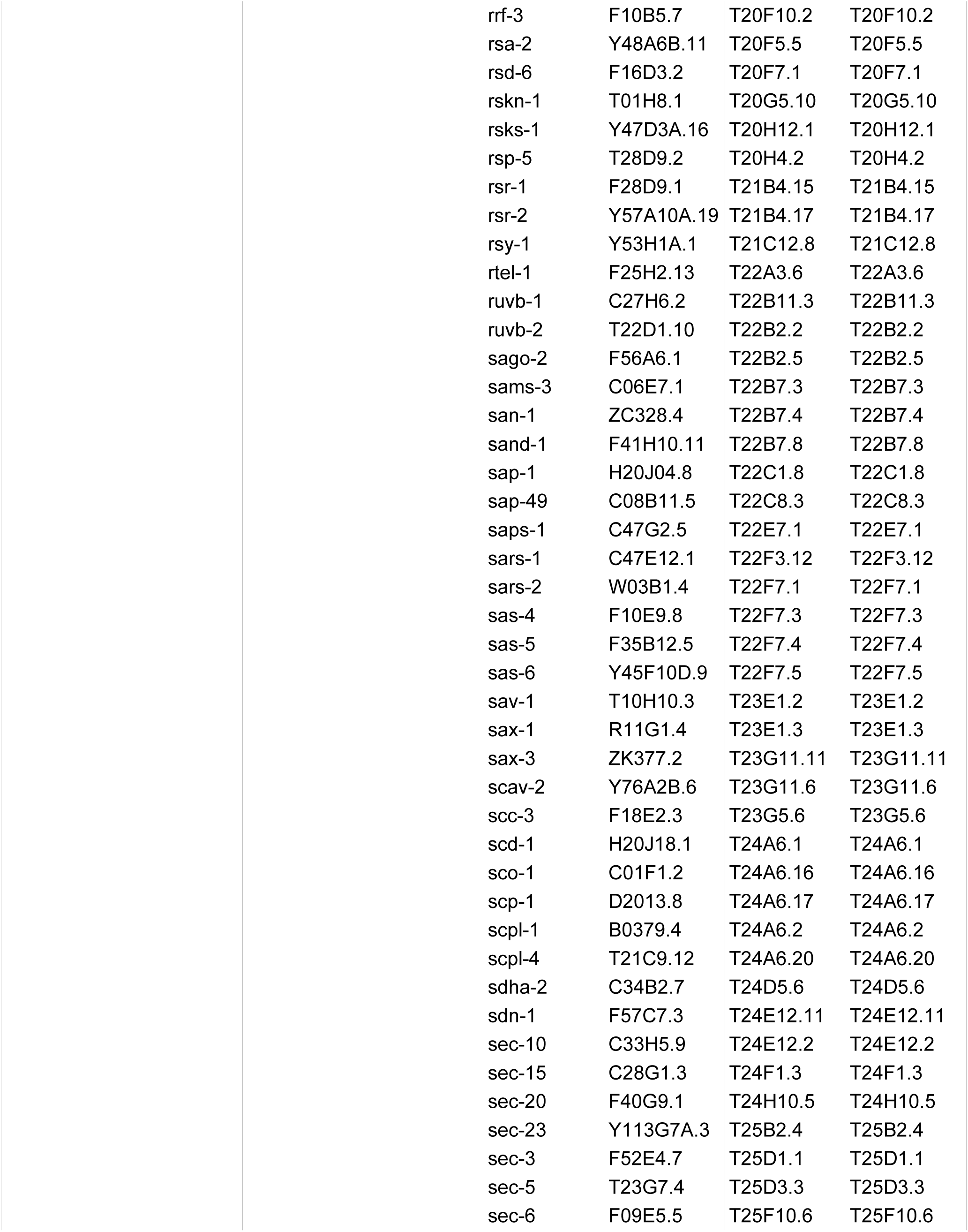
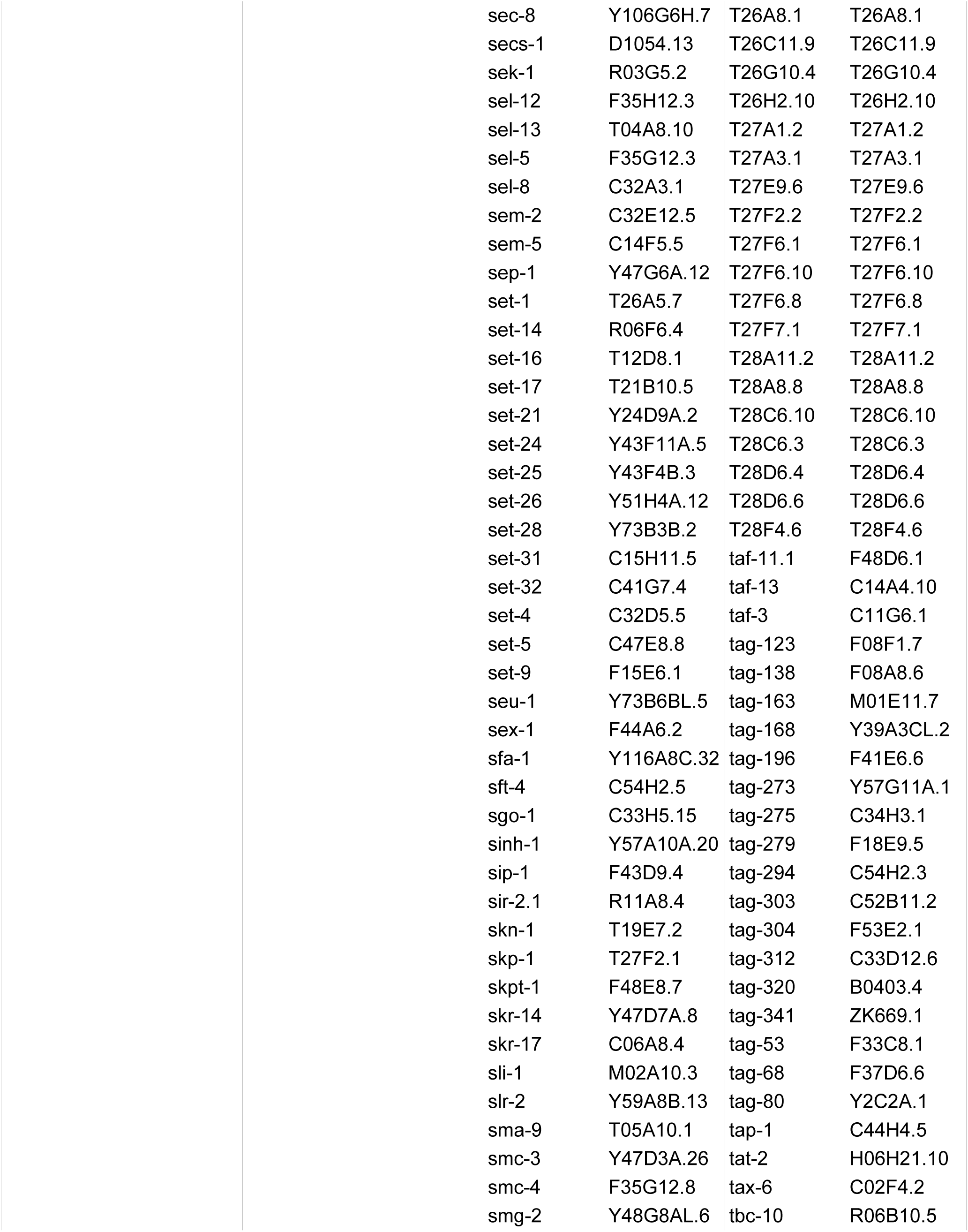
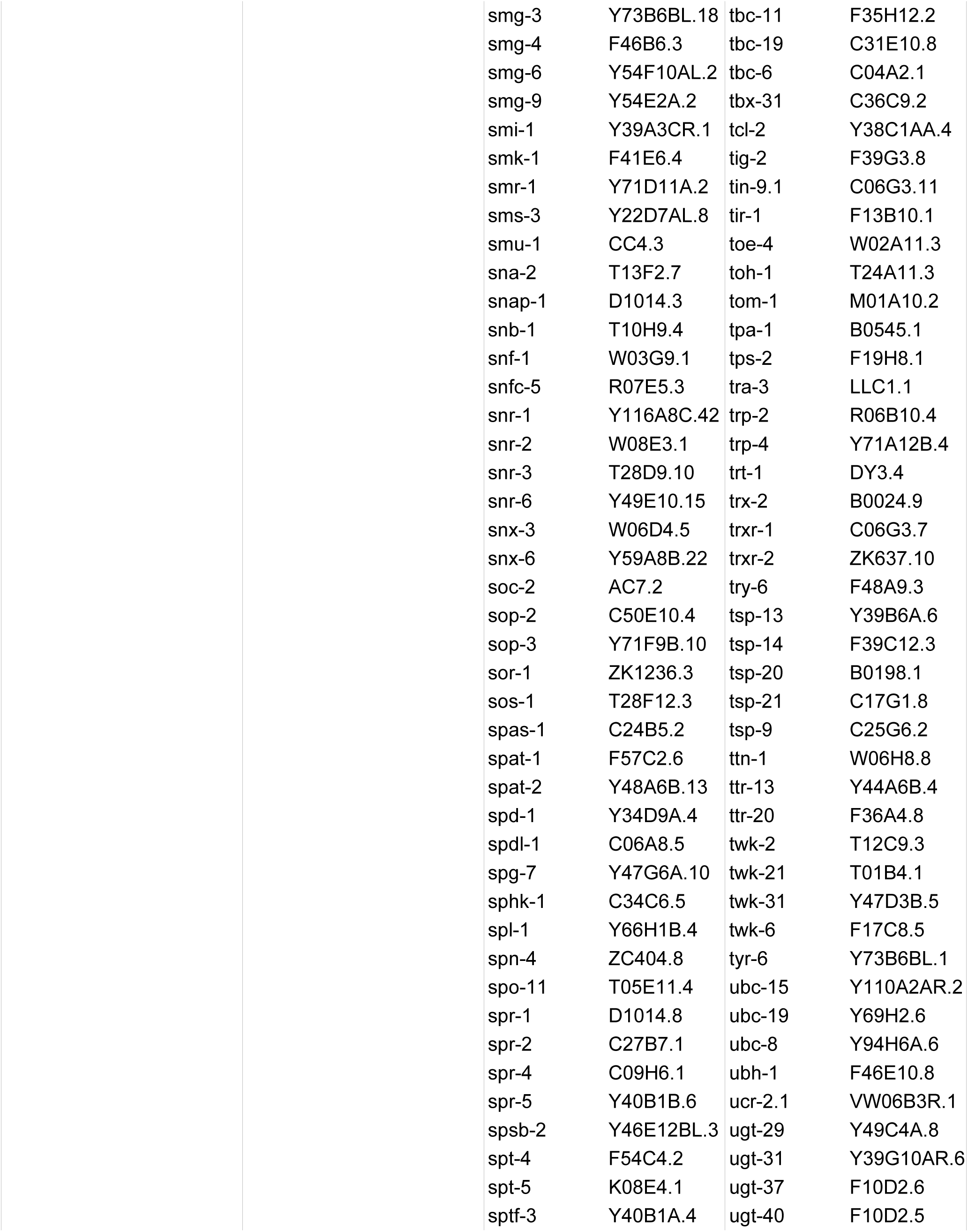
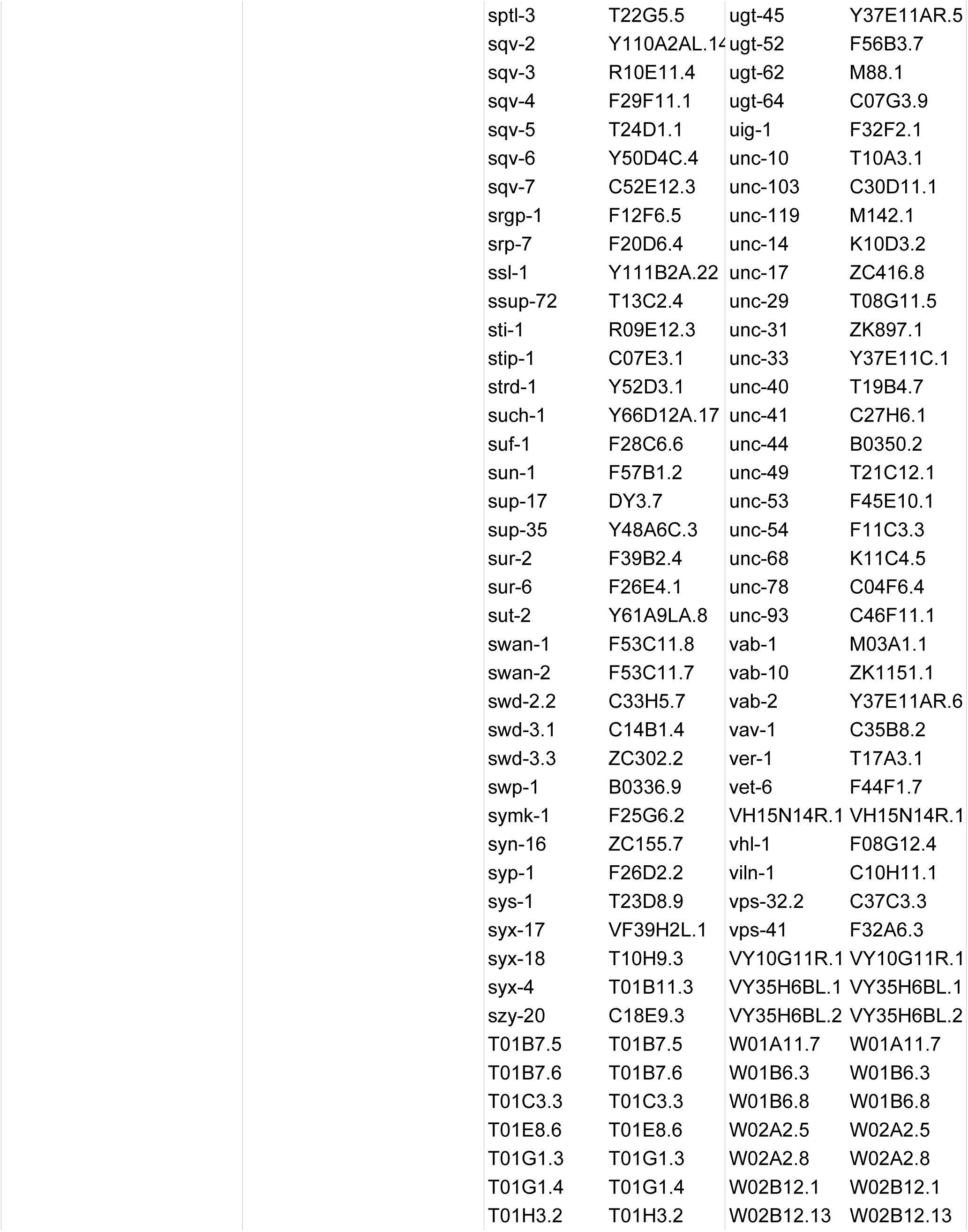
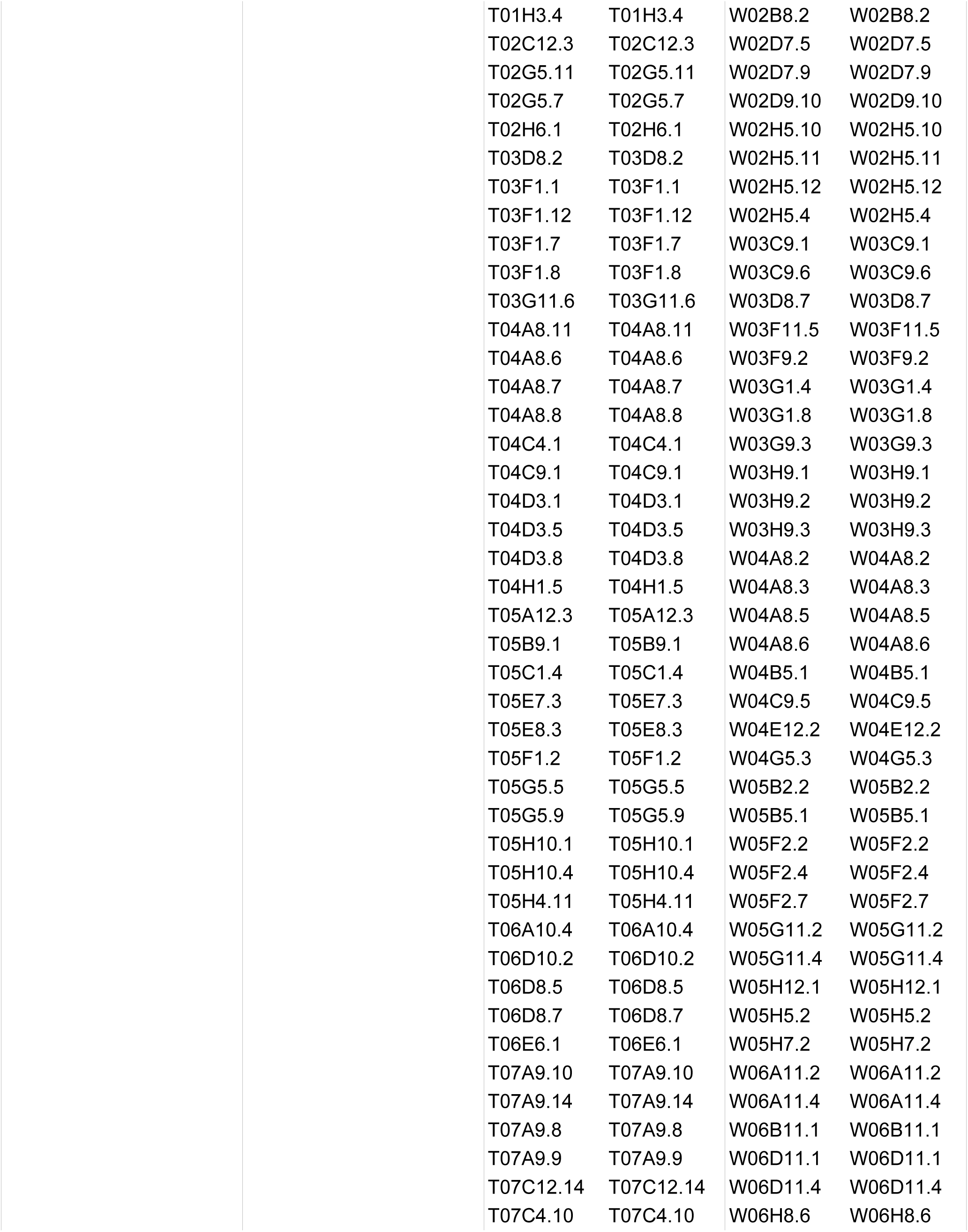
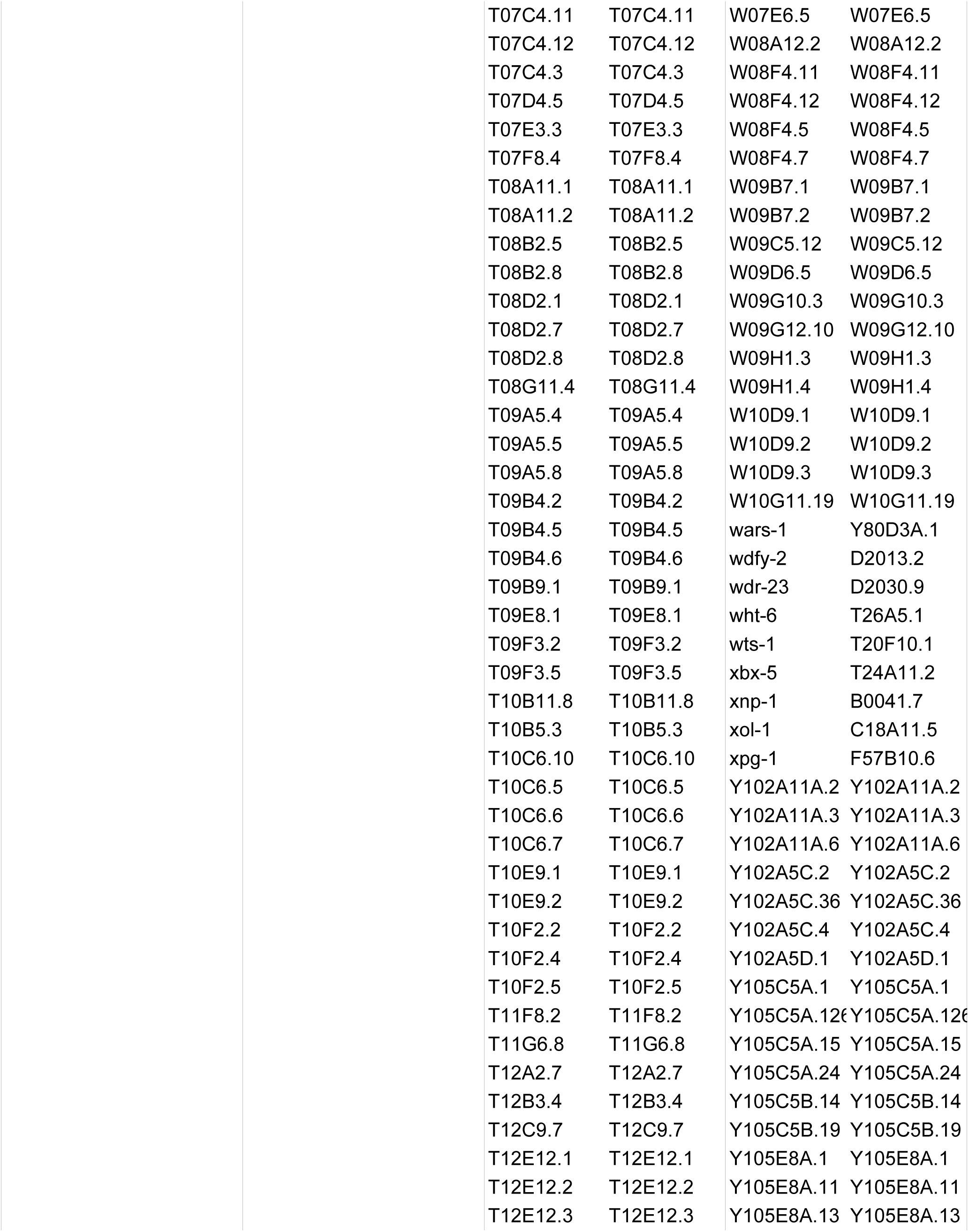
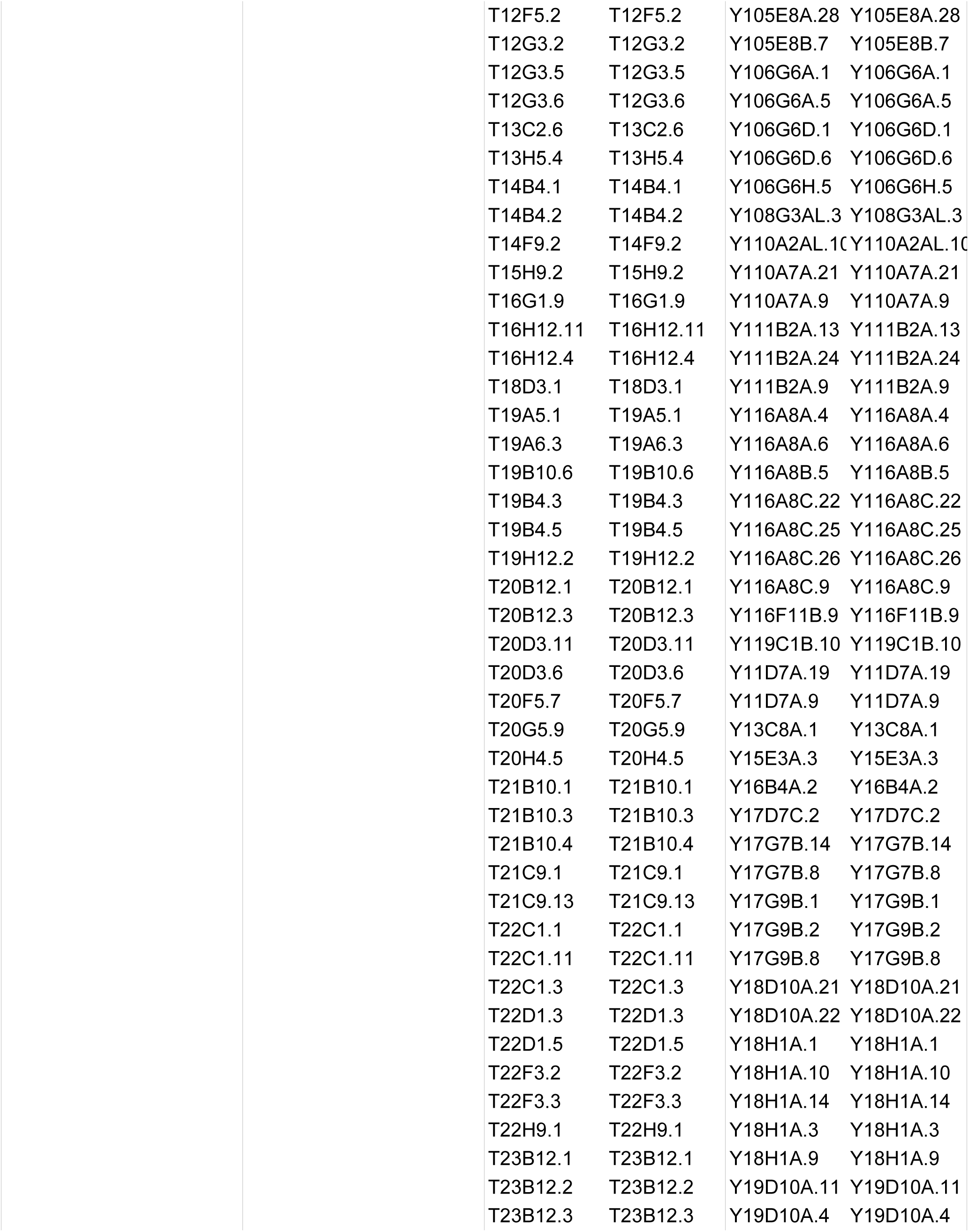
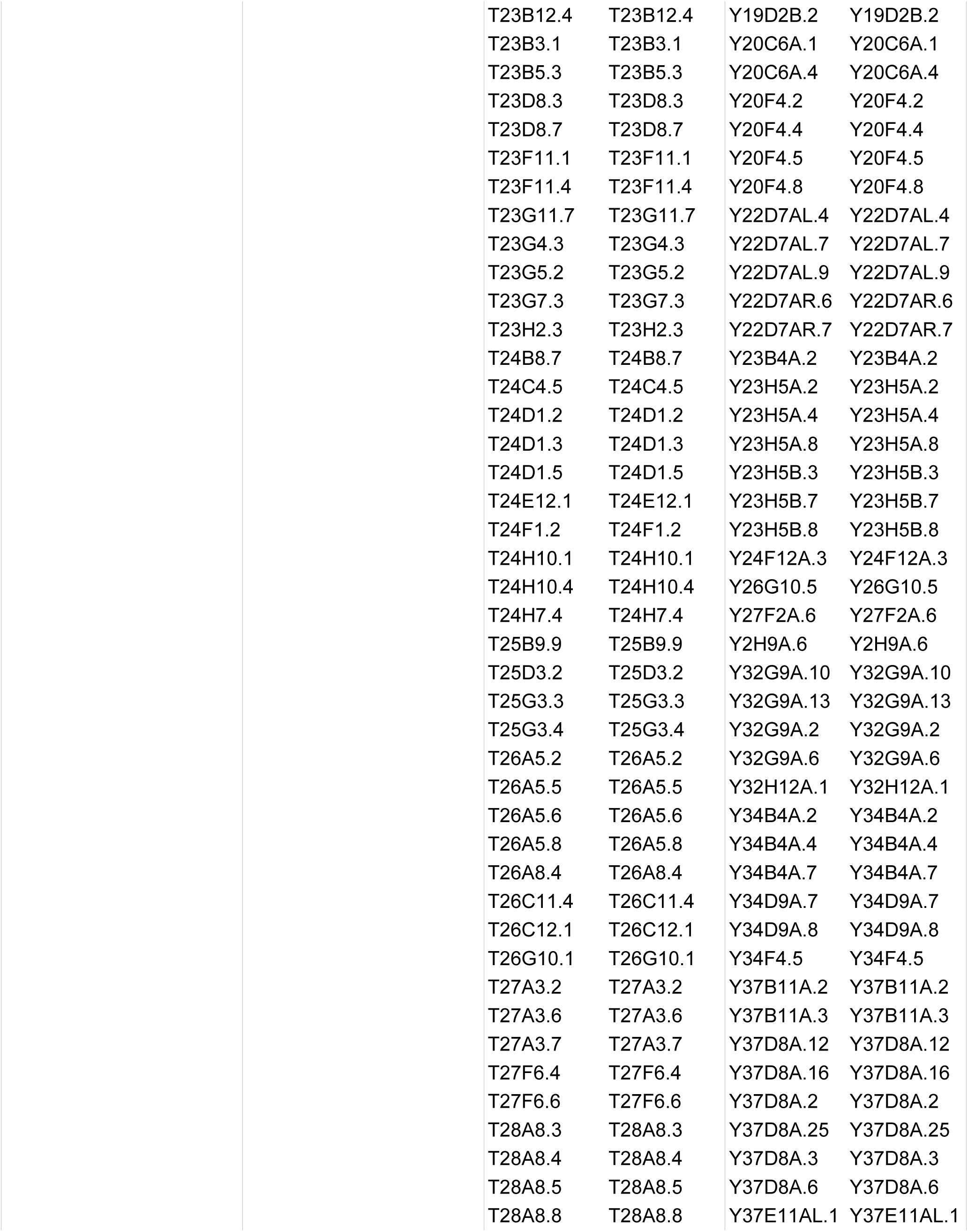
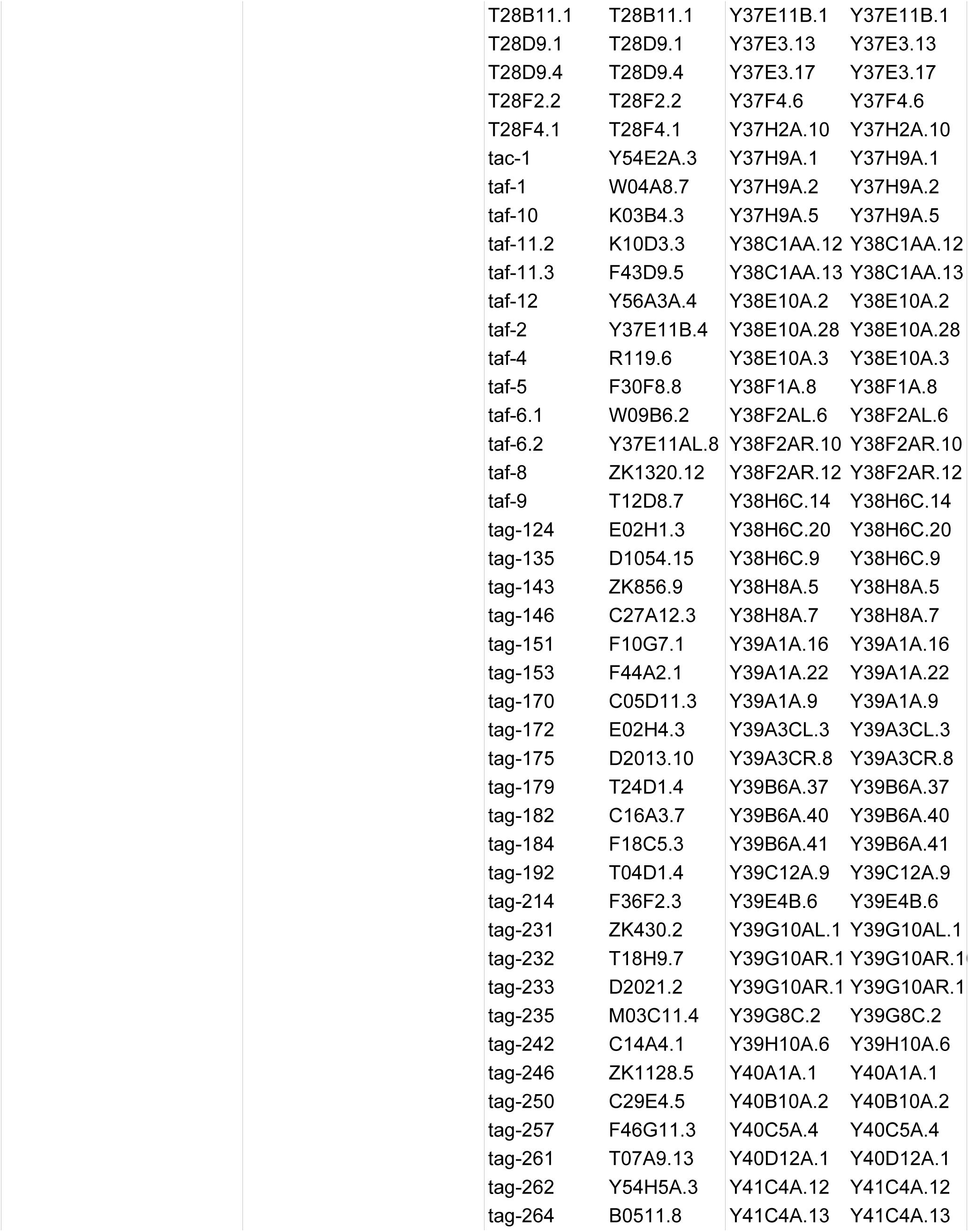
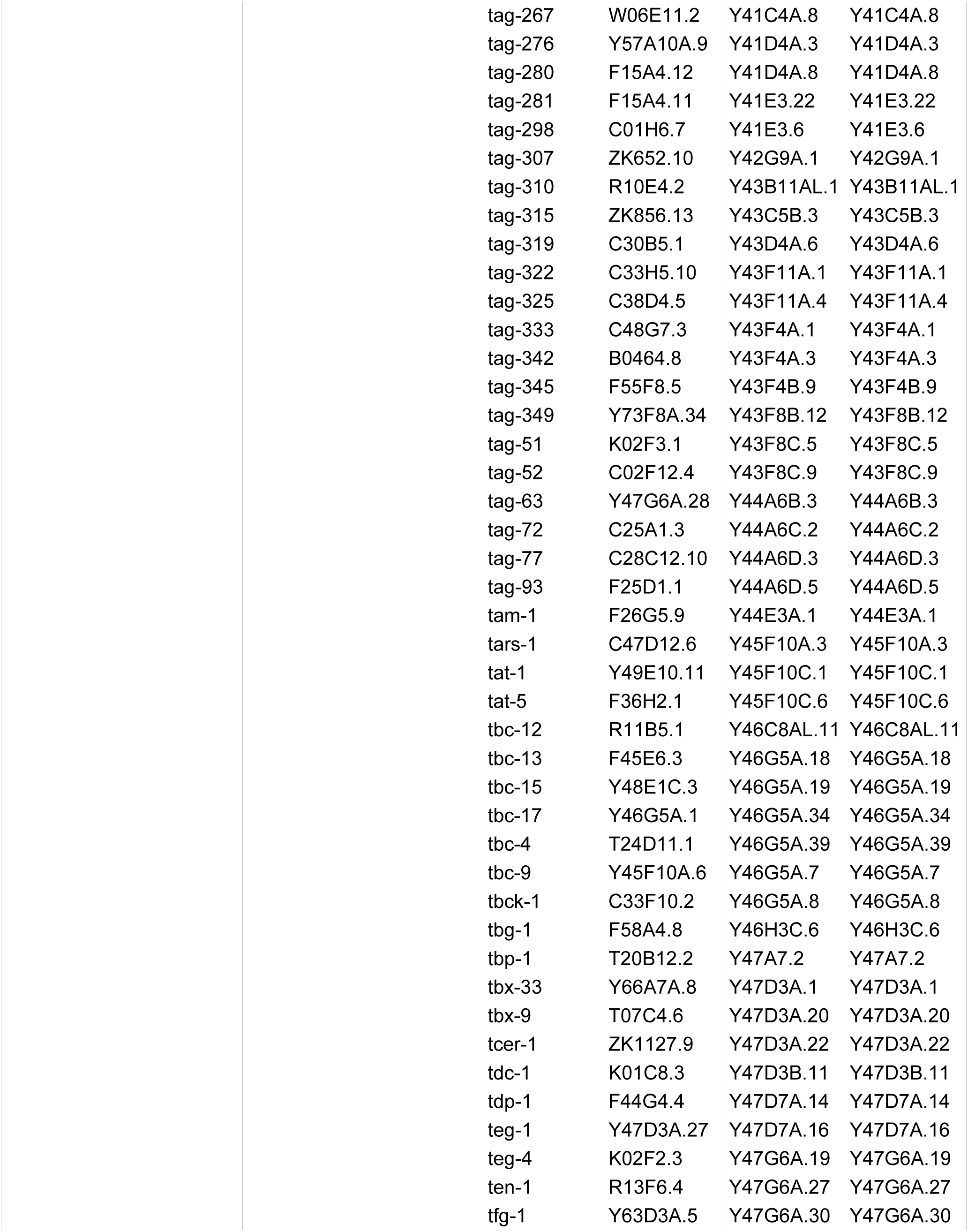
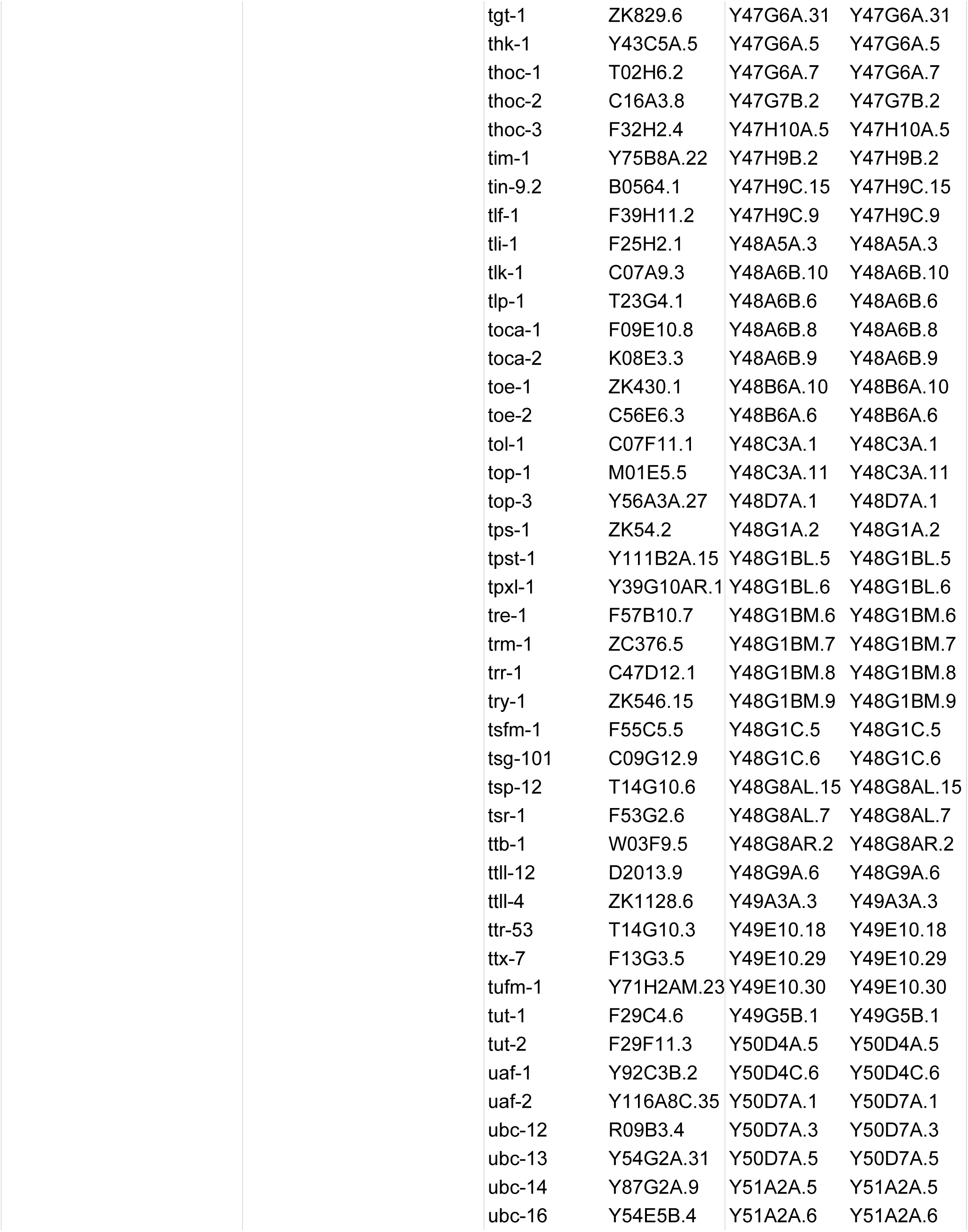
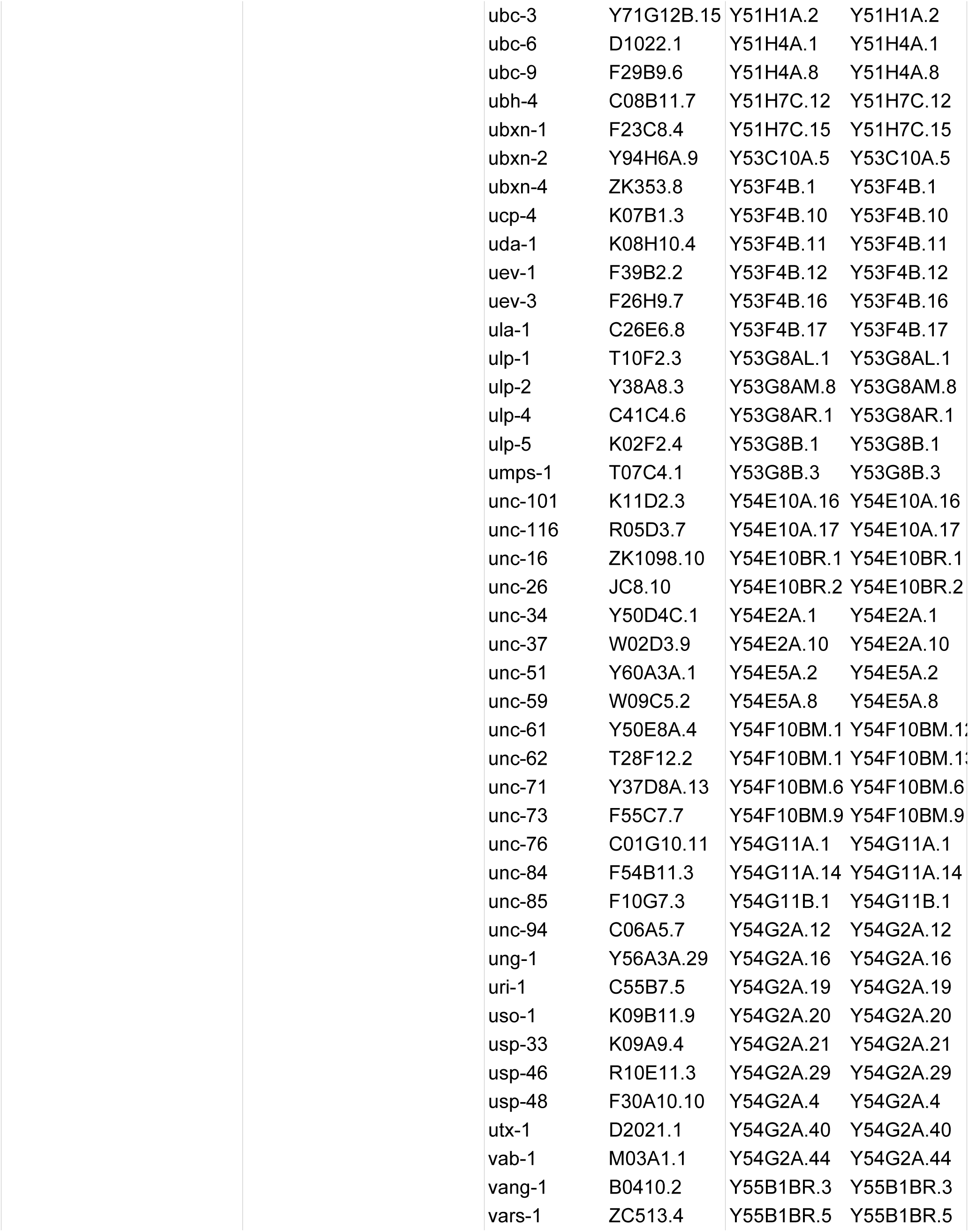
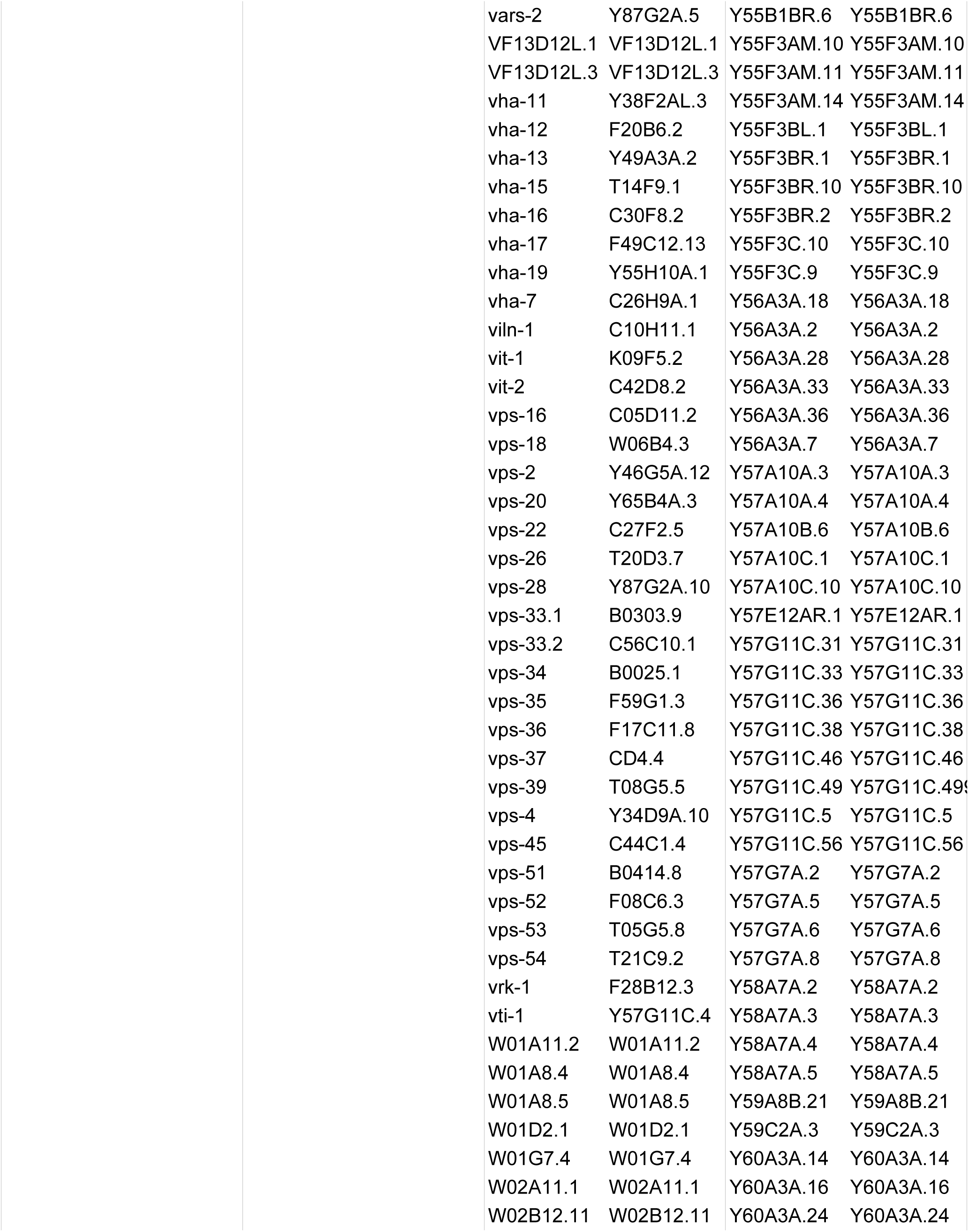
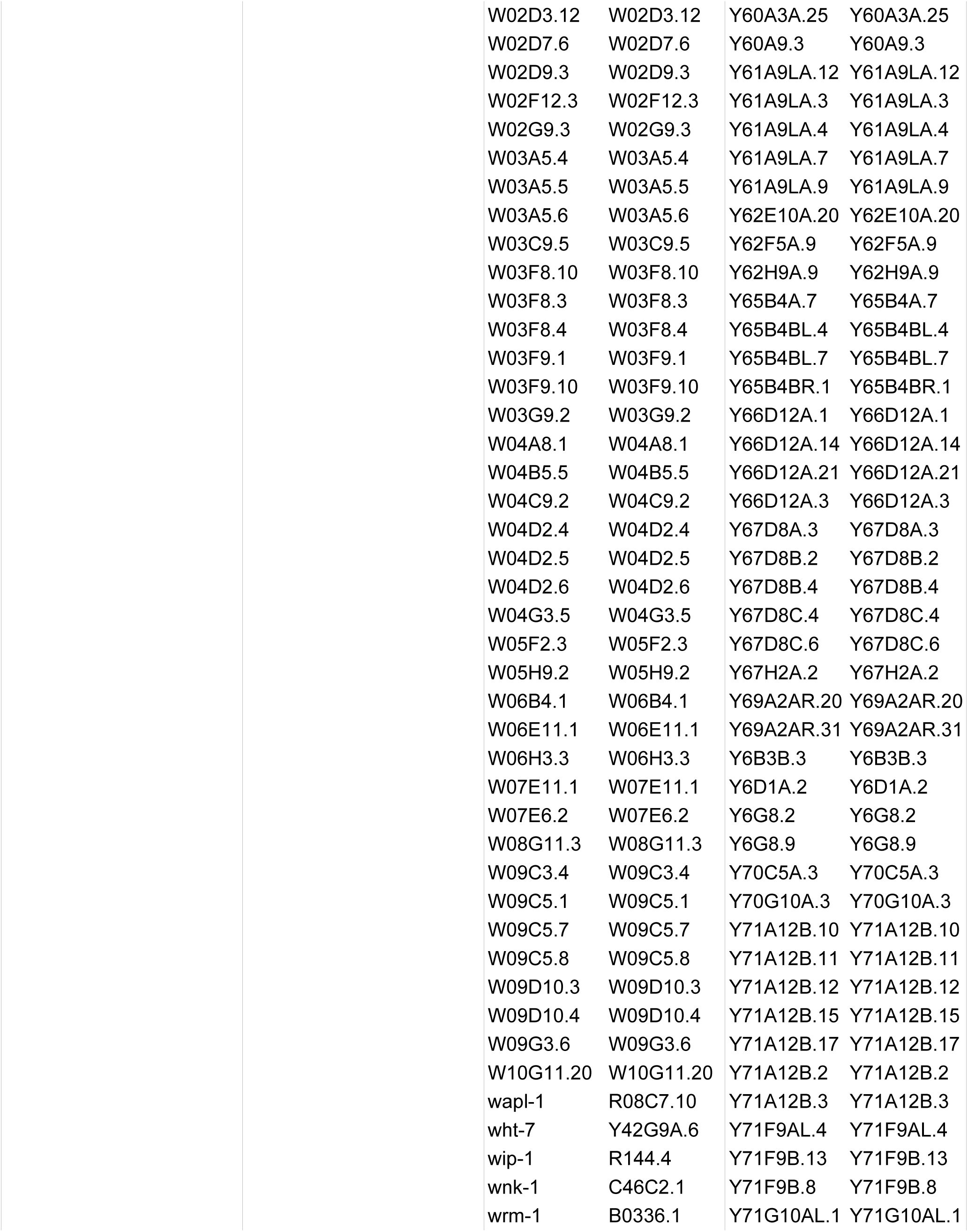
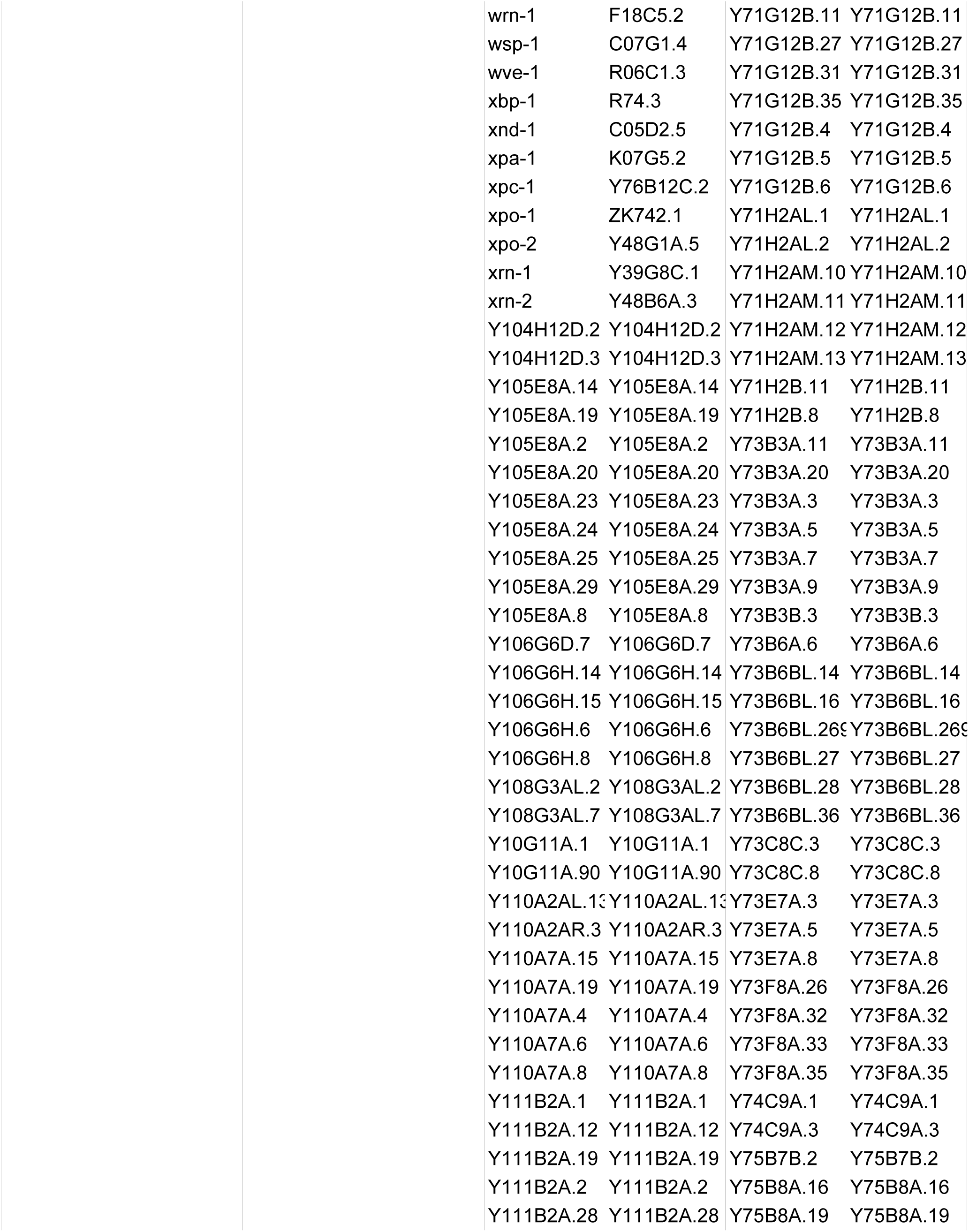
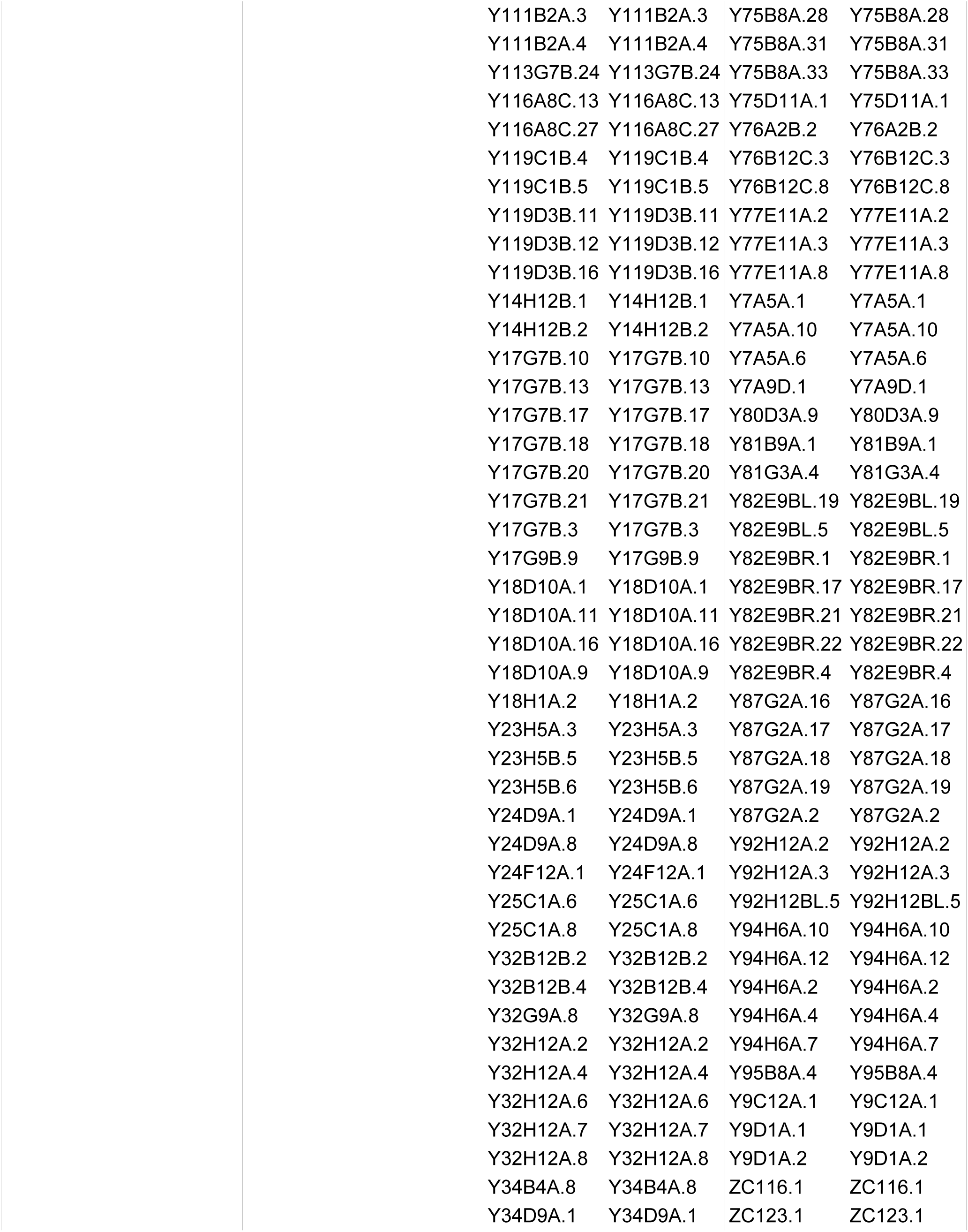
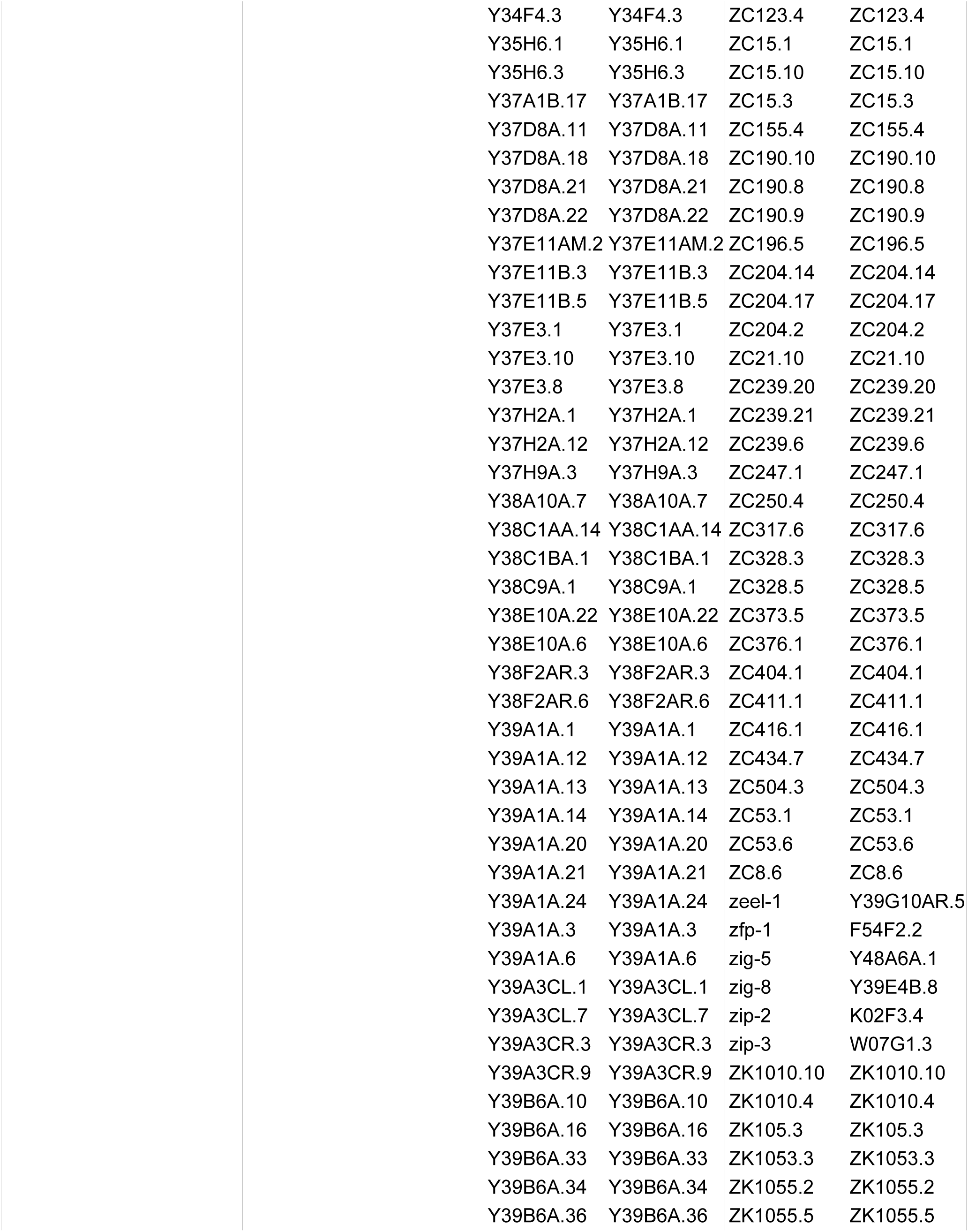
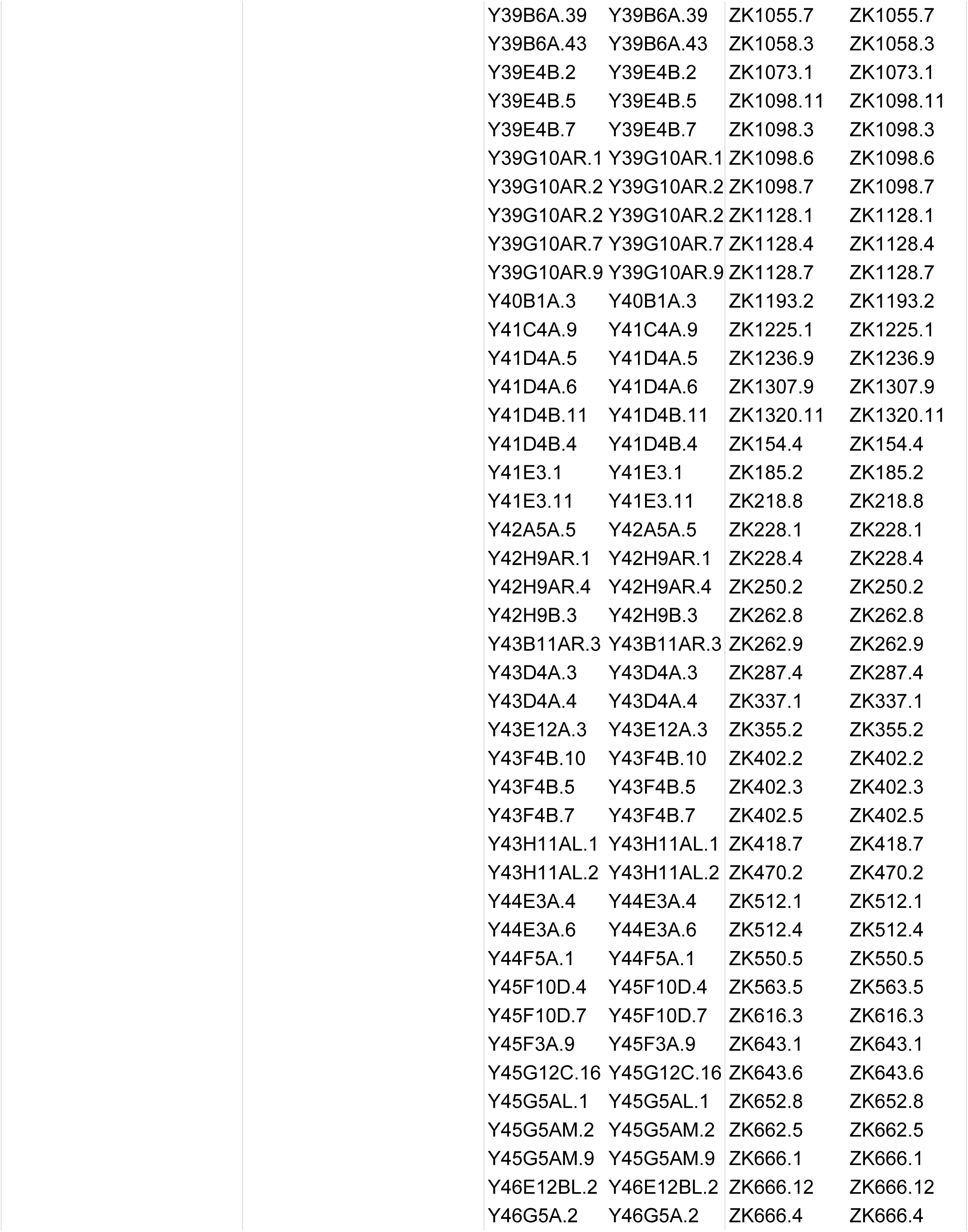
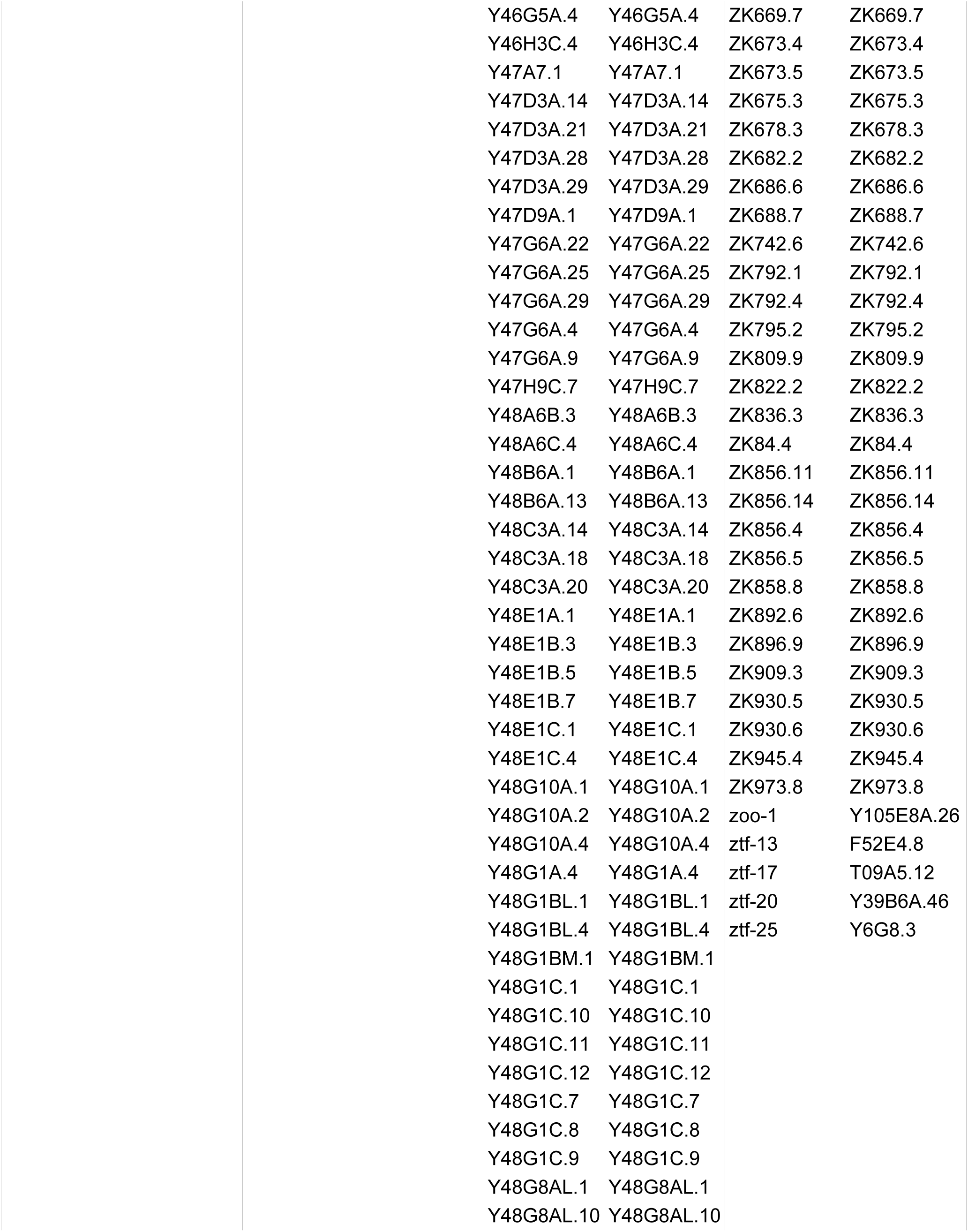
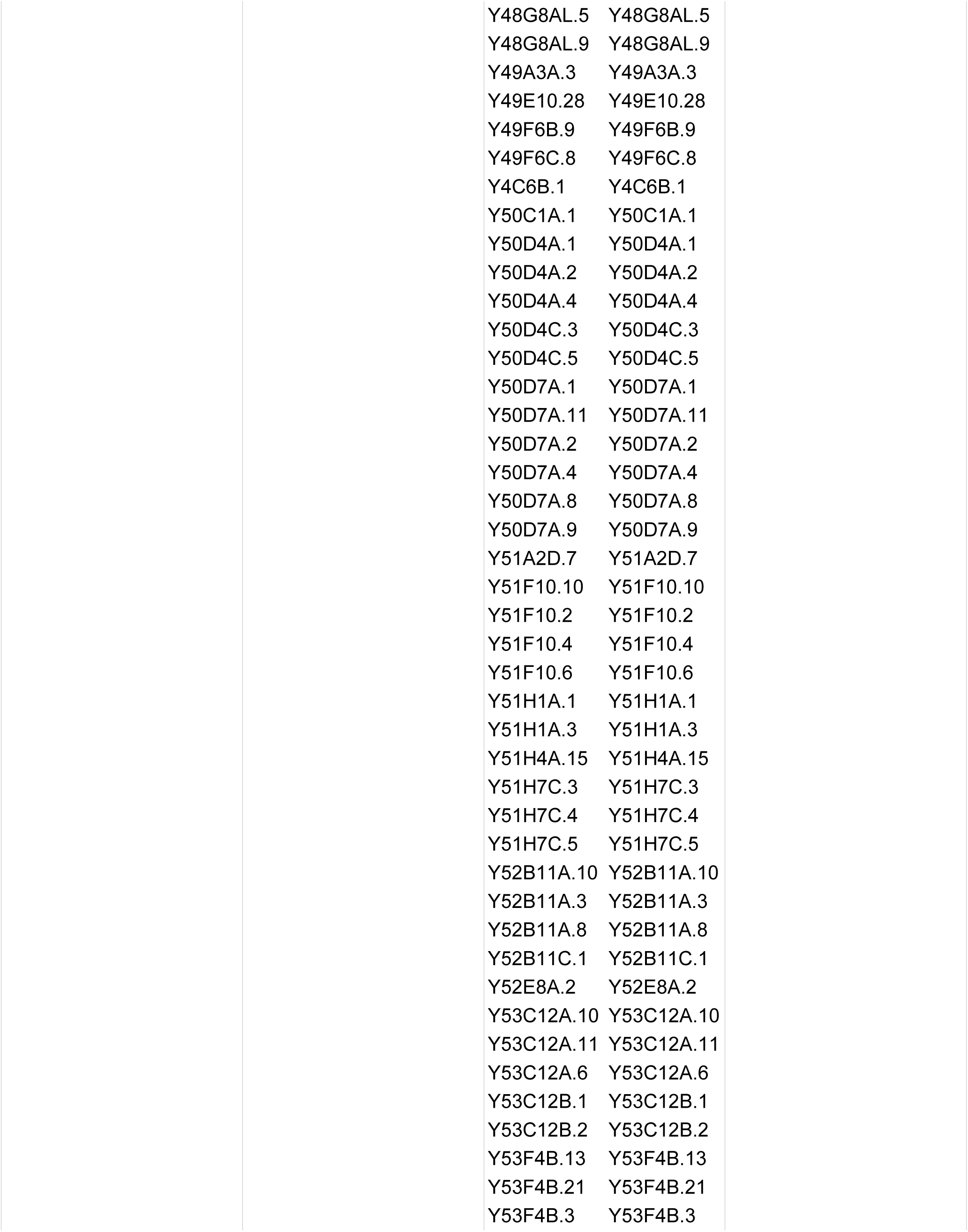
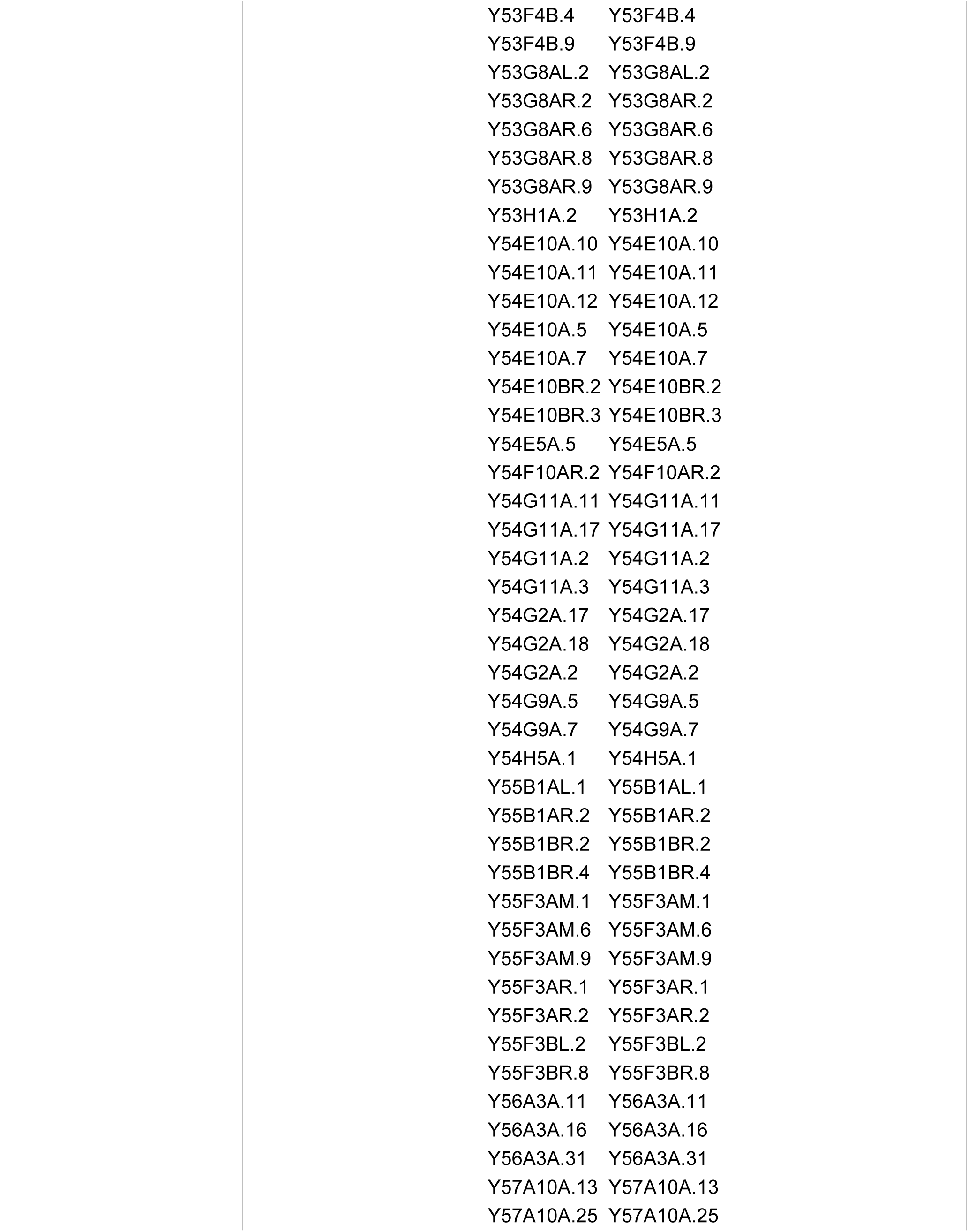
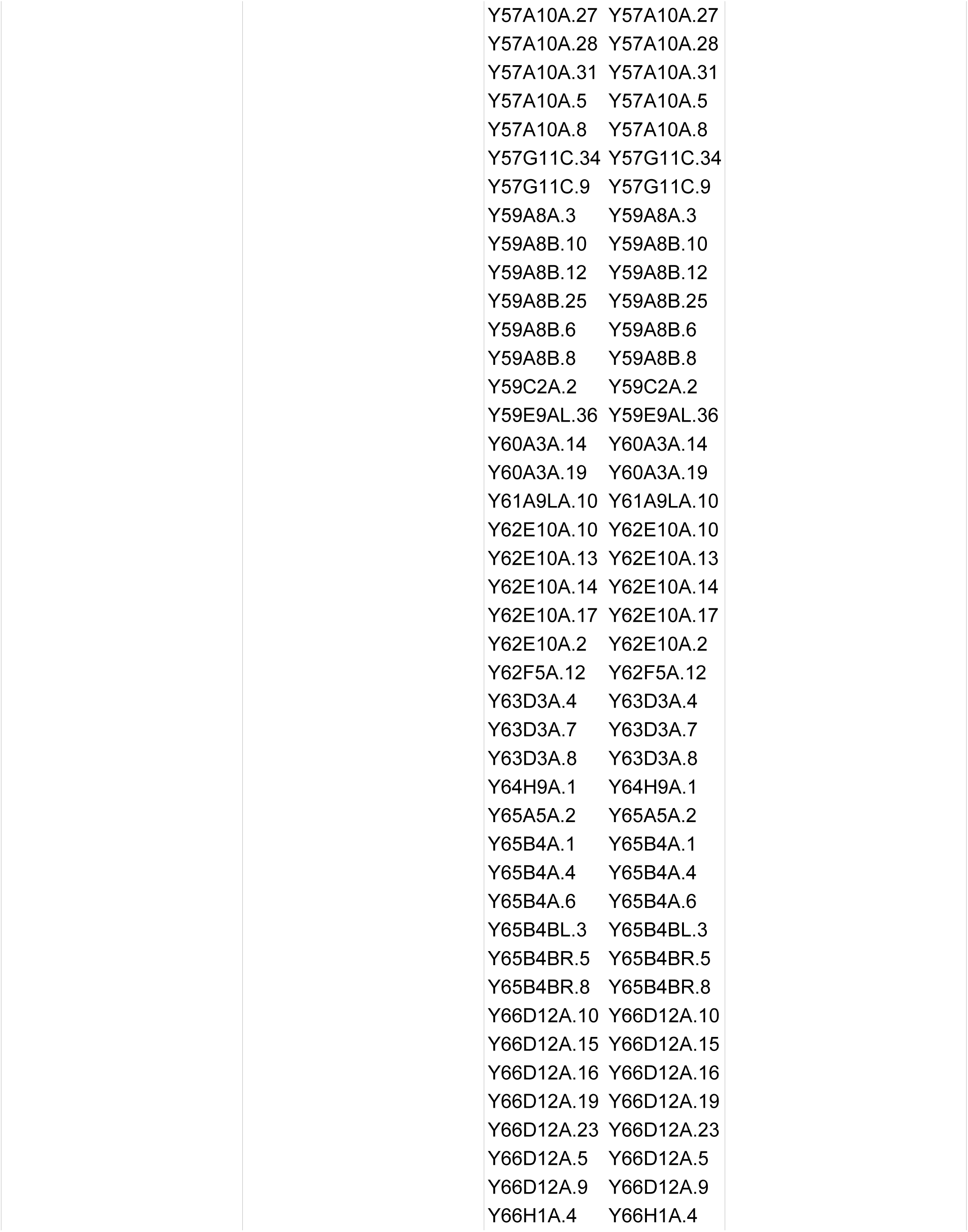
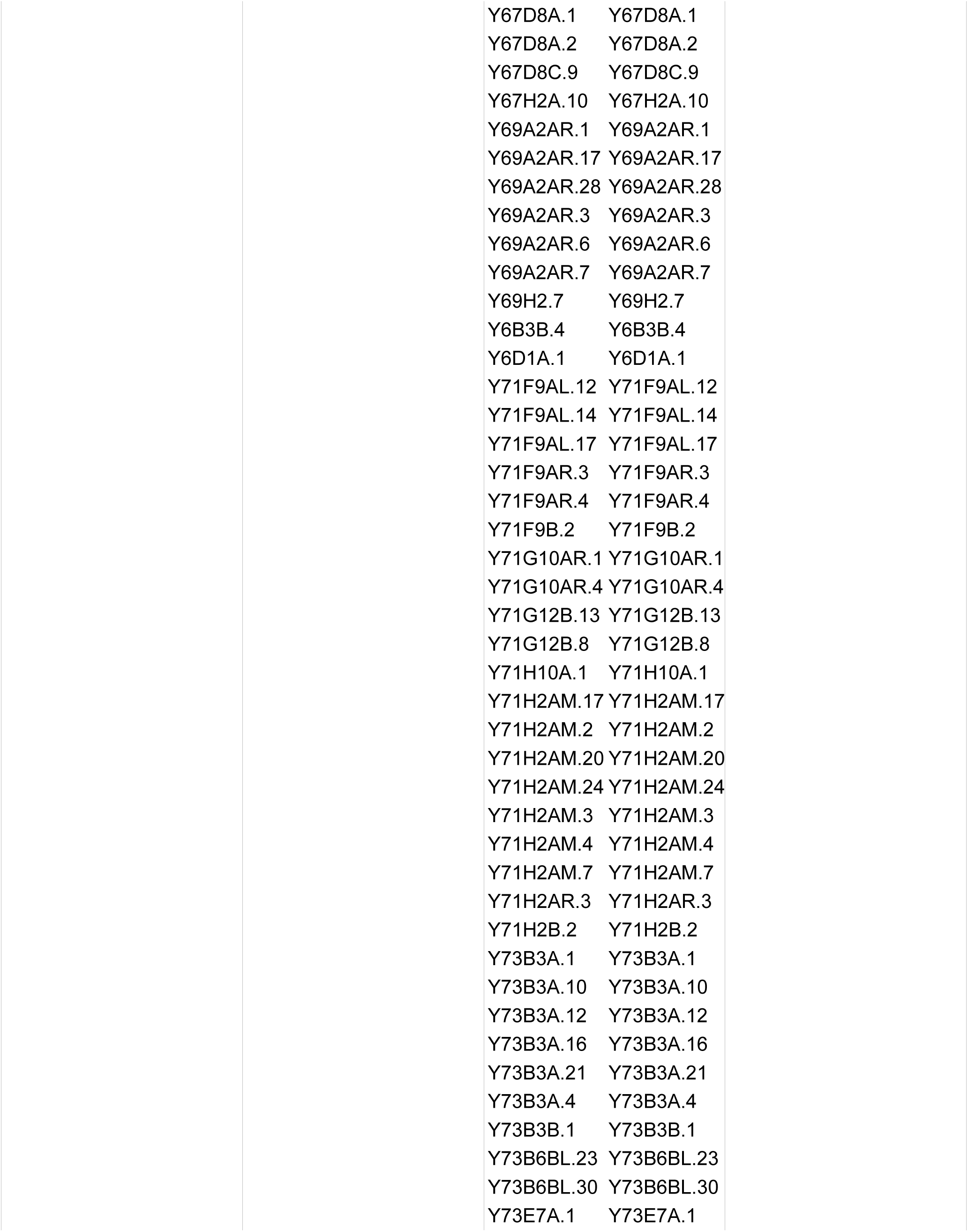
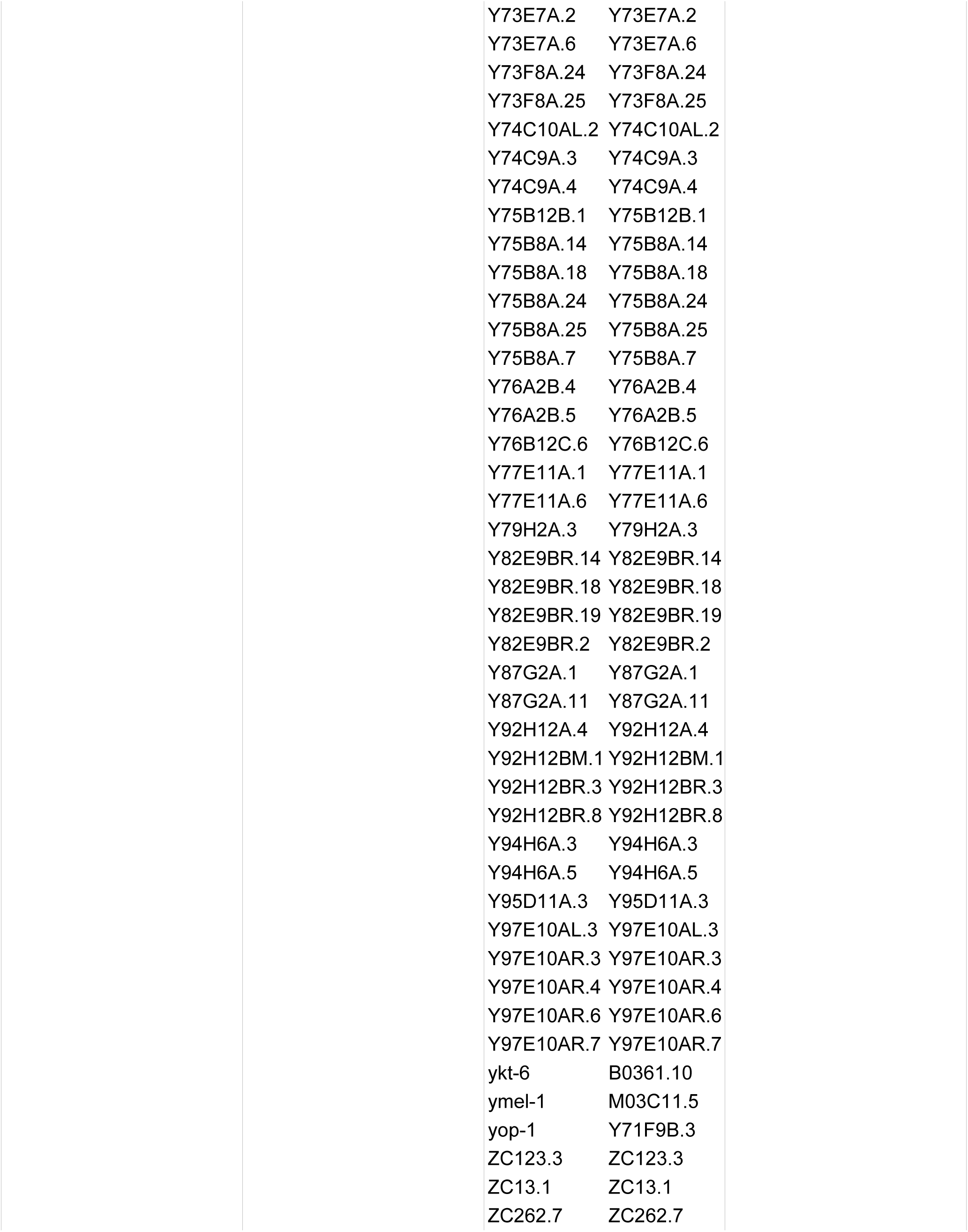
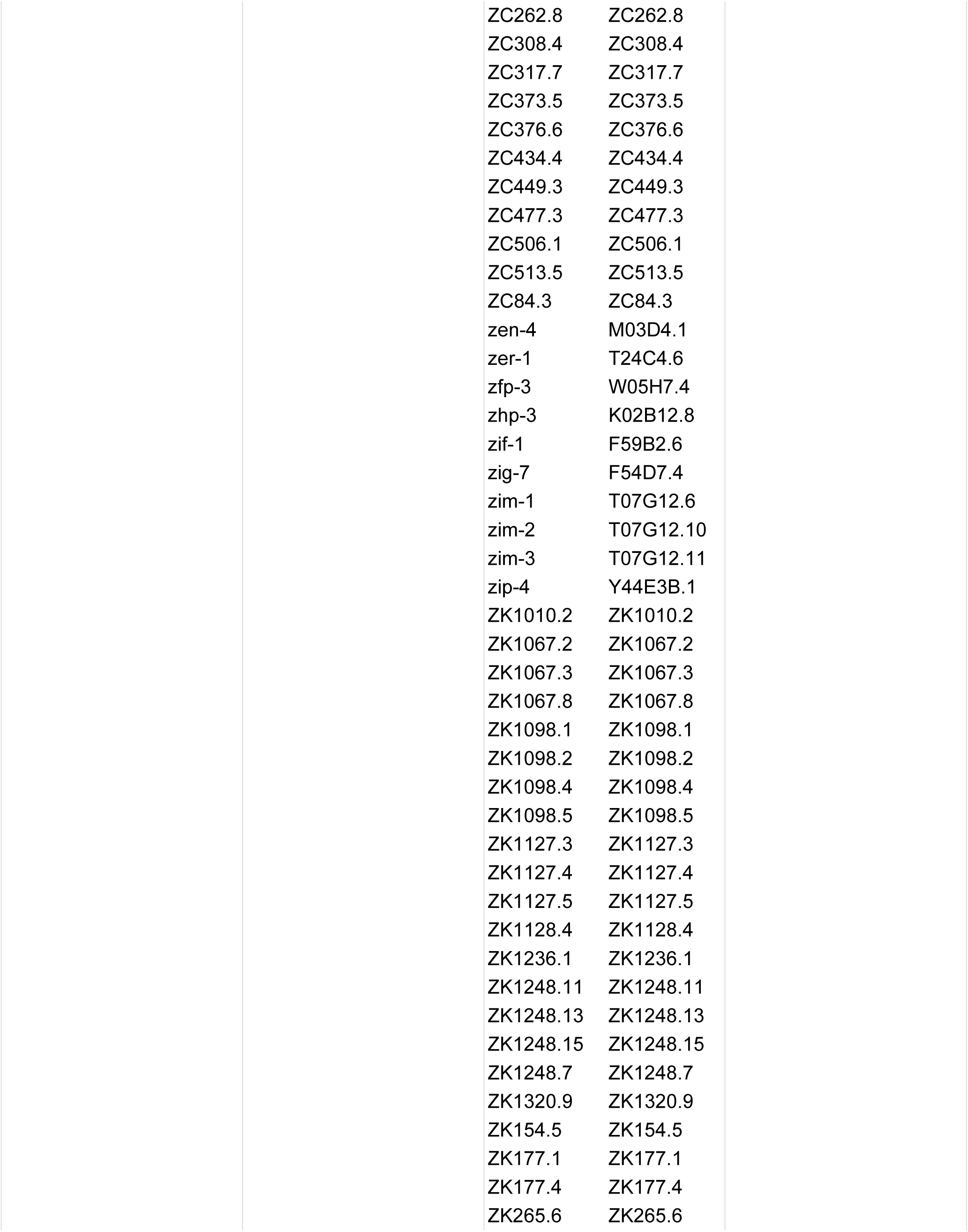
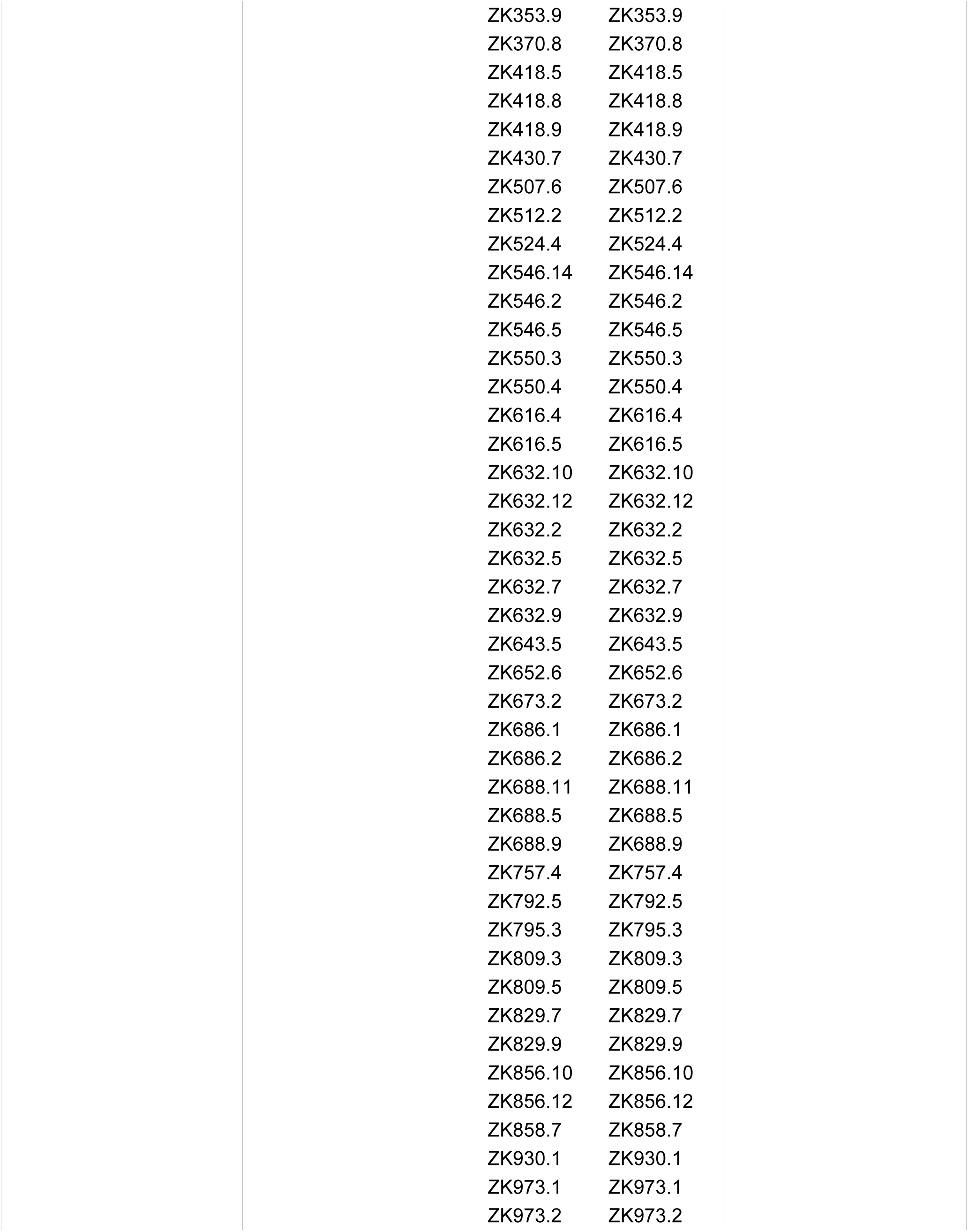
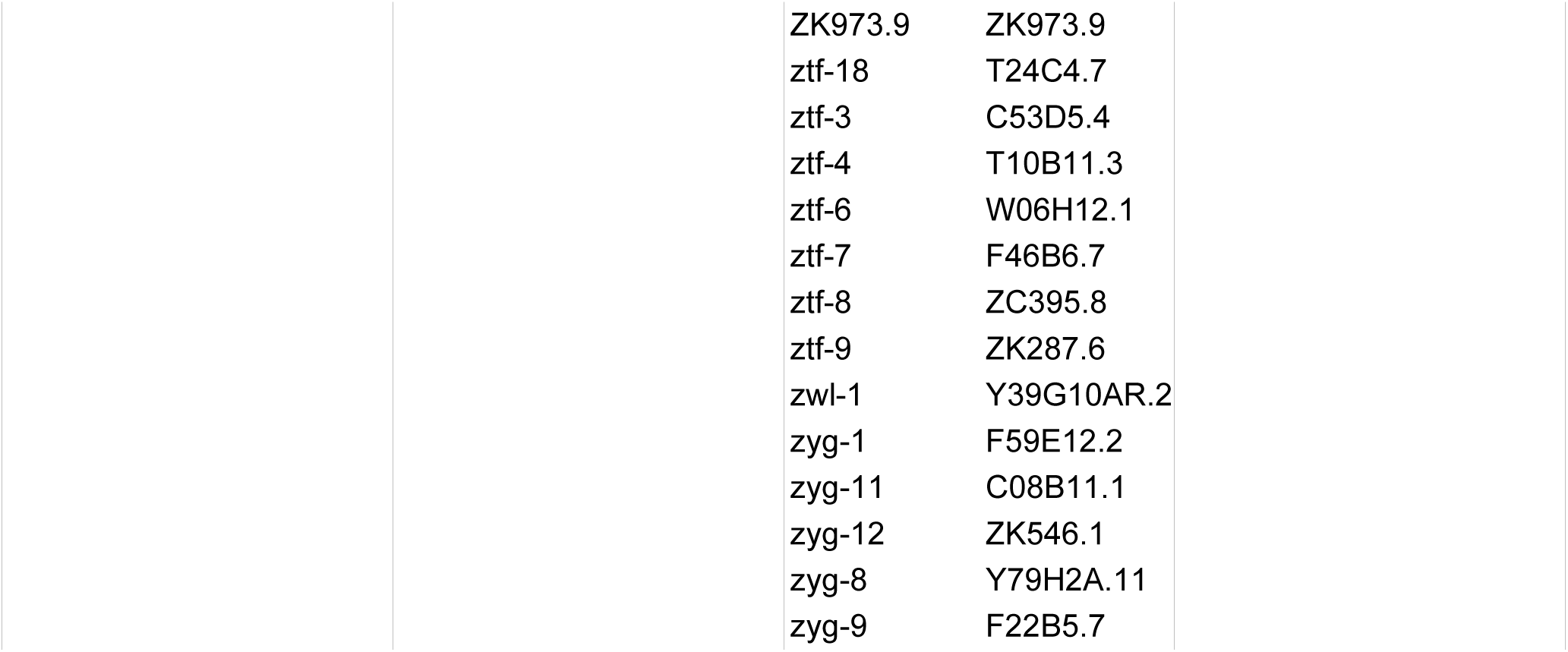

## REFERENCE

Ashe, A., Belicard, T., Le Pen, J., Sarkies, P., Frezal, L., Lehrbach, N.J., Felix, M.A., and Miska, E.A. (2013). A deletion polymorphism in the Caenorhabditis elegans RIG-I homolog disables viral RNA dicing and antiviral immunity. Elife 2, e00994.

Ashe, A., Sapetschnig, A., Weick, E.M., Mitchell, J., Bagijn, M.P., Cording, A.C., Doebley, A.L., Goldstein, L.D., Lehrbach, N.J., Le Pen, J., et al. (2012). piRNAs can trigger a multigenerational epigenetic memory in the germline of C. elegans. Cell 150, 88–99.

Bagijn, M.P., Goldstein, L.D., Sapetschnig, A., Weick, E.M., Bouasker, S., Lehrbach, N.J., Simard, M.J., and Miska, E.A. (2012). Function, Targets, and Evolution of Caenorhabditis elegans piRNAs. Science.

Baulcombe, D. (2004). RNA silencing in plants. Nature 431, 356–363.

Bernstein, E., Caudy, A.A., Hammond, S.M., and Hannon, G.J. (2001). Role for a bidentate ribonuclease in the initiation step of RNA interference. Nature 409, 363–366.

Bharadwaj, P.S., and Hall, S.E. (2017). Endogenous RNAi Pathways Are Required in Neurons for Dauer Formation in Caenorhabditis elegans. Genetics 205, 1503–1516.

Brenner, S. (1974). The genetics of Caenorhabditis elegans. Genetics 77, 71–94.

Cai, Q., Qiao, L., Wang, M., He, B., Lin, F.M., Palmquist, J., Huang, S.D., and Jin, H. (2018). Plants send small RNAs in extracellular vesicles to fungal pathogen to silence virulence genes. Science 360, 1126–1129.

Caplen, N.J., and Mousses, S. (2003). Short interfering RNA (siRNA)-mediated RNA interference (RNAi) in human cells. Ann N Y Acad Sci 1002, 56–62.

Claycomb, J.M., Batista, P.J., Pang, K.M., Gu, W., Vasale, J.J., van Wolfswinkel, J.C., Chaves, D.A., Shirayama, M., Mitani, S., Ketting, R.F., et al. (2009). The Argonaute CSR-1 and its 22G-RNA cofactors are required for holocentric chromosome segregation. Cell 139, 123–134.

Coffman, S.R., Lu, J., Guo, X., Zhong, J., Jiang, H., Broitman-Maduro, G., Li, W.X., Lu, R., Maduro, M., and Ding, S.W. (2017). Caenorhabditis elegans RIG-I Homolog Mediates Antiviral RNA Interference Downstream of Dicer-Dependent Biogenesis of Viral Small Interfering RNAs. mBio 8.

Conine, C.C., Batista, P.J., Gu, W., Claycomb, J.M., Chaves, D.A., Shirayama, M., and Mello, C.C. (2010). Argonautes ALG-3 and ALG-4 are required for spermatogenesis-specific 26G-RNAs and thermotolerant sperm in Caenorhabditis elegans. Proceedings of the National Academy of Sciences of the United States of America 107, 3588–3593.

Conine, C.C., Moresco, J.J., Gu, W., Shirayama, M., Conte, D., Yates, J.R., and Mello, C.C. (2013). Argonautes promote male fertility and provide a paternal memory of germline gene expression in C. elegans. Cell 155, 1532–1544.

Dent, J.A., Smith, M.M., Vassilatis, D.K., and Avery, L. (2000). The genetics of ivermectin resistance in Caenorhabditis elegans. Proc Natl Acad Sci U S A 97, 2674–2679.

Deshpande, T., Takagi, T., Hao, L., Buratowski, S., and Charbonneau, H. (1999). Human PIR1 of the protein-tyrosine phosphatase superfamily has RNA 5’-triphosphatase and diphosphatase activities. J Biol Chem 274, 16590–16594.

Duchaine, T.F., Wohlschlegel, J.A., Kennedy, S., Bei, Y., Conte, D., Pang, K., Brownell, D.R., Harding, S., Mitani, S., Ruvkun, G., et al. (2006). Functional proteomics reveals the biochemical niche of C. elegans DCR-1 in multiple small-RNA-mediated pathways. Cell 124, 343–354.

Fire, A., Xu, S., Montgomery, M.K., Kostas, S.A., Driver, S.E., and Mello, C.C. (1998). Potent and specific genetic interference by double-stranded RNA in Caenorhabditis elegans. Nature 391, 806–811.

Fischer, S.E., Montgomery, T.A., Zhang, C., Fahlgren, N., Breen, P.C., Hwang, A., Sullivan, C.M., Carrington, J.C., and Ruvkun, G. (2011). The ERI-6/7 helicase acts at the first stage of an siRNA amplification pathway that targets recent gene duplications. PLoS Genet 7, e1002369.

Gabel, H.W., and Ruvkun, G. (2008). The exonuclease ERI-1 has a conserved dual role in 5.8S rRNA processing and RNAi. Nat Struct Mol Biol 15, 531–533.

Gent, J.I., Lamm, A.T., Pavelec, D.M., Maniar, J.M., Parameswaran, P., Tao, L., Kennedy, S., and Fire, A.Z. (2010). Distinct phases of siRNA synthesis in an endogenous RNAi pathway in C. elegans soma. Mol Cell 37, 679–689.

Gerson-Gurwitz, A., Wang, S., Sathe, S., Green, R., Yeo, G.W., Oegema, K., and Desai, A. (2016). A Small RNA-Catalytic Argonaute Pathway Tunes Germline Transcript Levels to Ensure Embryonic Divisions. Cell 165, 396–409.

Grentzinger, T., Armenise, C., Brun, C., Mugat, B., Serrano, V., Pelisson, A., and Chambeyron, S. (2012). piRNA-mediated transgenerational inheritance of an acquired trait. Genome research.

Grishok, A., Pasquinelli, A.E., Conte, D., Li, N., Parrish, S., Ha, I., Baillie, D.L., Fire, A., Ruvkun, G., and Mello, C.C. (2001). Genes and mechanisms related to RNA interference regulate expression of the small temporal RNAs that control C. elegans developmental timing. Cell 106, 23–34.

Gu, W., Claycomb, J.M., Batista, P.J., Mello, C.C., and Conte, D. (2011). Cloning Argonaute-associated small RNAs from Caenorhabditis elegans. Methods in Molecular Biology (Clifton, NJ) 725, 251–280.

Gu, W., Lee, H.-C., Chaves, D., Youngman, E.M., Pazour, G.J., Conte, D., and Mello, C.C. (2012). CapSeq and CIP-TAP identify Pol II start sites and reveal capped small RNAs as C. elegans piRNA precursors. Cell 151, 1488–1500.

Gu, W., Shirayama, M., Conte, D., Vasale, J., Batista, P.J., Claycomb, J.M., Moresco, J.J., Youngman, E.M., Keys, J., Stoltz, M.J., et al. (2009). Distinct argonaute-mediated 22G-RNA pathways direct genome surveillance in the C. elegans germline. Mol Cell 36, 231–244.

Guo, X., Zhang, R., Wang, J., Ding, S.W., and Lu, R. (2013). Homologous RIG-I-like helicase proteins direct RNAi-mediated antiviral immunity in C. elegans by distinct mechanisms. Proc Natl Acad Sci U S A 110, 16085–16090.

Han, T., Manoharan, A.P., Harkins, T.T., Bouffard, P., Fitzpatrick, C., Chu, D.S., Thierry-Mieg, D., Thierry-Mieg, J., and Kim, J.K. (2009). 26G endo-siRNAs regulate spermatogenic and zygotic gene expression in Caenorhabditis elegans. Proc Natl Acad Sci U S A 106, 18674–18679.

Hannon, G.J. (2002). RNA interference. Nature 418, 244–251.

Hong, Y., Lee, R.C., and Ambros, V. (2000). Structure and function analysis of LIN-14, a temporal regulator of postembryonic developmental events in Caenorhabditis elegans. Mol Cell Biol 20, 2285–2295.

Hornung, V., Ellegast, J., Kim, S., Brzozka, K., Jung, A., Kato, H., Poeck, H., Akira, S., Conzelmann, K.K., Schlee, M., et al. (2006). 5’-Triphosphate RNA is the ligand for RIG-I. Science 314, 994–997.

Hou, Y., Zhai, Y., Feng, L., Karimi, H.Z., Rutter, B.D., Zeng, L., Choi, D.S., Zhang, B., Gu, W., Chen, X., et al. (2019). A Phytophthora Effector Suppresses Trans-Kingdom RNAi to Promote Disease Susceptibility. Cell Host Microbe 25, 153–165 e155.

Kato, H., Takeuchi, O., Sato, S., Yoneyama, M., Yamamoto, M., Matsui, K., Uematsu, S., Jung, A., Kawai, T., Ishii, K.J., et al. (2006). Differential roles of MDA5 and RIG-I helicases in the recognition of RNA viruses. Nature 441, 101–105.

Katsuma, S., Koyano, Y., Kang, W., Kokusho, R., Kamita, S.G., and Shimada, T. (2012). The baculovirus uses a captured host phosphatase to induce enhanced locomotory activity in host caterpillars. PLoS Pathog 8, e1002644.

Kennedy, S., Wang, D., and Ruvkun, G. (2004). A conserved siRNA-degrading RNase negatively regulates RNA interference in C. elegans. Nature 427, 645–649.

Langmead, B., Trapnell, C., Pop, M., and Salzberg, S.L. (2009). Ultrafast and memory-efficient alignment of short DNA sequences to the human genome. Genome Biology 10, R25.

Lee, H.C., Gu, W., Shirayama, M., Youngman, E., Conte, D., and Mello, C.C. (2012). C. elegans piRNAs mediate the genome-wide surveillance of germline transcripts. Cell 150, 78–87.

Li, L., Dai, H., Nguyen, A.P., and Gu, W. (2019). A convenient strategy to clone modified/unmodified small RNA and mRNA for high throughput sequencing. RNA 26, 218–227.

Macrae, I.J., Zhou, K., Li, F., Repic, A., Brooks, A.N., Cande, W.Z., Adams, P.D., and Doudna, J.A. (2006). Structural basis for double-stranded RNA processing by Dicer. Science 311, 195–198.

McCaffrey, A.P., Meuse, L., Pham, T.T., Conklin, D.S., Hannon, G.J., and Kay, M.A. (2002). RNA interference in adult mice. Nature 418, 38–39.

Pak, J., and Fire, A. (2007). Distinct populations of primary and secondary effectors during RNAi in C. elegans. Science (New York, NY) 315, 241–244.

Pal-Bhadra, M., Bhadra, U., and Birchler, J.A. (2002). RNAi related mechanisms affect both transcriptional and posttranscriptional transgene silencing in Drosophila. Mol Cell 9, 315–327.

Parrish, S., and Fire, A. (2001). Distinct roles for RDE-1 and RDE-4 during RNA interference in Caenorhabditis elegans. RNA 7, 1397–1402.

Pavelec, D.M., Lachowiec, J., Duchaine, T.F., Smith, H.E., and Kennedy, S. (2009). Requirement for the ERI/DICER complex in endogenous RNA interference and sperm development in Caenorhabditis elegans. Genetics 183, 1283–1295.

Posner, R., Toker, I.A., Antonova, O., Star, E., Anava, S., Azmon, E., Hendricks, M., Bracha, S., Gingold, H., and Rechavi, O. (2019). Neuronal Small RNAs Control Behavior Transgenerationally. Cell 177, 1814–1826 e1815.

Ruby, J.G., Jan, C., Player, C., Axtell, M.J., Lee, W., Nusbaum, C., Ge, H., and Bartel, D.P. (2006). Large-scale sequencing reveals 21U-RNAs and additional microRNAs and endogenous siRNAs in C. elegans. Cell 127, 1193–1207.

Sankhala, R.S., Lokareddy, R.K., and Cingolani, G. (2014). Structure of human PIR1, an atypical dual-specificity phosphatase. Biochemistry 53, 862–871.

Schirmer, E.C., Yates, J.R., 3rd, and Gerace, L. (2003). MudPIT: A powerful proteomics tool for discovery. Discov Med 3, 38–39.

Seth, M., Shirayama, M., Gu, W., Ishidate, T., Conte, D., and Mello, C.C. (2013). The C. elegans CSR-1 argonaute pathway counteracts epigenetic silencing to promote germline gene expression. Dev Cell 27, 656–663.

Shatkin, A.J. (1976). Capping of eucaryotic mRNAs. Cell 9, 645–653.

Shen, E.Z., Chen, H., Ozturk, A.R., Tu, S., Shirayama, M., Tang, W., Ding, Y.H., Dai, S.Y., Weng, Z., and Mello, C.C. (2018). Identification of piRNA Binding Sites Reveals the Argonaute Regulatory Landscape of the C. elegans Germline. Cell 172, 937–951 e918.

Shirayama, M., Seth, M., Lee, H.C., Gu, W., Ishidate, T., Conte, D., and Mello, C.C. (2012). piRNAs initiate an epigenetic memory of nonself RNA in the C. elegans germline. Cell 150, 65–77.

Simmer, F., Moorman, C., van der Linden, A.M., Kuijk, E., van den Berghe, P.V., Kamath, R.S., Fraser, A.G., Ahringer, J., and Plasterk, R.H. (2003). Genome-wide RNAi of C. elegans using the hypersensitive rrf-3 strain reveals novel gene functions. PLoS Biol 1, E12.

Simmer, F., Tijsterman, M., Parrish, S., Koushika, S.P., Nonet, M.L., Fire, A., Ahringer, J., and Plasterk, R.H. (2002). Loss of the putative RNA-directed RNA polymerase RRF-3 makes C. elegans hypersensitive to RNAi. Curr Biol 12, 1317–1319.

Stein, L.D., Mungall, C., Shu, S., Caudy, M., Mangone, M., Day, A., Nickerson, E., Stajich, J.E., Harris, T.W., Arva, A., et al. (2002). The generic genome browser: a building block for a model organism system database. Genome Research 12, 1599–1610.

Steiner, F.A., Okihara, K.L., Hoogstrate, S.W., Sijen, T., and Ketting, R.F. (2009). RDE-1 slicer activity is required only for passenger-strand cleavage during RNAi in Caenorhabditis elegans. Nat Struct Mol Biol 16, 207–211.

Tabara, H., Sarkissian, M., Kelly, W.G., Fleenor, J., Grishok, A., Timmons, L., Fire, A., and Mello, C.C. (1999). The rde-1 gene, RNA interference, and transposon silencing in C. elegans. Cell 99, 123–132.

Tabara, H., Yigit, E., Siomi, H., and Mello, C.C. (2002). The dsRNA binding protein RDE-4 interacts with RDE-1, DCR-1, and a DExH-box helicase to direct RNAi in C. elegans. Cell 109, 861–871.

Takagi, T., Taylor, G.S., Kusakabe, T., Charbonneau, H., and Buratowski, S. (1998). A protein tyrosine phosphatase-like protein from baculovirus has RNA 5’-triphosphatase and diphosphatase activities. Proc Natl Acad Sci U S A 95, 9808–9812.

Thivierge, C., Makil, N., Flamand, M., Vasale, J.J., Mello, C.C., Wohlschlegel, J., Conte, D., Jr., and Duchaine, T.F. (2011). Tudor domain ERI-5 tethers an RNA-dependent RNA polymerase to DCR-1 to potentiate endo-RNAi. Nat Struct Mol Biol 19, 90–97.

Timmons, L. (2004). Endogenous inhibitors of RNA interference in Caenorhabditis elegans. Bioessays 26, 715–718.

Vasale, J.J., Gu, W., Thivierge, C., Batista, P.J., Claycomb, J.M., Youngman, E.M., Duchaine, T.F., Mello, C.C., and Conte, D. (2010). Sequential rounds of RNA-dependent RNA transcription drive endogenous small-RNA biogenesis in the ERGO-1/Argonaute pathway. Proc Natl Acad Sci U S A 107, 3582–3587.

Welker, N.C., Maity, T.S., Ye, X., Aruscavage, P.J., Krauchuk, A.A., Liu, Q., and Bass, B.L. (2011). Dicer’s helicase domain discriminates dsRNA termini to promote an altered reaction mode. Mol Cell 41, 589–599.

Welker, N.C., Pavelec, D.M., Nix, D.A., Duchaine, T.F., Kennedy, S., and Bass, B.L. (2010). Dicer’s helicase domain is required for accumulation of some, but not all, C. elegans endogenous siRNAs. RNA 16, 893–903.

Yigit, E., Batista, P.J., Bei, Y., Pang, K.M., Chen, C.C., Tolia, N.H., Joshua-Tor, L., Mitani, S., Simard, M.J., and Mello, C.C. (2006). Analysis of the C. elegans Argonaute family reveals that distinct Argonautes act sequentially during RNAi. Cell 127, 747–757.

Yuan, Y., Li, D.M., and Sun, H. (1998). PIR1, a novel phosphatase that exhibits high affinity to RNA ribonucleoprotein complexes. J Biol Chem 273, 20347–20353.

Zhang, C., Montgomery, T.A., Gabel, H.W., Fischer, S.E., Phillips, C.M., Fahlgren, N., Sullivan, C.M., Carrington, J.C., and Ruvkun, G. (2011). mut-16 and other mutator class genes modulate 22G and 26G siRNA pathways in Caenorhabditis elegans. Proc Natl Acad Sci U S A 108, 1201–1208.

Zhang, D., Tu, S., Stubna, M., Wu, W.S., Huang, W.C., Weng, Z., and Lee, H.C. (2018). The piRNA targeting rules and the resistance to piRNA silencing in endogenous genes. Science 359, 587–592.

Zhang, H., Kolb, F.A., Brondani, V., Billy, E., and Filipowicz, W. (2002). Human Dicer preferentially cleaves dsRNAs at their termini without a requirement for ATP. EMBO J 21, 5875–5885.

## SUPPLEMENTARY REFERENCE

Schürer, H., Lang, K., Schuster J., Mörl M. (2002). A Universal Method to Produce in Vitro Transcripts with Homogeneous 3’ Ends. Nucleic Acids Res 30, e56

Phillips, C.M., McDonald, K.L., and Dernburg, A.F. (2009). Cytological analysis of meiosis in Caenorhabditis elegans. Methods Mol Biol 558, 171–195.

